# Regionalized regulation of actomyosin organization influences cardiomyocyte cell shape changes during chamber curvature formation

**DOI:** 10.1101/2025.01.07.631779

**Authors:** Dena M. Leerberg, Gabriel B. Avillion, Rashmi Priya, Didier Y.R. Stainier, Deborah Yelon

**Author notes:** Correspondence to Deborah Yelon.

## Abstract

Cardiac chambers emerge from a heart tube that balloons and bends to create expanded ventricular and atrial structures, each containing a convex outer curvature (OC) and a recessed inner curvature (IC). The cellular and molecular mechanisms underlying the formation of these characteristic curvatures remain poorly understood. Here, we demonstrate in zebrafish that the initially similar populations of OC and IC ventricular cardiomyocytes diverge in the organization of their actomyosin cytoskeleton and subsequently acquire distinct OC and IC cell shapes. Altering actomyosin dynamics hinders cell shape changes in the OC, and mosaic analyses indicate that actomyosin regulates cardiomyocyte shape in a cell-autonomous manner. Additionally, both biomechanical cues and the transcription factor Tbx5a influence the basal enrichment of actomyosin and squamous cell morphologies in the OC. Together, our findings demonstrate that intrinsic and extrinsic factors intersect to control actomyosin organization in OC cardiomyocytes, which in turn promotes the cell shape changes that accompany curvature morphogenesis.

## Introduction

Organs are replete with curvature, from the branching distal tips of the lung epithelium to the concave cup of the retina, and to the gyri and sulci of the brain. Essentially every organ exhibits some degree of curvature during its morphogenesis, and the precise curvature geometry required for a particular organ’s function is likely sculpted from inputs both extrinsic and intrinsic to its tissues (Schamberger et al., 2023). Decades of interest in this topic show us that some curvatures rely heavily on external influences. For example, the vertebrate gut acquires its dramatic twists and turns due to its rapid lengthening combined with its restrictive attachment to a relatively static partner, the dorsal mesentery (Savin et al., 2011). By contrast, the ingressing *Drosophila* ventral furrow is controlled largely by tissue-intrinsic processes, with apical constriction driven by actomyosin pulses in the epithelial cells resulting in robust invagination (Leptin & Grunewald, 1990; Martin et al., 2010; Sweeton et al., 1991). Here, we investigate the factors, both tissue-extrinsic and -intrinsic, that control the regionalized cell shape changes that accompany chamber curvature formation in the developing heart.

The heart begins as a simple tube but then transforms into a looped organ with ventricular and atrial chambers. This looping process involves coordinated bending and ballooning of myocardial tissue to form the chambers, the twisting of the heart into a slightly helical shape (known as torsion), and the asymmetric displacement of the ventricle and atrium to deliver the chambers to their ultimate positions (Ivanovitch et al., 2017; Männer, 2000, 2004, 2009; Tessadori et al., 2021). During formation of the chambers, each acquires a bulging outer curvature (OC) and a recessed inner curvature (IC) (Figure 1C) (Jensen et al., 2013; Moorman & Christoffels, 2003; Taber, 2006). In addition to distinctive tissue morphology, these curvatures exhibit disparate physiological features; for example, the OC exhibits higher conduction velocity (Chi et al., 2008; Panáková et al., 2010; Weber et al., 2017) and more trabeculation (Liu et al., 2010; Peshkovsky et al., 2011; Rana et al., 2007) than the IC, whereas the IC demonstrates greater tissue stiffness (Zamir et al., 2003). While the differences between the OC and IC have been well characterized, the mechanisms that initially identify these regions and control their subsequent morphogenesis and specialization remain enigmatic.

**Figure 1.**
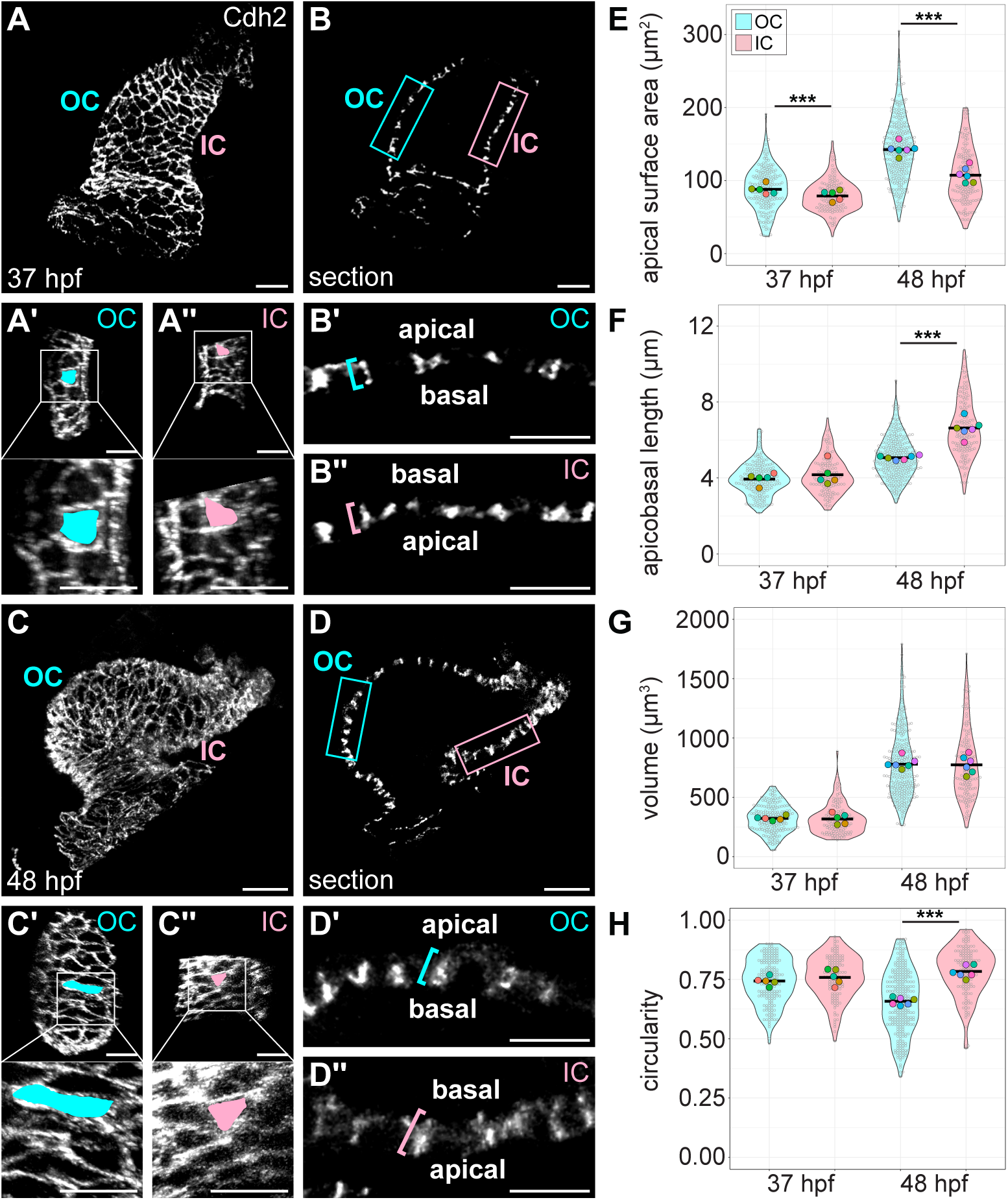
OC and IC cardiomyocyte morphologies diverge during curvature formation. (**A and C**) 3D reconstructions of 37 hpf (A) and 48 hpf (C) wild-type hearts. Immunostaining for Cdh2 labels lateral membranes of cardiomyocytes (see Figure 1—figure supplement 2 and Materials and Methods for more detail regarding use of Cdh2). OC (Aʹ and Cʹ) and IC (Aʹʹ and Cʹʹ) are shown for hearts in (A) and (C), respectively. Insets show higher magnification. Apical surface area of an individual cardiomyocyte is illustrated by blue or pink fill. (**B and D**) Sections through hearts in (A and C). (Bʹ, Bʹʹ, Dʹ, and Dʹʹ) are magnified views of blue (OC) and pink (IC) boxed regions in (B and D); blue and pink brackets highlight apicobasal length of individual cardiomyocytes. (**E-H**) Violin plots compare apical surface area, apicobasal length, volume, or circularity of OC and IC cardiomyocytes at 37 and 48 hpf. Each small grey dot represents an individual cell, each black bar represents the mean of values from individual cells, and each large colored dot represents the mean of all values from an individual embryo. Volume is calculated as LxWxH; circularity is calculated as 4⫪(A/P^2^). *** denotes p < 0.001, Wilcoxon test. Significance only shown for OC/IC comparisons; all metrics are significantly different between developmental stages of the same region (*i.e.* 37 hpf OC vs 48 hpf OC, and 37 hpf IC vs 48 hpf IC). 37 hpf OC (N=5 embryos, n=183 cells); 37 hpf IC (N=5 embryos, n=127 cells); 48 hpf OC (N=6 embryos, n=281 cells); 48 hpf IC (N=6 embryos, n=143 cells). Scale bars = 30 μm (A, B, C, and D); 20 μm (Aʹ, Aʹʹ, Cʹ, and Cʹʹ); 15 μm (Bʹ, Bʹʹ, Dʹ, and Dʹʹ).

Previous experimental and modeling studies have proposed cell behaviors such as hypertrophic growth, rearrangement within the tissue, and changes to cell shape as potential contributors to chamber curvature morphogenesis. For example, in chick, localized increases to cardiomyocyte volume in the OC appear to play a significant role in the creation of the ventricular bend (Shi et al., 2014; Soufan et al., 2006). Additionally, in both chick and zebrafish, differing cell rearrangement patterns between the curvatures (Kawahira et al., 2020; Tessadori et al., 2021), as well as regionalized cell shape changes (Auman et al., 2007; Manasek et al., 1972), likely play into the pronounced expansion of the OC tissue. In zebrafish in particular, the strikingly divergent morphologies that ventricular OC and IC cardiomyocytes attain during curvature formation have suggested that active cell shape change is a critical driver of curvature formation. Specifically, examination of the myocardial surface of the zebrafish heart tube has highlighted that its cells are fairly uniform in both apical surface area and circularity (Auman et al., 2007). However, after curvature formation, OC cells have a significantly larger, more elongated apical surface area than do IC cells (Auman et al., 2007). Consistent with the idea that regionalized cell shape changes create chamber curvatures, many mutations that cause chamber dysmorphia also disrupt the acquisition of OC- or IC-linked cell shapes (Auman et al., 2024; Chi et al., 2008; Choudhry & Trede, 2013; Guzzolino et al., 2020; Halabi et al., 2022; Yang & Xu, 2012). However, the subcellular mechanisms that drive these changes to cardiomyocyte morphology have not been well defined.

The actomyosin cytoskeleton is an attractive candidate for mediating cell shape changes. Although capable of inducing shape change in several ways, the network is perhaps best known for its ability to contract and thus create localized tension within a cell, as in the apical constriction observed during *Drosophila* ventral furrow formation (Martin et al., 2010; Yevick et al., 2019). This type of actomyosin-derived tension is indeed employed at later stages of zebrafish cardiac morphogenesis, when differential tensions between cardiomyocytes are important for the decision to either trabeculate or remain in the compact layer (Priya et al., 2020). If actomyosin also plays an earlier role in curvature cell shape acquisition, we would expect to see different presentations of the network between curvatures. Indeed, in zebrafish, where there is a single layer of cardiomyocytes at this stage, visualization of the outer surface of the myocardium has highlighted what is interpreted as “cortical” F-actin in both curvatures (*i.e.* intense F-actin signal outlining each cell) and an additional pool of “cytoplasmic” F-actin running throughout OC cells (Deacon et al., 2010). Although visualizations of the F-actin cytoskeleton in chick are somewhat more challenging due to the bilayered myocardium, distinctions between the OC and IC have been resolved, with circumferentially aligned F-actin at cell-cell boundaries in the IC and a less organized network of F-actin in the OC (Itasaki et al., 1989; Shiraishi et al., 1992). Furthermore, actin dynamics appear to be necessary for the curvature formation process: in both chick and zebrafish, explanted hearts exposed to inhibitors of actin polymerization fail to achieve the stereotypic curved chamber contours (Itasaki et al., 1991; Latacha et al., 2005; Noël et al., 2013). Altogether, these data position actomyosin as a likely regulator of curvature formation, but how the divergence of the actomyosin landscape is regulated and whether the network mediates its effect on tissue shape via active cell shape change remain unknown.

Curvature formation takes place in a heart that actively beats and pumps blood, suggesting the possible involvement of biomechanical cues in regulating the cellular and subcellular changes that occur in OC and IC cardiomyocytes. It is well known that biomechanical inputs, such as fluid forces stemming from blood flowing through the endocardial lumen, tissue tension arising from the sarcomeric contraction of myocardial cells, and the anchoring of venous and arterial poles to the vascular system, influence several aspects of cardiac development (Andrews & Priya, 2024; Goenezen et al., 2012; Le Garrec et al., 2017; Lindsey et al., 2014; Taber, 2006; Voorhees & Han, 2015; Voronov et al., 2004). For example, altering the dynamics of blood flow in zebrafish can prevent both trabeculation (Lee et al., 2016; Peshkovsky et al., 2011; Staudt et al., 2014) and valve formation (Bartman et al., 2004; Hove et al., 2003; Vermot et al., 2009). In addition, zebrafish hearts that lack atrial contractility and thus have reduced blood flow through the ventricle exhibit OC cells with a reduced apical surface area (Auman et al., 2007). This phenotype highlights the importance of extrinsic factors for attaining curvature-linked cell shapes, but it is not yet clear how these forces are translated into the subcellular events that instigate changes to cardiomyocyte morphology.

While extrinsic forces clearly impact curvature formation, factors intrinsic to the myocardium are certainly also involved in regulating the cytoskeleton, changing cell behavior, and ultimately sculpting tissue shape. In considering intrinsic factors that could influence curvature morphogenesis, the TBX family of transcription factors stand out as potentially important contributors. A set of these factors work in concert to pattern the expression of genes associated with the chamber curvatures: TBX5 promotes expression of OC-related genes, whereas TBX2 and TBX3 suppress expression of these genes (Bruneau et al., 2001; Christoffels et al., 2004; Habets et al., 2002; Harrelson et al., 2004; He et al., 2011; Hiroi et al., 2001; Ivanovitch et al., 2017; Sedletcaia & Evans, 2011; Tessadori et al., 2021). For example, the combination of these three factors determine the OC-enriched expression of *Nppa* in mice (Christoffels et al., 2004; Habets et al., 2002; Harrelson et al., 2004; Hiroi et al., 2001). In zebrafish, expression of *nppa* is already enriched in the future ventricular OC region of the linear heart tube, well before curvatures take shape (Auman et al., 2007). Although loss-of-function analyses indicate that *nppa* itself is not necessary for curvature formation (Grassini et al., 2018), its early prepattern signifies that there may be other factors regulated by Tbx factors that could influence the ability of OC or IC cells to change their morphology. This idea is especially intriguing in light of recent work in mouse showing that TBX5 influences epithelial tension and cell morphology in the second heart field (Guijarro et al., 2024).

In this study, we delve into new aspects of how OC and IC cardiomyocyte morphologies diverge during curvature formation in zebrafish. Specifically, by studying OC and IC cell shapes in three dimensions, we have found that OC cardiomyocytes expand primarily in the planar axis, becoming more squamous as curvatures form, while IC cardiomyocytes extend primarily in the apicobasal axis, becoming more cuboidal. We show that these cellular changes are preceded by regionalized patterns of subcellular actomyosin organization, and that actomyosin plays a cell-intrinsic role in determining cell shape. Intriguingly, this role appears to be in the induction of planar spread, rather than the role of constriction for which the actomyosin cytoskeleton is so well known. Finally, we show that blood flow through the ventricle and *tbx5a* each promote the curvature-associated divergence of actomyosin organization and cardiomyocyte morphologies, with particularly prominent effects in the OC. From these findings, we propose that several extrinsic and intrinsic factors converge to control the cytoskeletal dynamics that govern cell shape change, and that these changes to cardiomyocyte morphologies are a critical component in sculpting the characteristic contours of the cardiac chambers.

## Results

### Curvature formation coincides with the divergence of ventricular OC and IC cardiomyocyte morphologies

To study the cellular and molecular underpinnings of curvature formation, we first needed to develop a working method to standardize the boundaries of the ventricular curvatures. For this purpose, we used a combination of gene expression patterns and measurements of morphological features to guide boundary placement in the embryonic zebrafish ventricle at 48 hours post-fertilization (hpf) (Figure 1—figure supplement 1A-C). For gene expression, we relied on *nppa*, which is enriched in the OC of the ventricle (Auman et al., 2007; Christoffels et al., 2000). Generally, the highest *nppa*-expressing area of the ventricle was deemed to be the OC, and the lowest *nppa*-expressing area was deemed to be the IC (Figure 1—figure supplement 1A and C). After examining the *nppa* expression pattern in 12 wild-type ventricles at 48 hpf, we used the typical territory of *nppa* expression and its spatial relation to the atrioventricular canal (AVC) and distal end of the outflow tract (OFT) to determine the morphological features of the heart that could serve as boundaries for the curvatures (see Materials and Methods for more detail). Once we had drawn these boundaries, we found that the IC region coincided satisfyingly with the expression of *mb* (Figure 1—figure supplement 1B), a gene previously noted to be enriched in the ventricular IC (Burkhard & Bakkers, 2018). In a wild-type heart at 48 hpf, the OC region defined by this method typically includes ∼50-60 cardiomyocytes, and the IC region typically includes ∼25-30 cardiomyocytes. We modified this method slightly to determine the curvature boundaries in wild-type hearts at 36 hpf (Figure 1—figure supplement 1D; see Materials and Methods for more detail), and we used these rules to bound the curvatures in mutant hearts as well (see subsequent figures, including Figures 5 and 6). Although we think that these boundaries encompass the territories that are important for the study of ventricular curvature formation, it is possible that they also include some less relevant areas. Nevertheless, this method provides a useful and reproducible strategy for outlining the ventricular OC and IC in multiple contexts.

Our previous work has shown that certain characteristics of cardiomyocyte shape, such as apical surface area and circularity, are uniform throughout the myocardium at the linear heart tube stage (24 hpf), but eventually diverge in the ventricular curvatures by 48 hpf (Auman et al., 2007). Employing our new definitions of the OC and IC, we aimed to understand this divergence with greater temporal detail by looking at a developmental stage midway through curvature formation, in both the planar (X/Y) and apicobasal (Z) axes. At 37 hpf, when curvatures have just started to take shape, we found that OC and IC cardiomyocytes exhibit only a modest difference in morphologies (Figure 1A, B, and E-H). OC cells have a slightly but significantly larger apical surface area than IC cells, whereas the circularity of the apical surfaces are comparable (Figure 1A, E, and H). Additionally, cell thickness, as measured by apicobasal length, is similar between OC and IC cells at this stage (Figure 1B and F). Both OC and IC cells increase in volume between 37 and 48 hpf and extend this new volume in both planar and apicobasal axes (Figure 1C, D, and G). However, OC cells expand along their planar axis to a greater extent than do IC cells (Figure 1A and C), as observed by a greater increase in apical surface area (Figure 1E), and they become more elongated, typically along their circumferential axis (Figure 1C and H). By contrast, IC cells become taller along their apicobasal axis and remain more circular than OC cells (Figure 1B, D, F, and H). Together, these data suggest that cells in the developing curvatures grow to similar degrees during this time, but they choose to allocate their new volume along different axes: along the planar axis for OC cells and along the apicobasal axis for IC cells. As a consequence, OC cells become more squamous, and IC cells become more cuboidal. These data also place the onset of cardiomyocyte shape change at an early stage when the heart is still relatively tubular, highlighting active cell shape change as a potential driver of curvature formation.

### Divergence of cardiomyocyte morphologies is preceded by changes to actomyosin organization

To understand the cellular mechanisms behind the divergence in OC and IC cardiomyocyte shapes, we first wanted to identify when previously reported cytoskeletal characteristics of the curvatures diverged (Deacon et al., 2010; Itasaki et al., 1989; Shiraishi et al., 1992). Specifically, do the portions of the ventricle that will become the IC and OC exhibit differential actomyosin landscapes as soon as the linear heart tube is formed, or do they diverge sometime during curvature formation? If the latter is true, does this divergence precede the appearance of the observed differences between OC and IC cell shapes? Throughout the process of curvature formation, we found that F-actin primarily resides along the inner surface of the cardiomyocyte membranes. At 28 hpf, when a small bulge of OC is just becoming evident (Figure 2—figure supplement 1A), subcellular distribution of F-actin appears fairly uniform in the presumptive OC and IC (Figure 2—figure supplement 1B). As observed previously (Merks et al., 2018), proximal ventricular cells (those closer to the AVC) exhibit F-actin mostly at their basal membrane (Figure 2—figure supplement 1Bʹʹ and Bʹʹʹʹ), whereas distal ventricular cells (those closer to the OFT) exhibit F-actin distributed around all of the surfaces of their membrane, with particularly strong presence at the apical surface (Figure 2—figure supplement 1Bʹ and Bʹʹʹ). At 36 hpf, when OC and IC cells have begun to show small morphological differences (Figure 1A, B, and E-H), the F-actin in OC cells remains more enriched at the basal surface, particularly in the proximal ventricle (Figure 2—figure supplement 1Dʹ and Dʹʹ). In IC cells, F-actin is distributed more equally around the membrane surfaces, with a substantial amount at the lateral and apical surfaces (Figure 2—figure supplement 1Dʹʹʹ and Dʹʹʹʹ). At 48 hpf, OC and IC cells look fairly similar to each other, with F-actin distributed around all membrane surfaces and particularly intense signal at the lateral membranes (Figure 2—figure supplement 1E and F). Our observations that OC and IC cells begin with similar F-actin organization but diverge at 36 hpf, only to converge again by 48 hpf, drew our attention to 36 hpf as a stage worthy of more intense study. These findings correspond well with published visualizations of the myocardial surface showing that F-actin organization differs between OC and IC cells at 36 hpf (but not at 24 hpf) (Deacon et al., 2010) and that *myosin vb* mutants, which fail to undergo OC cell planar expansion, exhibit aberrant F-actin organization in OC cells at 36 hpf (Grassini et al., 2019).

To better understand these qualitative findings, we adopted a more quantitative approach to the examination of the subcellular localization of F-actin and pMyosin in individual cardiomyocytes of the OC and IC (Figure 2A-E). For every cardiomyocyte analyzed, we quantified signal intensities to reveal what proportion of the total F-actin and pMyosin is localized to its basal, lateral, and apical membranes (Figure 2B; see Materials and Methods for more detail). Using this approach, we found that OC cells have a higher proportion of basal actomyosin than do IC cells (Figure 2C-E), whereas IC cells have a higher proportion of apical and lateral actomyosin than do OC cells (Figure 2C-E). In addition, we wondered whether we could observe disparities between cells in the proximal OC/IC and cells in the distal OC/IC, as we could in Figure 2—figure supplement 1. Comparing these subsets of cells, we found that proximodistal differences are quite clear in the OC (Figure 2—figure supplement 2 and Figure 2—figure supplement 3). In proximal OC cells, F-actin and pMyosin are highly enriched at the basal surface, whereas in the distal OC, cells have a significantly lower proportion of actomyosin at the basal membrane and a higher proportion of actomyosin at the lateral and apical membranes, giving an overall organization more similar to cells of the IC, which do not exhibit striking differences between proximal and distal regions. These data from individual cells (Figure 2) support what we observed at the tissue level (Figure 2—figure supplement 1) and highlight a potentially important aspect of cytoskeletal dynamics during a critical period of curvature formation.

**Figure 2.**
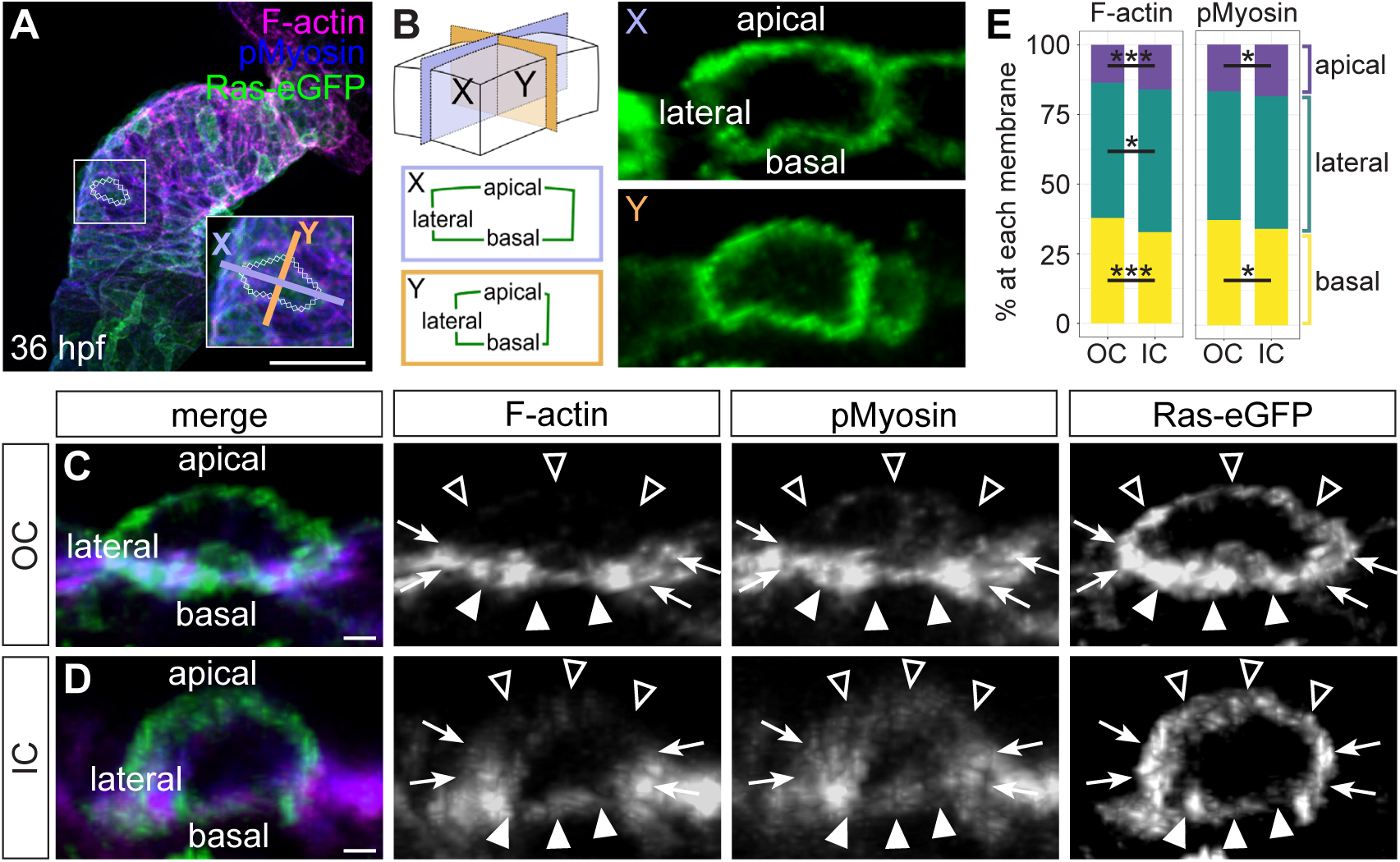
OC and IC cardiomyocytes exhibit differential localization of subcellular actomyosin during early curvature formation. (**A**) Whole heart from a 36 hpf embryo carrying *Tg(myl7:eGFP-Hsa.HRAS)* (D’Amico et al., 2007), immunostained for membrane-bound GFP and pMyosin and stained with Phalloidin to label F-actin. Inset magnifies a single cardiomyocyte. (**B**) Visual representation of approach for measuring the percentages of F-actin or pMyosin at each membrane in individual cardiomyocytes. Briefly, each OC or IC cardiomyocyte was bisected in two axes, resulting in two cross-sections for each cell (X, purple; Y, orange). The cell boundaries in X and Y are visible due to eGFP localization to the membranes. In these images, the basal membrane is always at the bottom. (**C and D**) Cross-sections through representative individual cardiomyocytes from the OC (C) or IC (D). Empty arrowheads: apical membranes. Filled arrowheads: basal membranes. Arrows: lateral membranes. (**E**) Stacked bar charts showing the mean percentage of F-actin or pMyosin at each membrane. Refer to Tables S1 and S2 for summary statistics. * denotes p < 0.05 and *** denotes p < 0.001, Wilcoxon test. OC (N=10 embryos, n=199 cells); IC (N=10 embryos, n=123 cells). Scale bars = 50 μm (A); 2 μm (C and D).

Altogether, our observations indicate that, in the linear heart tube, cardiomyocytes in the future OC and IC have similar actomyosin organization, and that the greater disparity lies between the proximal ventricle and the distal ventricle. As chambers begin to form, cells in the OC (particularly those in the proximal region) retain basal enrichment of actomyosin, while cells in the IC accumulate actomyosin at their apical and lateral surfaces. Notably, the process of acquiring these differential cytoskeletal landscapes divergence closely precedes the initiation of the cell shape changes observed in Figure 1. These data suggest that the actomyosin network in cardiomyocytes is differentially regulated depending upon their location within the ventricle. In addition, these data imply potentially distinct roles for actomyosin in the curvatures, given that the precise location of actomyosin within a cell determines its ability to engage drivers of specific cell behaviors.

### Modulation of actomyosin dynamics dampens the divergence of OC and IC cardiomyocte morphologies

Given the intriguing regionalized patterns of subcellular actomyosin localization that arise during chamber curvature formation, we wondered whether the actomyosin cytoskeleton plays a role in the divergence of curvature cell shapes. Previous studies in chick and zebrafish have shown that actin polymerization is crucial for chamber curvature formation (Itasaki et al., 1991; Latacha et al., 2005; Noël et al., 2013), but these studies did not delve into the impact of actin polymerization on cardiomyocyte morphology. We treated wild-type embryos from 24-36 hpf or from 36-48 hpf with either Latrunculin B (Lat B) or Blebbistatin (Bleb) to block actin polymerization or non-muscle myosin II (NMII) activity, respectively (Figure 3—figure supplement 1). In embryos treated with either LatB or Bleb from 24-36 hpf, OC cardiomyocytes exhibited significantly smaller apical surface areas when compared with DMSO-treated controls (Figure 3—figure supplement 1A, B, D, E, F, and H). In contrast to the effects of treatment from 24-36 hpf, Lat B treatment from 36-48 hpf had no effect on planar expansion of OC cells (Figure 3—figure supplement 1C and D), and, curiously, Bleb treatment from 36-48 hpf resulted in excessive planar expansion (Figure 3—figure supplement 1G and H). These data highlight 24-36 hpf as a particularly important window for actomyosin activity in promoting OC cell planar expansion. They also hint at a second phase of NMII activity between 36 and 48 hpf, during which Myosin-based contractility might be important for maintaining cell shape.

We next wanted to evaluate whether actomyosin activity is required in cardiomyocytes themselves, or if the effect we observe on cardiomyocyte shape upon LatB or Bleb treatment is due to a required function in another tissue, such as the endocardium. To assess this, we used established transgenic lines to express different versions of the non-muscle myosin light chain 9 gene (*myl9*) driven throughout the myocardium by the cardiomyocyte-specific *myl7* promoter: wild-type *myl9* (*Tg(myl7:WT-myl9-mScarlet)*), a dominant negative version of *myl9* (*Tg(myl7:DN-myl9-eGFP)*), or a constitutively active version of *myl9* (*Tg(myl7:CA-myl9-eGFP)*) (Priya et al., 2020). When assessing cardiomyocyte morphologies at 48 hpf, we found that expression of *WT-myl9* resulted in cell morphologies fairly reminiscent of those in wild-type embryos (Figure 1C-F, and H and Figure 3A, B, and G-I). By contrast, expression of *DN-myl9* resulted in OC cardiomyocytes with decreased apical surface area and increased apicobasal length when compared with OC cells expressing *WT-myl9* (Figure 3C, D, G, and H). Together, these changes resulted in a more cuboidal phenotype. Additionally, IC cells expressing *DN-myl9* exhibited a slightly (but significantly) decreased apical surface area and equivalent apicobasal length when compared with those expressing *WT-myl9* (Figure 3C, D, G, and H). Expression of *CA-myl9* resulted in OC cells with a slightly (but significantly) decreased apical surface area and increased apicobasal length when compared with those expressing *WT-myl9*, whereas IC cells expressing *CA-myl9* exhibited a similar apical surface area and slightly (but significantly) decreased apicobasal length when compared with those expressing *WT-myl9* (Figure 3E, F, G, and H). Interestingly, both OC and IC cells expressing *CA-myl9* were more elongated than those expressing *WT-myl9* (Figure 3I). Altogether, these data suggest that actomyosin dynamics are indeed important within the myocardium to support the divergence of OC and IC cell morphologies, and that modulating NMII activity produces OC and IC cells with ultimately similar shapes.

**Figure 3.**
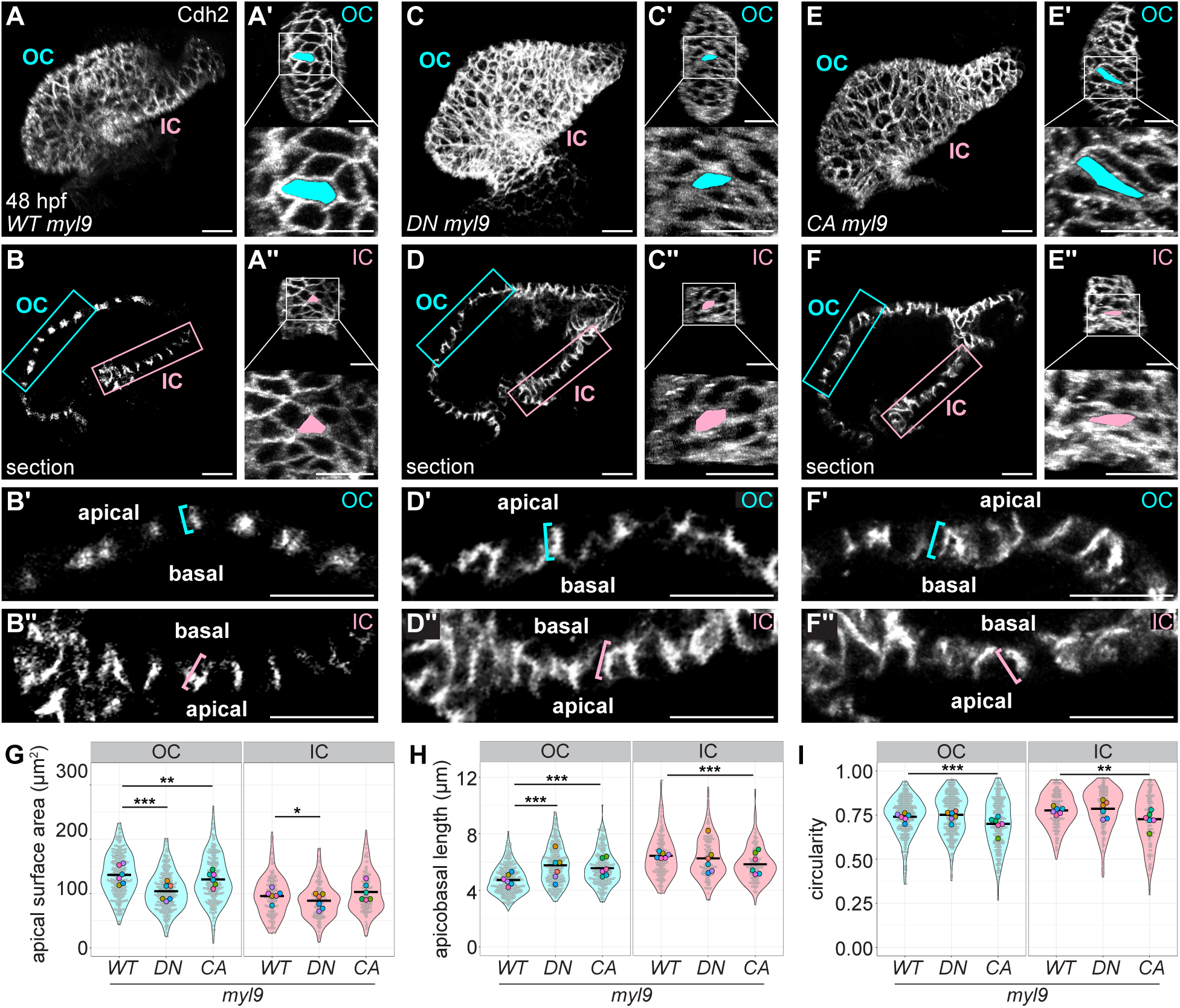
Tissue-specific modulation of NMII activity dampens the divergence of OC and IC cardiomyocyte morphologies. (**A**, **C**, **and E**) 3D reconstructions of 48 hpf hearts expressing *Tg(myl7:WT-myl9-mScarlet)* (A), *Tg(myl7:DN-myl9-eGFP)* (C), or *Tg(myl7:CA-myl9-eGFP)* (E). Immunostaining for Cdh2 labels lateral membranes of cardiomyocytes. OCs (Aʹ, Cʹ, and Eʹ) and ICs (Aʹʹ, Cʹʹ, and Eʹʹ) are shown for hearts in (A, C, and E). Insets show higher magnification. Apical surface area of an individual cardiomyocyte is illustrated by blue or pink fill. (**B**, **D**, **and F**) Sections through hearts in (A, C, and E). (Bʹ, Bʹʹ, Dʹ, Dʹʹ, Fʹ, and Fʹʹ) are magnified views of blue (OC) and pink (IC) boxed regions in (B, D, and F); blue and pink brackets highlight apicobasal length of individual cardiomyocytes. (**G-I**) Violin plots compare apical surface area, apicobasal length, or circularity of cardiomyocytes between embryos expressing the different transgenes, split by curvature. Circularity is calculated as 4⫪(A/P^2^). Each small grey dot represents an individual cell, each black bar represents the mean of values from individual cells, and each large colored dot represents the mean of all values from an individual embryo. * denotes p < 0.05, ** denotes p < 0.01, and *** denotes p < 0.001, Wilcoxon test. *Tg(myl7:WT-myl9-mScarlet)* OC (N=6 embryos, n=308 cells); *Tg(myl7:WT-myl9-mScarlet)* IC (N=6 embryos, n=159 cells); *Tg(myl7:DN-myl9-eGFP)* OC (N=6 embryos, n=298 cells); *Tg(myl7:DN-myl9-eGFP)* IC (N=6 embryos, n=143 cells); *Tg(myl7:CA-myl9-eGFP)* OC (N=6 embryos, n=253 cells); *Tg(myl7:CA-myl9-eGFP)* IC (N=6 embryos, n=123 cells). Scale bars = 20 μm.

To better understand the effect of these transgenes on cardiomyocyte shape, we wondered whether the changes in NMII activity that led to cell shape alterations also had a parallel effect on the F-actin network. We were intrigued to find that expression of *DN-myl9* or *CA-myl9* reduced the overall amount of F-actin in the ventricle at 36 hpf (Figure 3—figure supplement 2). We further found that subcellular localization of F-actin was disrupted in both the *DN-myl9* and *CA-myl9* contexts, and in cardiomyocytes of both curvatures (Figure 3—figure supplement 3). Of particular interest, we found a shift in F-actin enrichment from the basal membrane to the apical and lateral membranes in *DN-myl9-*expressing (Figure 3—figure supplement 3B and G) and *CA-myl9*-expressing (Figure 3—figure supplement 3C and G) OC cells. This shift prompted us to quantify the ratio of the amount of F-actin at the basal membrane to the amount of F-actin at the apical and lateral membranes (basal / (apical + lateral)). This ratio revealed a striking shift away from the basal enrichment normally seen in *WT-myl9*-expressing OC cells (Figure 3—figure supplement 3H). From other contexts, we know that Myosin itself can alter F-actin characteristics. For example, it can support F-actin bundles by crosslinking (Bridgman et al., 2001; Cai et al., 2010; Laevsky & Knecht, 2003), can induce disassembly of stress fibers (Huang et al., 2021; Matsui et al., 2011; Wilson et al., 2010), and can organize F-actin at specific locations within the cell (Murthy & Wadsworth, 2005; Yamashiro et al., 2018; Zhou & Wang, 2008). We therefore posit that altering the function of the Myosin network by expression of *DN-my9* or *CA-myl9* in turn impacts the F-actin landscape.

### NMII function plays a cell-intrinsic role in promoting divergence of OC and IC cardiomyocyte morphologies

Having seen that cardiomyocytes fail to attain expected shapes when either a dominant negative or constitutively active form of NMII is expressed throughout the myocardium (Figure 3), we were curious about whether actomyosin plays a cell-intrinsic role in promoting OC and IC cell shape divergence, or, rather, if NMII-driven tissue-level tensions are responsible for the acquisition of cell morphologies. We therefore performed blastomere transplantation to create genetically mosaic embryos (Figure 4A). In these experiments, we analyzed hearts that contained donor-derived cardiomyocytes expressing *CA-myl9* or *DN-myl9* surrounded by host-derived wild-type cardiomyocytes (Figure 4B-M; see Materials and Methods regarding how specific cardiomyocytes were chosen for analysis). In comparison with their wild-type host-derived neighbors, *CA-myl9-*expressing OC cells had a dramatically decreased apical surface area and increased apicobasal length, resulting a strikingly more cuboidal phenotype (Figure 4B, C, E, and F). In addition, *CA-myl9*-expressing donor-derived OC cells were significantly more circular than their wild-type neighbors (Figure 4B, C, and G). In the IC, *CA-myl9*-expressing donor-derived cells were also more cuboidal than their host-derived wild-type neighbors, but the magnitude of this effect was less than in the OC (Figure 4B and D-F). This OC phenotype is particularly interesting, given its contrast to the effect of uniform expression of *CA-myl9* throughout the myocardium (Figure 3E-I), in which cardiomyocytes were only slightly more cuboidal and were in fact more elongated than *WT-myl9*-expressing cells. This disparity highlights the importance of tissue-scale tension and how individual cell shape can be influenced depending on whether neighboring cells exhibit normal or increased tensions.

**Figure 4.**
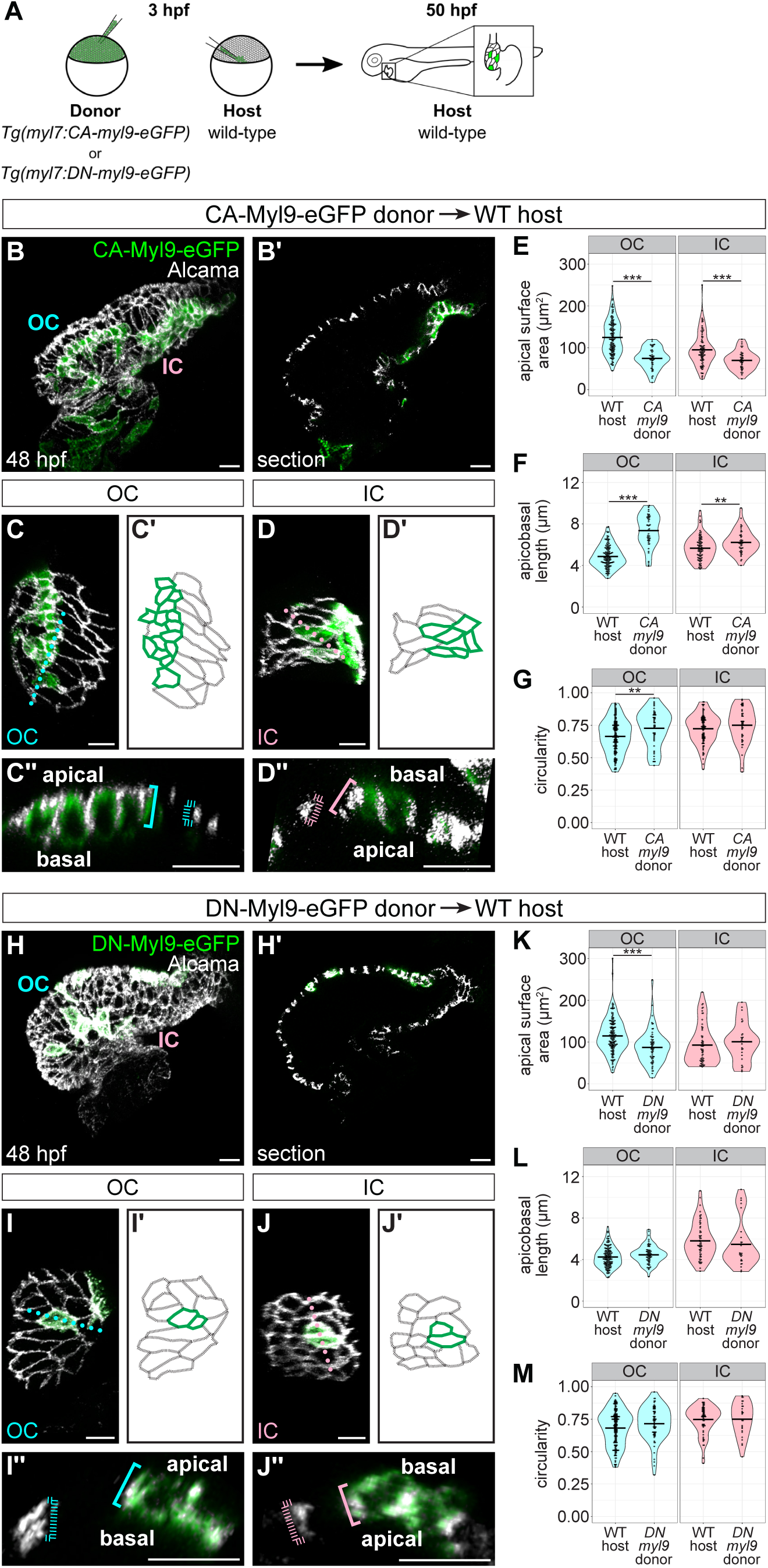
Cell-intrinsic modulation of NMII activity alters cardiomyocyte shape. (A) Schematic of blastomere transplantation experiment. (**B and H**) 3D reconstructions of 48 hpf wild-type (WT) host hearts containing donor-derived cardiomyocytes expressing *Tg(myl7:CA-myl9-eGFP)* (B) or *Tg(myl7:DN-myl9-eGFP)* (H); immunostaining for Alcama labels lateral membranes of cardiomyocytes. (Bʹ and Hʹ) Sections through hearts in (B and H). OC (**C and I**) and IC (**D and J**) are shown from additional mosaic hearts other than those in (B and H). (Cʹ, Dʹ, Iʹ, and Jʹ) Tracings of the cardiomyocytes in (C, D, I, and J). Green outlines indicate donor-derived cardiomyocytes; black outlines indicate host-derived cardiomyocytes. (Cʹʹ, Dʹʹ, Iʹʹ, and Jʹʹ) Cross-sections through positions indicated by dotted lines in (C, D, I, and J). Blue and pink brackets highlight apicobasal length of individual cardiomyocytes, with dashed brackets for host-derived cardiomyocytes and solid brackets for donor-derived cardiomyocytes. (**E-G and K-M**) Violin plots compare apical surface area, apicobasal length, and circularity of host-derived cardiomyocytes to those of donor-derived cardiomyocytes. Each dot represents an individual cell. ** denotes p < 0.01 and *** denotes p < 0.001, Wilcoxon test. For *Tg(myl7:CA-myl9-eGFP)* into WT transplants: host OC (N=8 embryos, n=145 cells); donor OC (N=8 embryos, n=49 cells); host IC (N=7 embryos, n=91 cells); donor IC (N=7 embryos, n=44 cells). For *Tg(myl7:DN-myl9-eGFP)* into WT transplants: host OC (N=10 embryos, n=156 cells); donor OC (N=10 embryos, n=65 cells); host IC (N=8 embryos, n=60 cells); donor IC (N=8 embryos, n=27 cells). Scale bars = 15 μm.

In the dominant negative situation, we found that *DN-myl9-*expressing OC cardiomyocytes had a smaller apical surface area when compared with *WT-myl9*-expressing cells (Figure 4H, I, and K), but that apicobasal length was unchanged (Figure 4H, I, and L). In the IC, we found that *DN-myl9*-expressing donor-derived cells were very similar to their host-derived neighbors (Figure 4H and J-M). This OC cell phenotype is fairly similar to the effect of uniform expression of *DN-myl9* throughout the myocardium (Figure 3C, D, and G-I). These data indicate that NMII plays a cell-intrinsic role in promoting the planar expansion of OC cells and that this role is not altered by the degree of tension in neighboring cells. Taken together, our analysis of cardiomyocytes expressing *myl9* transgenes emphasizes that actomyosin activity supports the divergence of OC and IC cell shapes by promoting the squamous identity of OC cells in a cell-intrinsic manner, perhaps by facilitating active expansion in the planar axis, as has been hypothesized (Latacha et al., 2005). In contrast, IC cells appear to be affected only when actomyosin contractility is intensified throughout the process of curvature formation, as is the case with *CA-myl9* expression.

### Divergence of OC and IC actomyosin organization and cardiomyocyte morphology depends upon biomechanical cues

Our results thus far had highlighted actomyosin organization and function as drivers of cell shape change during curvature formation, and we next wanted to know what types of upstream regulators might control these subcellular features. We were particularly curious to evaluate how blood flow might affect actomyosin localization and cardiomyocyte morphology, as our previous work has shown that *myosin heavy chain 6* (*myh6*) mutants (Berdougo et al., 2003), which lack an atrium-specific myosin heavy chain and therefore lack atrial contractility, have reduced blood flow through the ventricle as well as reduced apical surface area of OC cardiomyocytes (Auman et al., 2007). Consistent with previous findings, we found that OC cells of *myh6* mutants fail to extend along the planar axis (Figure 5A, C, and E). In addition, we observed that *myh6* OC cells also extend inappropriately along the apicobasal axis (Figure 5B, D, and F). Taken together, these data indicate that *myh6* OC cells acquire a more cuboidal phenotype than their wild-type counterparts. In contrast to the OC results, we found that the apical surface area and apicobasal length of *myh6* IC cells are indistinguishable from those of wild-type IC cells (Figure 5A-F); however, both OC and IC cells exhibit a modest but significant decrease in volume in *myh6* mutants (Figure 5G).

**Figure 5.**
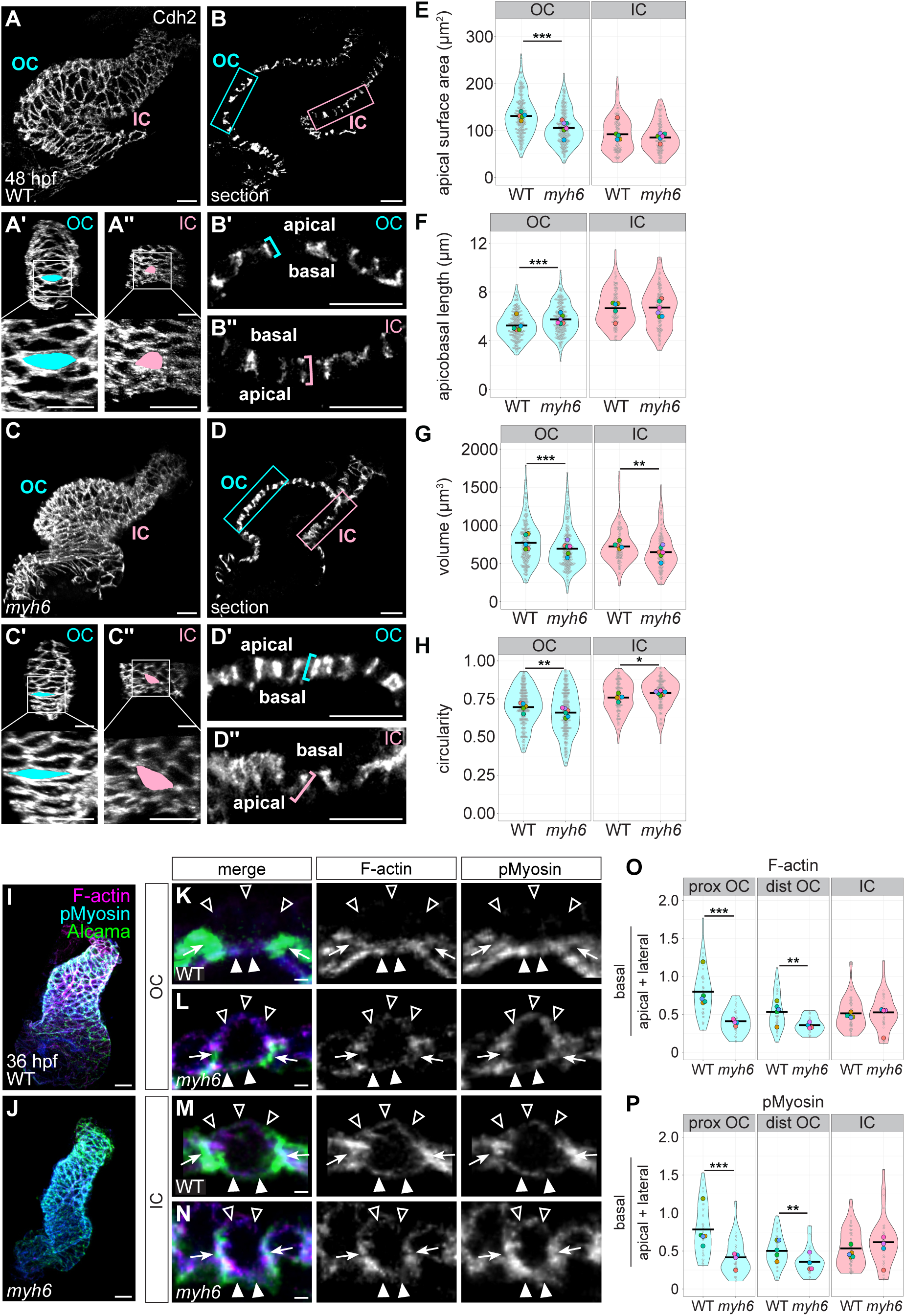
Reduced blood flow inhibits the divergence of OC and IC cardiomyocyte shapes and actomyosin organization. (**A-H**) Comparison of cardiomyocyte morphologies in wild-type and *myh6* mutant hearts. (A and C) 3D reconstructions of wild-type (A) and *myh6* mutant (C) hearts at 48 hpf; immunostaining for Cdh2 labels lateral membranes of cardiomyocytes. OCs (Aʹ and Cʹ) and ICs (Aʹʹ and Cʹʹ) are shown for hearts in (A and C). Insets show higher magnification. Apical surface area of an individual cardiomyocyte is illustrated by blue or pink fill. (B and D) Sections through hearts in (A and C). (Bʹ, Bʹʹ, Dʹ, and Dʹʹ) Magnified views of blue (OC) and pink (IC) boxed regions in (B and D); blue and pink brackets highlight apicobasal length of individual cells. (E-H) Violin plots compare apical surface area, apicobasal length, volume, and circularity for cells from wild-type and *myh6* mutant hearts. (H) Note that *myh6* OC cardiomyocytes are more elongated than wild-type OC cardiomyocytes, a finding that contrasts with our previous work (Auman et al., 2007). We posit that this discrepancy could arise from changes in the genetic background over time or to differences in how data were collected (*e.g.* where the OC boundary was drawn or how individual cardiomyocytes were measured). (**I-P**) Comparison of subcellular actomyosin organization in wild-type and *myh6* mutant cardiomyocytes. (I and J) Wild-type (I) and *myh6* mutant (J) hearts at 36 hpf, immunostained for pMyosin and stained with Phalloidin to label F-actin. Immunostaining for Alcama labels lateral membranes of cardiomyocytes. (K-N) Cross-sections through representative cardiomyocytes from the OC (K and L) and IC (M and N) of 36 hpf wild-type (K and M) and *myh6* mutant (L and N) hearts. Empty arrowheads: apical membranes. Filled arrowheads: basal membranes. Arrows: lateral membranes. (O) Violin plots show calculated values of (mean basal F-actin / (mean apical F-actin + mean lateral F-actin)) for individual cells. (P) Violin plots of calculated values as in (O), but for pMyosin. For all violin plots, each small grey dot represents an individual cell, each black bar represents the mean of values from individual cells, and each large colored dot represents the mean of all values from an individual embryo; * denotes p < 0.05, ** denotes p < 0.01, and *** denotes p < 0.001, Wilcoxon test. For morphometrics: wild-type OC (N=5 embryos, 267 cells); *myh6* OC (N=6 embryos, n=275 cells); wild-type IC (N=5 embryos, n=120 cells); *myh6* IC (N=6 embryos, n=135 cells). For actomyosin localization: wild-type proximal OC (N=5 embryos, n=39 cells); *myh6* proximal OC (N=4 embryos, n=32 cells); wild-type distal OC (N=5 embryos, n=41 cells); *myh6* distal OC (N=4 embryos, n=24 cells); wild-type IC (N=5 embryos, n=54 cells); *myh6* IC (N=4 embryos, n=35 cells). Scale bars = 20 μm (A-D, I, and J); 2 μm (K-N).

Given the effect of reduced blood flow on OC cell morphology, we next asked if this phenotype is coupled with a disordered actomyosin cytoskeleton. Although *myh6* mutant hearts are morphologically similar to wild-type hearts up to 36 hpf (Figure 5I and J) (Auman et al., 2007), we found that the overall levels of F-actin in the ventricular myocardium in *myh6* mutants at 36 hpf are reduced relative to wild-type (similar to our observations in *DN-myl9* and *CA-myl9* transgenic hearts (Figure 3—figure supplement 2)), whereas the levels of pMyosin are comparable to wild-type (Figure 5—figure supplement 1A-D). In addition to an overall deficit in F-actin, we found mislocalization of subcellular actomyosin in *myh6* mutant OC cells (Figure 5I-L, O, and P): both F-actin and pMyosin shift away from the basal surface and to the lateral and apical surfaces in the proximal OC and distal OC, with a particularly striking shift in the proximal OC (Figure 5K, L, O, and P; Figure 5—figure supplement 1E). In contrast to this OC cell phenotype, both F-actin and pMyosin organization in *myh6* mutant IC cells are comparable to wild-type (Figure 5M-P; Figure 5—figure supplement 1E). Taken together, these data highlight a previously unappreciated role for extrinsic factors such as the biomechanical cues produced by blood flow in ensuring the regionalization of cytoskeletal landscapes and, subsequently, the planar expansion of OC cells.

### Cell-autonomous function of *tbx5a* contributes to the divergence of OC and IC actomyosin organization and cardiomyocyte morphology

Alongside considering the extrinsic factors that influence curvature formation, we also wondered what types of intrinsic factors, operating within individual cells, could impact the divergence of cardiomyocyte cell shapes and actomyosin organization. We were particularly curious about the influence of the T-box transcription factor gene *tbx5a*. In addition to the known role for *Tbx5* in promoting OC transcriptional programs (Bruneau et al., 2001; He et al., 2011; Hiroi et al., 2001), zebrafish homozygous for mutant alleles of *tbx5a* exhibit a highly dysmorphic heart by 48 hpf (Garrity et al., 2002; Tessadori et al., 2021). As reported previously (Garrity et al., 2002), the outward bulge of the *tbx5a* mutant OC is substantially reduced and the *tbx5a* mutant IC exhibits less of a kink when compared with a wild-type heart (Figure 6A and C). We therefore investigated whether *tbx5a* mutant cardiomyocytes underwent the stereotypical divergence in shape associated with the ventricular curvatures. At the cellular scale, and similar to the phenotypes seen in *DN-myl9*-expressing hearts (Figure 3A-D, G, and H) and in *myh6* mutants (Figure 5A-F), OC cells in *tbx5a* mutants fail to expand in the planar axis and undergo aberrant apicobasal expansion (Figure 6A-F). This combination results in *tbx5a* mutant OC cells with a more cuboidal phenotype than wild-type OC cells. Interestingly, even though at the tissue level *tbx5a* mutant ICs fail to form the stereotypic kink (Figure 6C), IC cell shapes in *tbx5a* mutants appear relatively normal with just a slight trend toward reduction in apicobasal length (Figure 6A-F). In contrast to *myh6* mutants (Figure 5G and H), *tbx5a* OC and IC cells exhibit volumes and circularity similar to that of wild-type cells (Figure 6G and H).

**Figure 6.**
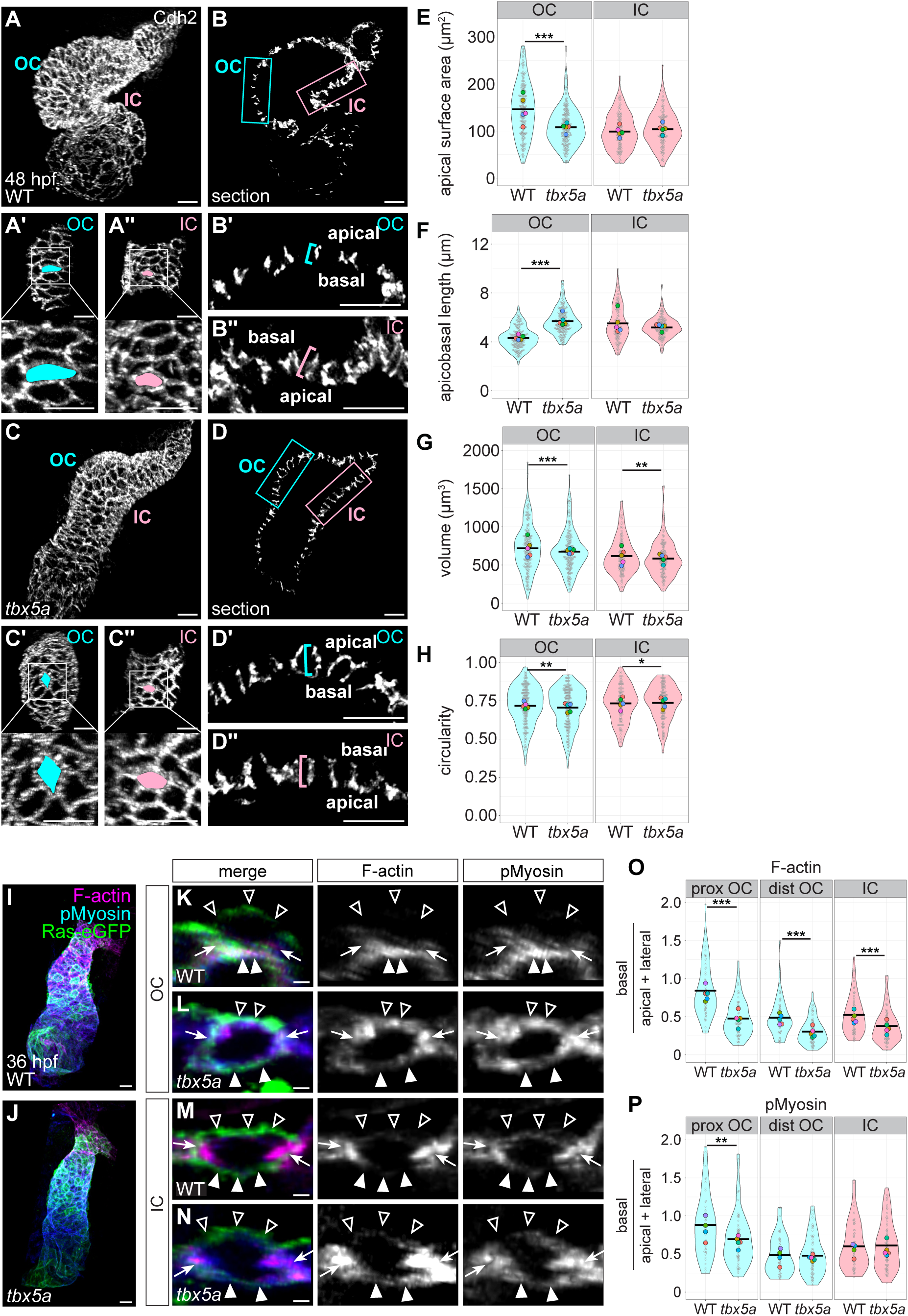
*tbx5a* regulates pathways that support the divergence of OC and IC cardiomyocyte morphologies and actomyosin organization. **(A-H)** Comparison of cardiomyocyte morphologies in wild-type and *tbx5a* mutant hearts. (A and C) 3D reconstructions of wild-type (A) and *tbx5a* mutant (C) hearts at 48 hpf; immunostaining for Cdh2 labels lateral membranes of cardiomyocytes. OCs (Aʹ and Cʹ) and ICs (Aʹʹ and Cʹʹ) are shown for hearts in (A and C). Insets show higher magnification. Apical surface area of an individual cardiomyocyte is illustrated by blue or pink fill. (B and D) Sections through hearts in (A and C). (Bʹ, Bʹʹ, Dʹ, and Dʹʹ) Magnified views of blue (OC) and pink (IC) boxed regions in (B and D); blue and pink brackets highlight apicobasal length of individual cardiomyocytes. (E-H) Violin plots compare apical surface area, apicobasal length, volume, and circularity for cardiomyocytes from wild-type and *tbx5a* mutant hearts. **(I-P)** Comparison of subcellular actomyosin organization in wild-type and *tbx5a* mutant cardiomyocytes. (I and J) Wild-type (I) and *tbx5a* mutant (J) hearts at 36 hpf, immunostained for membrane-bound eGFP and pMyosin, and stained with Phalloidin to label F-actin. (K-N) Cross-sections through representative cardiomyocytes from the OC (K and L) and IC (M and N) of 36 hpf wild-type (K and M) and *tbx5a* mutant (L and N) hearts. Empty arrowheads: apical membranes. Filled arrowheads: basal membranes. Arrows: lateral membranes. (O) Violin plots show calculated values of (mean basal F-actin / (mean apical F-actin + mean lateral F-actin)) for individual cells. (P) Violin plots of calculated values as in (O), but for pMyosin. For all violin plots, each small grey dot represents an individual cell, each black bar represents the mean of values from individual cells, and each large colored dot represents the mean of all values from an individual embryo. ** denotes p < 0.01 and *** denotes p < 0.001, Wilcoxon test. For morphometrics: wild-type OC (N=5 embryos, n=214 cells); *tbx5a* OC (N=5 embryos, n=220 cells); wild-type IC (N=5 embryos, n=126 cells); *tbx5a* IC (N=5 embryos, n=141 cells). For actomyosin localization: wild-type proximal OC (N=5 embryos, n=58 cells); *tbx5a* proximal OC (N=5 embryos, n=64 cells); wild-type distal OC (N=5 embryos, n=61 cells); *tbx5a* distal OC (N=5 embryos, n=67 cells); wild-type IC (N=5 embryos, n=69 cells); *tbx5a* IC (N=5 embryos, n=81 cells). Scale bars = 20 μm (A-D, I, and J); 2 μm (K-N).

We next wondered whether the effect of *tbx5a* loss-of-function on OC cardiomyocyte morphology was preceded by a dysregulation of the actomyosin cytoskeleton. Similar to the *DN-myl9* (Figure 3—figure supplement 2A, B, and D) and *myh6* (Figure 5—figure supplement 1A-D) phenotypes, the overall levels of F-actin in the *tbx5a* mutant ventricular myocardium are reduced at 36 hpf, whereas levels of pMyosin are comparable to wild-type (Figure 6—figure supplement 1A-D). In addition to an overall deficit in F-actin, we found mislocalization of subcellular F-actin and pMyosin, and this was particularly prominent in the proximal OC (Figure 6I-P). In the *tbx5a* mutant proximal OC, F-actin and pMyosin enrichment shift away from the basal surface and accumulate more at the lateral and apical surfaces (Figure 6O and P; Figure 6—figure supplement 1E). In distal OC and in IC cells, F-actin is also shifted laterally and apically (albeit to a lesser extent than in the proximal OC), but the ratio of basal to apical/lateral pMyosin is comparable to wild-type (Figure 6O and P; Figure 6—figure supplement 1E).

Intriguingly, in contrast to the *myh6* mutant scenario where the effect on pMyosin localization is very similar to the effect on F-actin localization (Figure 5O and P; Figure 5—figure supplement 1E), the F-actin phenotype observed in *tbx5a* mutants is more severe than the pMyosin phenotype (Figure 6O and P; Figure 6—figure supplement 1E). Overall, these data suggest that Tbx5a-regulated pathways are particularly important for the enrichment of F-actin at the basal surface of OC cells.

Given the actomyosin and cell morphology phenotypes in flow-deficient *myh6* mutants (Figure 5) and the reduced heart rate in *tbx5a* mutants (Garrity et al., 2002), we wondered whether the aberrant effects on F-actin localization in *tbx5a* mutants reflect indirect consequences of altered blood flow or a cell-intrinsic loss of *tbx5a* function. To address this, we took a mosaic approach in which we transplanted wild-type blastomeres into *tbx5a* mutant host embryos and *tbx5a* mutant blastomeres into wild-type host embryos at ∼3 hpf, as well as performing wild-type into wild-type and mutant into mutant controls (Figure 7A-D). We fixed host embryos at 36 hpf and assessed amounts of F-actin at basal, lateral, and apical membranes (Figure 7E-P; Figure 7—figure supplement 1). In a *tbx5a* into wild-type scenario, we found that the F-actin organization in *tbx5a* mutant donor-derived proximal OC cells largely resembled that of their wild-type neighbors (Figure 7K and M; Figure 7—figure supplement 1C), indicating that something about the wild-type environment can rescue this aspect of the *tbx5a* mutant phenotype. In light of this result, we were particularly intrigued by the wild-type into *tbx5a* scenario. Here, we found that wild-type donor-derived cells in a *tbx5a* mutant proximal OC contained a significantly higher proportion of basal F-actin compared with apical and lateral F-actin, when compared with their *tbx5a* mutant neighbors (Figure 7H and J; Figure 7—figure supplement 1B). This suggests an ability of wild-type cells to organize their F-actin network regardless of the *tbx5a* status of surrounding cells. Together, these data indicate that there is at least a partially cell-autonomous role for *tbx5a* in establishing the divergent actomyosin organization observed in the developing curvatures, but that there are also likely to be non-autonomous ways that *tbx5a* influences actomyosin organization.

**Figure 7.**
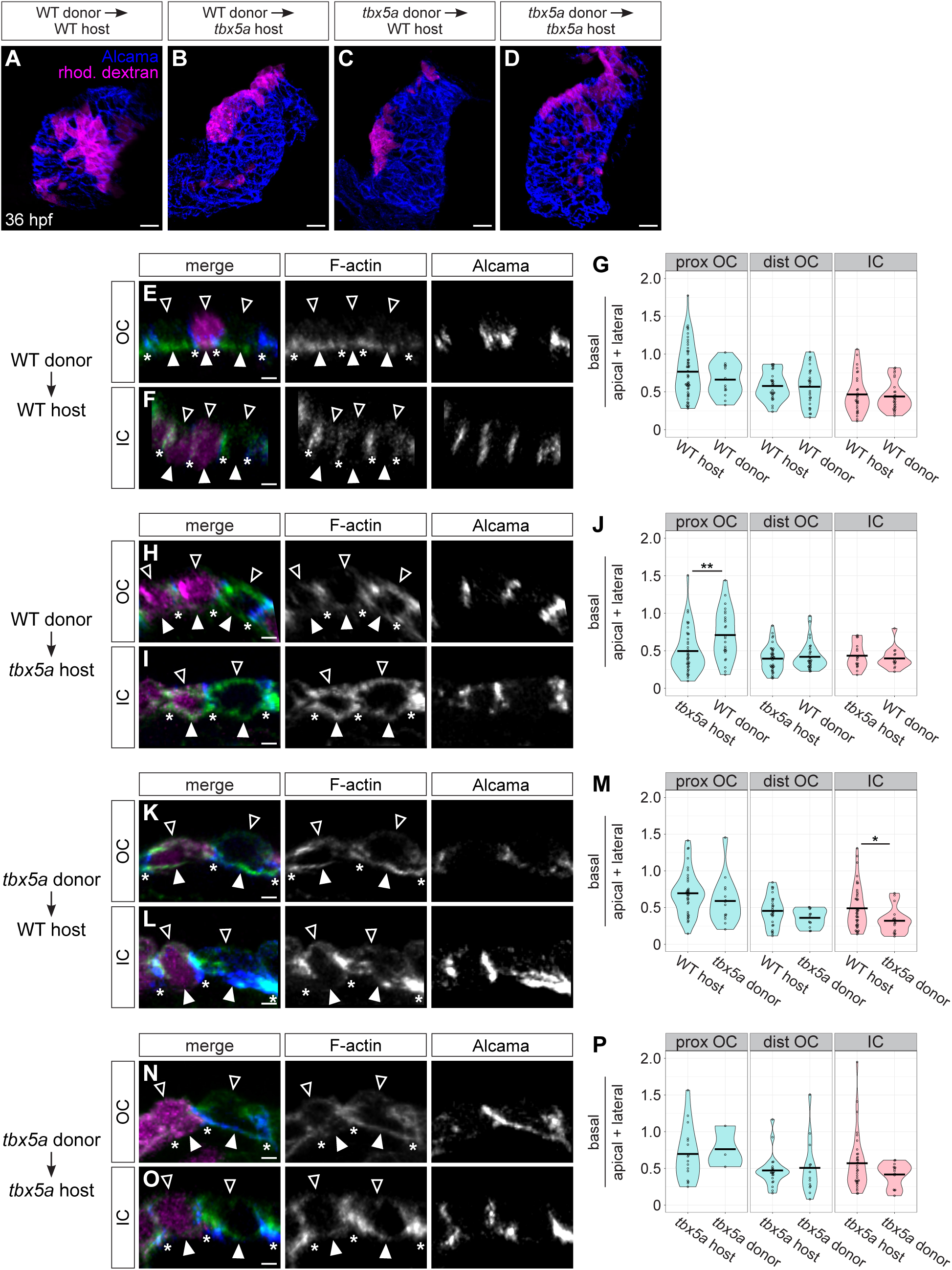
*tbx5a* functions in a partially cell-autonomous manner to support subcellular F-actin organization. (**A-D**) 3D reconstructions show examples of mosaic 36 hpf hearts resulting from blastomere transplantation. Immunostaining for Alcama labels lateral membranes of cardiomyocytes (blue), and donor-derived cells are labeled with rhodamine-dextran (magenta). (**E**, **F**, **H**, **I**, **K**, **L**, **N**, **and O**) Cross-sections through representative cardiomyocytes from the OC (E, H, K, and N) and IC (F, I, L, and O) from each of the four transplant scenarios. Immunostaining for Alcama labels lateral membranes of cardiomyocytes (blue) and staining with Phalloidin labels F-actin (green); rhodamine dextran labels donor-derived cardiomyocytes (magenta). Empty arrowheads: apical membranes. Filled arrowheads: basal membranes. Asterisks: basal extreme of lateral membranes. (**G**, **J**, **M**, **and P**) Violin plots compare calculated values of (mean basal F-actin / (mean apical F-actin + mean lateral F-actin)) of host-derived cardiomyocytes to those of donor-derived cardiomyocytes for each transplant scenario. Each dot represents an individual cell. * denotes p < 0.05, and ** denotes p < 0.01, Wilcoxon test. For WT into WT transplants: host proximal OC (N=6 embryos, n=55 cells); donor proximal OC (N=7 embryos, n=14 cells); host distal OC (N=5 embryos, n=30 cells); donor distal OC (N=6 embryos, n=29 cells); host IC (N=4 embryos, n=30 cells); donor IC (N=4 embryos, n=29 cells). For WT into *tbx5a* transplants: host proximal OC (N=5 embryos, n=40 cells); donor proximal OC (N=5 embryos, n=25 cells); host distal OC (N=5 embryos, n=40 cells); donor distal OC (N=5 embryos, n=35 cells); host IC (N=5 embryos, n=18 cells); donor IC (N=5 embryos, n=14 cells). For *tbx5a* into WT transplants: host proximal OC (N=7 embryos, n=38 cells); donor proximal OC (N=5 embryos, n=14 cells); host distal OC (N=6 embryos, n=28 cells); donor distal OC (N=5 embryos, n=13 cells); host IC (N=5 embryos, n=44 cells); donor IC (N=5 embryos, n=18 cells). For *tbx5a* into *tbx5a* transplants: host proximal OC (N=4 embryos, n=16 cells); donor proximal OC (N=2 embryos, n=3 cells); host distal OC (N=4 embryos, n=21 cells); donor distal OC (N=4 embryos, n=14 cells); host IC (N=5 embryos, n=36 cells); donor IC (N=5 embryos, n=14 cells). Scale bars = 20 μm (A-D); 3 μm (E, F, H, I, K, L, N, and O).

We next wanted to know whether *tbx5a* also plays a cell-autonomous role in promoting the divergence of OC and IC cardiomyocyte morphologies, or if the cardiomyocyte shape anomalies observed in Figure 6A-H are due to cardiac function defects present in *tbx5a* mutants (Garrity et al., 2002). To distinguish between these possibilities, we used the same experimental scheme as in Figure 7, but we instead fixed host embryos at 48 hpf and assessed cardiomyocyte shapes (Figure 7—figure supplement 2). Here, we focused on assessing apical surface area, as our data suggest a primary role for actomyosin in promoting the planar expansion of cardiomyocytes, particularly in the OC. Interestingly, consistent with our analysis of actomyosin localization in wild-type into *tbx5a* mosaic hearts (Figure 7H and J), wild-type donor-derived OC cells extended in the planar axis significantly more than their *tbx5a* mutant host-derived neighbors, allowing them to assume a more squamous identity (Figure 7—figure supplement 2H and J). Also like our mosaic analysis of actomyosin localization (Figure 7K and M), *tbx5a* mutant donor-derived cells that resided in a wild-type OC largely exhibited morphology similar to their wild-type neighbors (Figure 7—figure supplement 2K and M). Finally, donor-derived IC cells in both wild-type into mutant and mutant into wild-type scenarios achieved morphologies similar to their host-derived neighbors (Figure 7—figure supplement 2I-J and L-M). Taking our transplantation experiments together, it appears that the environment of a wild-type heart is able to largely override the absence of *tbx5a* function in donor-derived mutant cardiomyocytes, but, conversely, wild-type cells are capable of achieving stereotypical OC traits even in a *tbx5a* mutant background. This suggests that the ventricular phenotypes we see in *tbx5a* mutants are not solely due to defects in blood flow and that there is some cell-autonomous aspect to *tbx5a* function during curvature formation.

An unexpected finding from our mosaic studies was that, in some cases, donor-derived cells sent basal projections underneath their host-derived neighbors (Figure 7—figure supplement 3). We observed such projections in all four of our transplant scenarios; these projections were typically rather short and slim and were usually restricted to the junctions between neighboring cells (Figure 7—figure supplement 3C, D, and I). However, in the wild-type into *tbx5a* mutant transplant scenario, it was not uncommon for wild-type donor-derived OC cells to send long, broad projections underneath the cell body of a neighboring *tbx5a* mutant host-derived cell (Figure 7—figure supplement 3E-I). This could be a reflection of the comparatively robust ability of wild-type cells to expand their basal domain, a relatively reduced cell-extracellular matrix (ECM) connection in *tbx5a* mutant tissue, or both. This idea is further supported by the observation that, in the same wild-type into *tbx5a* mutant scenario, donor-derived OC cells that did not contact any other donor-derived OC cells (*i.e.* those in single-cell clones) had, on average, a higher apical surface area than donor-derived OC cells that contacted other donor-derived cells (Figure 7—figure supplement 4A). Further morphological assessment of these single-cell clones showed that all of them sent out basal projections: 3 of the 7 clones sent thin basal projections that extended between the junctions of neighboring host cells (Figure 7—figure supplement 4B and C), and the remaining 4 sent larger basal projections underneath neighboring host-derived cells (Figure 7—figure supplement 4D and E). We propose that this ability to send basal projections is a mechanism that normally drives planar expansion of OC cells, and that this activity is increased when neighboring cells maintain a looser attachment to the underlying ECM. Additionally, these data further support a cell-autonomous role for *tbx5a* in directing cardiomyocyte behavior.

## Discussion

Through analysis of the cellular and subcellular dynamics of individual cardiomyocytes within the developing ventricle, we have gained several new insights into the mechanisms that drive the morphological divergence of OC and IC cardiomyocytes and the potential contribution of this divergence to chamber curvature formation. Although cardiomyocytes in both curvatures increase in volume during curvature emergence, cells in the OC expand preferentially in the planar axis, whereas cells in the IC expand preferentially in the apicobasal axis. These contrasting behaviors are closely preceded by a divergence in subcellular actomyosin localization between the curvatures. Further, modulation of actomyosin activity affects the ability of OC cells to attain the expected squamous shape, indicating that the actomyosin network plays an important role in promoting squamous instead of cuboidal cardiomyocyte morphologies. Finally, the regionalized distinctions between the actomyosin landscapes and, subsequently, the cell morphologies in the curvatures are influenced by both extrinsic factors, such as blood flow through the ventricle, and intrinsic factors, like the transcription factor Tbx5a.

Taken together, our data suggest a novel model in which regional regulation of the actomyosin cytoskeleton mediates the planar expansion of OC cells, ultimately causing a divergence in OC and IC cell morphologies during a critical period of curvature formation (Figure 8). Prior to the onset of curvature formation, the largest distinctions in actomyosin organization within the ventricle are primarily proximal versus distal (Figure 2—figure supplement 1), in agreement with previous studies (Merks et al., 2018). As development proceeds, however, differences begin to emerge between the subcellular landscapes of OC and IC cells. Specifically, by 36 hpf cells throughout the proximodistal length of the IC acquire more apical actomyosin, whereas cells especially in the proximal OC maintain strong basal localization of actomyosin (Figure 2 and Figure 2—figure supplement 2). We propose that this basal enrichment of actomyosin promotes subsequent planar expansion of OC cardiomyocytes between 36 and 48 hpf, perhaps by engaging the focal adhesion machinery. The coincident lack of actomyosin at the apical surface of proximal OC cardiomyocytes may allow for low localized tension, ensuring that the apical surface can passively expand along with the actively expanding basal surface. In the IC, we speculate that the redistribution of actomyosin away from the basal surface reduces certain associations with the ECM and therefore reduces outward pushing forces; simultaneously, more actomyosin at the apical surface could play a more active and stereotypically contractile role in maintaining a compact apical surface area. Thus, we envision that the actomyosin landscapes in each curvature support OC cells in expanding outward along the planar axis and IC cells in remaining more constricted.

**Figure 8.**
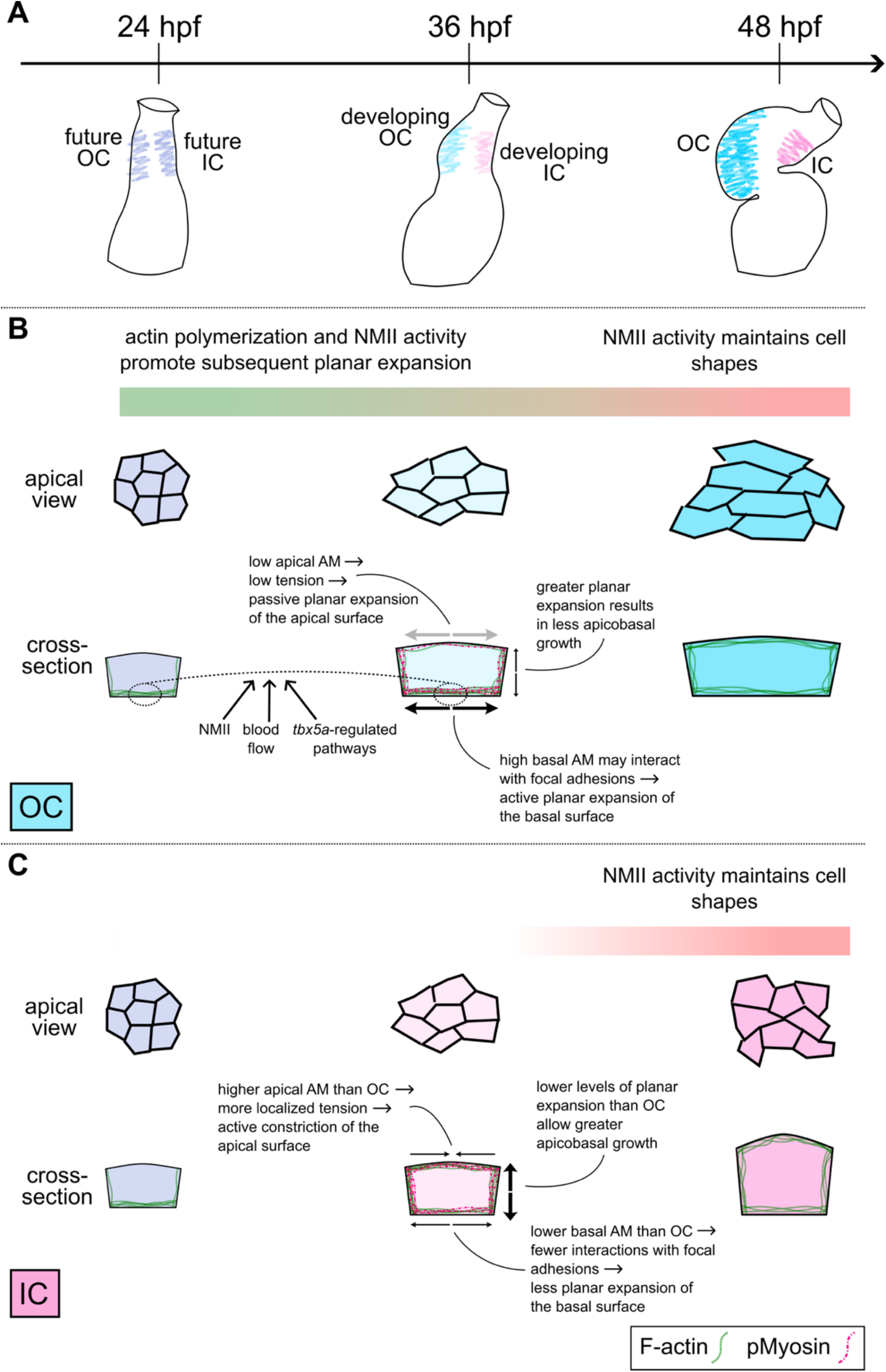
Changes to the actin cytoskeleton and cardiomyocyte shape during curvature formation. (**A**) During the second day of zebrafish development, the linear heart tube transforms into a two-chambered heart. Once similarly contoured regions (light purple) bend and stretch to become the convex OC (blue) and the concave IC (pink). (**B and C**) Schematic portrays select cellular and subcellular traits and events that accompany curvature formation. At the linear heart tube stage, cardiomyocytes in the future OC and IC (particularly those positioned more proximally, as shown here; light purple) present similar morphologies and F-actin organization, with most F-actin restricted to the basal surface. Over the next 12 hours, OC (B, light blue) and IC (C, light pink) cardiomyocytes have begun to diverge slightly in shape, and F-actin organization has also diverged. In the OC, there remains a large pool of basal F-actin, whereas in the IC, a lower proportion of F-actin resides at the basal surface and there instead exists a larger pool of apical and lateral F-actin. This difference is facilitated by NMII activity, as well as by blood flow through the ventricle and by *tbx5a*-regulated pathways; specifically, these factors appear to influence either the retention of basal F-actin or the prevention of formation of F-actin networks at the lateral and apical surfaces in OC cardiomyocytes. We propose that the basal enrichment of F-actin in OC cardiomyocytes (B) allows for interaction with focal adhesions to enact outward pushing of the basal surface (thick black arrows). Simultaneously, maintaining a low amount of actomyosin at the apical surface may allow this surface to passively expand along with the actively spreading basal surface (grey arrows). In the IC (C), we speculate that lower levels of basal F-actin might result in weakened interaction with focal adhesions and less outward pushing (thin black arrows), while more actomyosin at the apical surface may increase tension and prevent passive spreading (inward-facing thin black arrows). As a consequence of limited planar expansion, the increasing volume of IC cardiomyocytes instead leads to expansion along the apicobasal axis (thick black arrows). By 48 hpf, OC (darker blue) and IC (darker pink) cells have acquired strikingly different cell morphologies. In contrast, the actin cytoskeleton seems to have converged into similar arrangements in both OC and IC cardiomyocytes, with F-actin distributed fairly equally around all cell membranes, and, in both regions, it seems that the role of actomyosin changes altogether, with the primary role of NMII being to maintain cell shapes.

This model of contrasting OC and IC cell behaviors is also supported by aspects of our mosaic analyses. For example, we found that wild-type cells in the midst of a *tbx5a* mutant OC often send large, broad projections underneath neighboring mutant cells, whereas wild-type cells in a wild-type OC send much smaller projections that are usually restricted to cell junctions (Figure 7—figure supplement 3 and Figure 7—figure supplement 4). This suggests that wild-type OC cells have a mechanism for sending projections, and that *tbx5a* mutant OC cells (which, at the earlier stage of 36 hpf, have a reduced pool of basal actomyosin (Figure 6 and Figure 6—figure supplement 1)) may have a weakened connection to the underlying ECM. In addition, we found that cells expressing *DN-myl9* (which also exhibit dysregulated actomyosin localization at 36 hpf (Figure 3—figure supplement 3)) do not undergo proper planar expansion when surrounded by either wild-type cells (Figure 4) or other *DN-myl9*-expressing cells (Figure 3).

These data suggest that cells with reduced actomyosin activity or disrupted F-actin organization are not capable of the outward pushing behavior that we propose enables planar expansion in the OC. Intriguingly, recent work has identified actin-based protrusions that drive the elongation of zebrafish atrial cardiomyocytes as the atrial wall thickens after 72 hpf (Albu et al., 2024), and we are keen for our future work to explore whether similar protrusions promote planar expansion in the ventricular OC. Another intriguing possibility is that the OC cardiomyocyte projections could help to advance the cellular intercalations involved in the twisting of the heart tube; consistent with this idea, *tbx5a* mutant embryos exhibit reduced cardiomyocyte intercalation and cardiac torsion (Tessadori et al., 2021).

Although our model of outward pushing contrasts with the frequent demonstration that actomyosin constricts cell boundaries during morphogenesis, the idea that a growing actin cytoskeleton could force cardiomyocyte membranes outward during curvature formation was originally put forth by Latacha and colleagues nearly two decades ago (Latacha et al., 2005). Additionally, we know that this effect of the actin cytoskeleton is necessary in other contexts. For example, in the leading edge of migrating cells, actin branching, polymerization, and depolymerization pushes the membrane outward, specifically in the direction of movement (Bisi et al., 2013; Innocenti, 2018; Schaks et al., 2019). These pushing forces are also observed in epithelial contexts: in the *Xenopus* epidermis, cell-autonomous actin dynamics at the apical surface drive outward pushing and thus apical emergence of multiciliated cells (Sedzinski et al., 2016, 2017), and, in *Drosophila* follicle cells, actomyosin and its association with the ECM promotes expansion of the basal surface, resulting in cell flattening and tissue elongation (Li et al., 2024; Santa-Cruz Mateos et al., 2020). It would be valuable for future studies to explore potential mechanistic similarities between these contexts and the planar spread of OC cardiomyocytes.

Our data clearly indicate that both extrinsic and intrinsic factors contribute to the regulation of the divergent cytoskeletal landscapes observed in the ventricular curvatures. For the former, we demonstrated that mutants with reduced blood flow through the ventricle not only fail to acquire the expected OC cell morphologies, as previously reported (Auman et al., 2007), but also exhibit earlier defects in the OC actomyosin network (Figure 5). Specifically, in *myh6* mutants, F-actin levels are dramatically reduced throughout the ventricular myocardium (Figure 5—figure supplement 1) and both F-actin and pMyosin are shifted away from the basal membrane to the lateral and apical membranes in OC cells (Figure 5). For the latter, we found that *tbx5a* mutants, already appreciated for their underformed curvatures (Garrity et al., 2002; Tessadori et al., 2021), also exhibit reduced F-actin levels (Figure 6—figure supplement 1), shifted F-actin away from the basal membrane in OC cells (Figure 6), and defects in the planar expansion of OC cells (Figure 6). It is interesting to note that reduced blood flow and loss of *tbx5a* function both primarily impacted cells in the OC; although both curvatures are exposed to blood flow and *tbx5a* is expressed throughout the ventricular myocardium (Garrity et al., 2002), these factors do not influence the cytoskeleton in IC cells to the same extent that they do in OC cells. This intimates the existence of yet unidentified curvature-specific factors that could help translate extrinsic and intrinsic influences into cytoskeletal changes. Regarding blood flow, this biomechanical input may be transmitted to the myocardial cytoskeleton through strain on the chamber or through myocardial-endocardial signaling, but it is unclear what molecules mediate this conversation. It will therefore be beneficial to understand the regional differences in gene expression that exist in the OC and IC, both before and during curvature formation. More specifically, it will be valuable for future work to identify and investigate a roster of actomyosin modifiers whose expression or activity may be regulated, even indirectly, by blood flow or by *tbx5a*. One interesting candidate is *adducin3a* (*add3a*), which is regulated by a heartbeat-dependent microRNA, *miR-143*, and encodes an F-actin capping protein that stabilizes actin polymers (Deacon et al., 2010; Matsuoka et al., 2000; Miyasaka et al., 2011). Overexpression of *add3a* has been shown to prevent planar expansion of OC cells, likely due to the hyperstabilization of the actin cytoskeleton (Deacon et al., 2010), suggesting that regionalized regulation of Add3a activity could contribute to the normal acquisition of OC cell morphology. Finally, it will be interesting to investigate whether any TBX5 targets that may mediate myosin phosphorylation in the mouse second heart field epithelium (Guijarro et al., 2024) also function downstream of Tbx5a to modify actomyosin in the OC.

Although actin dynamics have long been implied as a driver of chamber curvature formation, the role of NMII has been debated, with some studies dismissing its role and others supporting it. For example, chick embryos or explanted hearts treated with inhibitors of NMII just after the initiation of curvature formation go on to form relatively normal curvatures (Rémond et al., 2006). By contrast, explanted zebrafish heart tubes treated with Blebbistatin fail to form chamber curvatures (Noël et al., 2013). Of course, these disparate findings could simply signify differences between species. However, our studies offer an alternative explanation in which there are at least two distinct phases of the influence of actomyosin on the regulation of cardiomyocyte shape. Our treatments with Blebbistatin or Latrunculin B (Figure 3—figure supplement 1) imply that, early in curvature formation (∼24-36 hpf), both actin polymerization and NMII activity are required to allow the subsequent planar expansion of OC cells. In contrast, later in curvature formation (∼36-48 hpf), actin polymerization does not seem to impact OC cell planar expansion, and NMII even plays the opposing role of restricting planar spread, presumably due to its canonical contractile properties. These findings, together with the impact of DN-Myl9 and CA-Myl9 activity on both total F-actin (Figure 3—figure supplement 2) and subcellular localization (Figure 3—figure supplement 3), support a model wherein the early F-actin dynamics that are crucial for subsequent OC cell shape change may be regulated by NMII.

Indeed, the influence of NMII on F-actin dynamics is a commonly reported phenomenon (Bridgman et al., 2001; Cai et al., 2010; Huang et al., 2021; Laevsky & Knecht, 2003; Matsui et al., 2011; Murthy & Wadsworth, 2005; Wilson et al., 2010; Yamashiro et al., 2018; Zhou & Wang, 2008), and we therefore recommend special attention to the timing of any such actomyosin manipulations in studies of curvature formation. The perduring activity of DN-Myl9 and CA-Myl9 throughout curvature formation in our experiments may cloud our interpretations, as their aberrant activity during later curvature formation could suppress the effects of their earlier activity (and *vice versa*). In future studies, having both spatial and temporal control over actomyosin cytoskeleton modification will help to test these ideas more directly.

Cardiac looping involves the coordination of several morphogenetic events in both the ventricle and the atrium, as well as in structures like the OFT and AVC. However, the degree of interdependence of events in these different domains remains unclear. *Ex vivo* studies in zebrafish suggest that accretion of second heart field cells to the outflow and inflow tract regions are not required for the repositioning of chambers, torsion, or formation of the curvatures (Noël et al., 2013; Tessadori et al., 2021), implying that these processes exhibit some level of independence. In contrast, modulation of Bmp signaling in the AVC and IC leads to reduced chamber displacement, including a fairly unkinked IC (Lombardo et al., 2019), highlighting a link between AVC and IC morphogenesis. Our studies show that it is primarily the OC that is influenced by blood flow and *tbx5a* (Figures 5 and 6) and that the cell-autonomous effects of *tbx5a* and of *DN-myl9* expression are only observed in the OC (Figure 6—figure supplement 3 and Figure 4, respectively), suggesting that the pathways studied here are mostly relevant to OC development. Taking these data together, we speculate that the shape of the IC is largely coupled to morphogenesis of the AVC, whereas OC dynamics and shape are at least partially driven by the cell-intrinsic, actomyosin-based mechanisms presented here. However, we cannot rule out the possibility of interplay between OC and IC morphogenesis or the potential interaction of OC formation with other elements of looping, such as torsion and chamber displacement.

Furthermore, although our data highlight distinct actomyosin landscapes in the proximal and distal regions of the OC at 36 hpf, cells in similar regions at 48 hpf attain quite similar morphological attributes. In addition, disrupting actomyosin function affects cardiomyocyte shape change in both proximal and distal OC regions. We therefore speculate that different mechanisms may drive the cell and tissue shape changes in these regions, although both likely utilize actomyosin to some degree. We posit that the future use of methods that alter gene function in larger but distinct territories – such as throughout the entire or partial OC – will help to answer many of these questions.

Altogether, our work supports the notions that intrinsic regulators and extrinsic inputs converge to control the actomyosin cytoskeleton in the OC, that these events are crucial for changes to cell morphology, and that these changes ultimately influence the proper sculpting of the ventricle. These findings offer a mechanistic framework both for the specific circumstance of cardiac chamber emergence and for tissue curvature formation in general. In addition, they provide a foundation for further articulation of the pathways that create the distinctions between the OC and IC, which could ultimately shed light on the origins of certain types of congenital defects in chamber morphology and could also enhance efforts to engineer ventricular tissue *in vitro*.

## Materials and Methods

### Zebrafish

The following transgenic lines were used in this study: *Tg(myl7:WT-myl9-mScarlet)^bns524Tg^* (Priya et al., 2020), *Tg(myl7:DN-myl9-eGFP)^bns333Tg^*(Priya et al., 2020), *Tg(myl7:CA-myl9-eGFP)^bns332Tg^* (Priya et al., 2020), *Tg(myl7:eGFP-Hsa.HRAS)^s883Tg^* (D’Amico et al., 2007), and *Tg(myl7:mKate-CAAX)^sd11Tg^* (Lin et al., 2012). We also used lines carrying the *tbx5a^m21^* (Garrity et al., 2002) and *myh6^m58^*(Berdougo et al., 2003) mutations. Heterozygous carriers of mutations were identified by PCR amplification and subsequent restriction enzyme digest, as previously described for *tbx5a^m21^* (Auman et al., 2024) and *myh6^m58^* (Lin et al., 2012). Homozygous mutant embryos were identified by previously characterized morphological phenotypes (Berdougo et al., 2003; Garrity et al., 2002). All work presented here followed protocols approved by the Institutional Animal Care and Use Committee at the University of California, San Diego.

### Pharmacological treatments

Embryos were dechorionated and placed in 6-well culture dishes and exposed to either 1.25% DMSO, 1.25% DMSO + 100 ng/mL Latrunculin B (Millipore Sigma, L5228), or 1.25% DMSO + 30 μM Blebbistatin (Millipore Sigma, B0560) in E3 medium. DMSO controls were kept in solution between 24-48 hpf. Latrunculin B and Blebbistatin treated embryos were either kept in drug-containing media between 24-36 hpf and switched to 1.25% DMSO between 36-48 hpf, or *vice versa*. At 48 hpf, all embryos were fixed and processed as described below.

### Immunofluorescence and phalloidin staining

Embryos were dechorionated, fixed in 1% methanol-free formaldehyde (ThermoFisher Scientific, 28908) for 1 hour and 10 minutes with gentle rocking, and then rinsed three times in PBT (1X PBS containing 0.1% Tween 20 (Sigma-Aldrich, P9416)). Embryos were rocked rapidly in a 0.2% saponin solution (Sigma-Aldrich, S4521) in PBT containing 0.5% Triton X-100 (G-Biosciences, 786513) until yolks were dissolved. This took 20-35 minutes and was monitored under a dissecting microscope. Embryos were rinsed three times in PBT, placed in 4% formaldehyde, and incubated at 4°C overnight. On the following day, to improve antibody signal-to-noise ratio, embryos were rinsed three times in PBT, placed in prechilled 100% acetone, and incubated at −20°C for 8 minutes. Acetone was replaced with PBT + 0.5% Triton X-100 for one hour at room temperature. To ensure reagent access to the heart, forceps were used to open the pericardial cavity. Embryos were then blocked with 2 mg/ml bovine serum albumin (Sigma-Aldrich, A9647) and 10% goat serum in PBT for at least one hour at room temperature, then incubated with primary antibody in block overnight at 4°C. Embryos were then washed extensively with PBT, blocked as described for primary antibody incubation, and incubated with the secondary antibodies in blocking solution overnight at 4°C. Embryos were then washed extensively with PBT.

The following primary antibodies were used at the specified dilutions: mouse anti-Alcama (Developmental Studies Hybridoma Bank, Zn-8 supernatant, 1:50); rabbit anti-Cdh2 (GeneTex, GTX125885, 1:200); rabbit anti-phospho-Myosin (Abcam, ab2480, 1:100); rabbit (Life Technologies, A11122, 1:500) or chicken (Life Technologies, A10262, 1:1000) anti-GFP; rabbit anti-dsRed (also detects mScarlet; Clontech, 632496, 1:1000); rabbit anti-TagRFP (also detects mKate; Evrogen, AB233, 1:500); mouse anti-myosin heavy chain (Developmental Studies Hybridoma Bank, MF20 supernatant, 1:50); and mouse anti-Myh6 (Developmental Studies Hybridoma Bank, S46 supernatant, 1:50).

All secondary antibodies were produced in goat by ThermoFisher Scientific and used at a dilution of 1:300. Secondary antibodies used were: anti-mouse-AlexaFluor 488 (A11001); anti-mouse-AlexaFluor 568 (A11031); anti-mouse-AlexaFluor 647 (A21236); anti-rabbit AlexaFluor 488 (A11008); anti-rabbit AlexaFluor 568 (A11011); anti-rabbit AlexaFluor 647 (A21245); and anti-chicken AlexaFluor 488 (A11039). In some cases, Phalloidin-AlexaFluor 488 or Rhodamine-Phalloidin (Life Technologies, A12379 or R415; each reconstituted to 200U/mL in MeOH) was added to the secondary antibody solution at a dilution of 1:30.

### Fluorescent *in situ* hybridization

Fluorescent *in situ* hybridization was performed as previously described (Leerberg et al., 2017), but embryos were fixed and deyolked as for immunofluorescence before MeOH dehydration and were incubated with Proteinase K (50 μg/mL) in PBT for 2 minutes at room temperature prior to hybridization with probe. Probe for *nppa* (ZDB-GENE-030131-95) was produced as previously described (Berdougo et al., 2003). Probe for *mb* (ZDB-GENE-040426-1430) was produced by PCR amplification of 48 hpf embryonic cDNA followed by T7 *in vitro* transcription. The forward primer used was 5′-GGACAAACACCGCGACAGAC-3′ and the reverse primer used was: 5′-<u>TAATACGACTCACTATAGGG</u>CCTGAGACCCTAACGAACCATTAT-3′; underlined portion is the T7 binding site.

### Transplantation

Blastomere transplantation was performed similarly to a previous description (Auman et al., 2007). Briefly, donor embryos were injected with 5% lysine-fixable rhodamine dextran (ThermoFisher Scientific, D1817) in 0.2M KCl at the one-cell stage. Approximately 10-30 blastomeres were taken from dechorionated 3-4 hpf donor embryos and placed into the margin of similarly staged host embryos. Host embryos were raised in E3 medium containing penicillin-streptomycin (Millipore Sigma, P4458; diluted 1:50), in agarose-coated wells. Host embryos were allowed to develop until 36 or 48 hpf when they were screened for the presence of donor-derived cells in the myocardium, which could be identified by fluorescence under a Zeiss Axiozoom microscope. Hosts containing donor-derived cardiomyocytes were fixed and processed for immunostaining as described above. In order to be included for analysis, cardiomyocytes needed to meet specific criteria. For donor-derived cells, we analyzed any cell with at least half of its body within the OC or IC boundary. For host-derived cells, we analyzed any cell (1) in direct contact with a donor-derived cell or one cell-distance from a donor-derived cell and (2) with at least half of its body within the OC or IC boundary.

For *Tg(myl7:CA-myl9-eGFP)* and *Tg(myl7:DN-myl9-eGFP)* transplants, transgenic animals were crossed to wild-type animals, and the resulting embryos served as donors; wild-type embryos served as hosts. For the *tbx5a* transplantation experiments, animals heterozygous for the *tbx5a* mutation were crossed to obtain both donor and host embryos. After transplantations were complete, donor embryos were genotyped by PCR and restriction enzyme digest for the *tbx5a* mutation, as described above. Genotypes of host embryos were determined based on the presence or absence of pectoral fin buds at 36 hpf or pectoral fins at 48 hpf (Garrity et al., 2002). While it is possible that a large donor-derived clone could affect the development of one pectoral fin bud, it is unlikely, given the number of donor cells transplanted, that this would impact both pectoral fin fields in the same host. Both sides were therefore assessed for the presence of pectoral fin buds/fins, and only those host embryos that had matching sides were analyzed.

### Image acquisition

After all immunostaining and fluorescent in situ hybridization procedures, hearts were dissected from embryos under a Zeiss AxioZoom microscope. To keep organ morphology as intact as possible, hearts were simply mounted in a droplet of PBT on a cover slip and immediately imaged using a Leica SP8 laser-scanning confocal microscope and a 25X water-immersion objective. In all experiments where signal intensity was measured, laser settings were kept consistent between samples. Z-stacks were captured with a slice thickness of 0.3 μm.

### Image analysis

All image files captured by the Leica SP8 confocal were rendered as 3D reconstructions in Imaris (Bitplane). Often, secondary analysis was completed in FIJI (ImageJ).

#### Defining the ventricular OC and IC

To develop a method to standardize the boundaries of the OC and IC between 48 hpf samples (Figure 1—figure supplement 1C), we first assessed the expression domain of *nppa* in 12 wild-type hearts at 48 hpf (Figure 1—figure supplement 1A). By using the following measurements, we found that we could capture, on average, the highest *nppa* signal in the resulting “OC” and, on average, the lowest *nppa* signal in the resulting “IC”. These territories also corresponded well to the regions with the lowest (OC) and highest (IC) *mb* expression (Figure 1—figure supplement 1B). These measurements were then applied to 48 hpf hearts throughout this study.

The following abbreviations are used in our description of this measurement method: PB_OC_: OC proximal boundary; DB_OC_: OC distal boundary; DB_OFT(OC)_: distal boundary of the OC side of the OFT; PB_IC_: IC proximal boundary; DB_IC_: IC distal boundary; DB_OFT(IC)_: distal boundary of the IC side of the OFT.

To delineate the OC in Imaris (Figure 1—figure supplement 1C), the arc of the ventricular OC was measured beginning one cell diameter from the AVC (the PB_OC_) to the DB_OFT(OC)_. This arc is represented by a blue dotted line in Figure 1—figure supplement 1C. Starting from the PB_OC_, the DB_OC_ was marked at 2/3 of the distance from the PB_OC_ to the DB_OFT(OC)_. An *Oblique Slicer* was placed between the PB_OC_ and DB_OC_, ensuring that it was perpendicular to the arc of the OC. The position of the *Oblique Splicer* is represented by a solid blue line in Figure 1—figure supplement 1C. The portion of tissue bounded by the solid blue line was considered the OC.

To delineate the IC in Imaris (Figure 1—figure supplement 1C), the arc of the ventricular IC was measured from the AVC (the PB_IC_) to the DB_OFT(IC)_. This arc is represented by a dotted pink line in Figure 1—figure supplement 1C. Point I (red “I” in Figure 1—figure supplement 1C) was marked at 1/2 of the distance between PB_OC_ and DB_OC_. Point II (red “II” in Figure 1—figure supplement 1C) was marked at 1/3 of the distance between PB_IC_ and Point I. The proximal border of the IC was considered to run from the PB_IC_ to Point II. This border is represented by a solid pink line in Figure 1—figure supplement 1C. Starting from the PB_IC_, the DB_IC_ was marked at 1/2 of the distance from the PB_IC_ to the DB_OFT(IC)_. Point III (red “III” in Figure 1—figure supplement 1C) was marked halfway between the DB_IC_ and DB_OC_. The distal border of the IC was considered to run from the DB_IC_ to Point III. This border is also represented by a solid pink line in Figure 1—figure supplement 1C. Finally, an *Oblique Slicer* was placed between Points II and III, ensuring that it was perpendicular to the arc of the IC. The position of this *Oblique Splicer* is represented by a solid pink line in Figure 1—figure supplement 1C. The portion of tissue bounded by the solid pink lines was considered the IC.

To account for the reduced contours of the OC and IC in hearts at 36-37 hpf (Figure 1—figure supplement 1D), slight adjustments were made in our measurement method. Specifically, the arc of the ventricular OC was measured beginning one cell diameter from the AVC (the PB_OC_) to the DB_OFT(OC)_. This arc is represented by a dotted blue line in Figure 1—figure supplement 1D. Starting from the PB_OC_, the DB_OC_ was marked at 2/3 of the distance from the PB_OC_ to the DB_OFT(OC)_. The arc of the ventricular IC was measured from the AVC (the PB_OC_) to the DB_OFT(IC)_. This arc is represented by a dotted pink line in Figure 1—figure supplement 1D. A line was drawn between the PB_OC_ and the PB_IC_, as well as between the DB_OC_ and the DB_IC_; these are represented by dotted purple lines in Figure 1—figure supplement 1D. Point I (red “I” in Figure 1—figure supplement 1D) was marked at 1/3 of the distance between PB_OC_ and PB_IC_, and Point II (red “II” in Figure 1—figure supplement 1D) was marked at 2/3 of the distance between PB_OC_ and PB_IC_. Point III (red “III” in Figure 1—figure supplement 1D) was marked at 1/3 of the distance between DB_OC_ and DB_IC_, and Point IV (red “IV” in Figure 1—figure supplement 1D) was marked at 2/3 of the distance between DB_OC_ and DB_IC_. The medial border of the OC was considered to run from Point I to Point III (solid blue line in Figure 1—figure supplement 1D), and the medial border of the IC was considered to run from Point II to Point IV (solid pink line in Figure 1—figure supplement 1D). As for 48 hpf hearts, *Oblique Slicers* were used in accordance with the medial border measurements in Figure 1—figure supplement 1D to bound the OC and IC.

#### Labeling lateral cell membranes

When the labeling of lateral membranes was necessary, we utilized one of four reagents: *Tg(myl7:mKate-CAAX)*, *Tg(myl7:Ras-eGFP)*, anti-Cdh2 antibody, or anti-Alcama antibody. The first two reagents label all membranes of the cardiomyocyte, whereas the latter two are cell adhesion molecules and are therefore expected to label only the lateral membranes. Given the possibility that Cdh2 and Alcama could be polarized along the apicobasal axis, we tested the extent to which these signals overlapped with *Tg(myl7:Ras-eGFP)* signal (Figure 1—figure supplement 2). The consistent overlap of Cdh2 and Alcama signal with lateral Ras-eGFP signal (Figure 1—figure supplement 2; (Cherian et al., 2016)) led us to consider both Cdh2 and Alcama as sufficient representations of the entire lateral membrane, and we therefore used antibodies against either, depending on the specific requirements of an experiment.

#### Cardiomyocyte morphometrics

For each cardiomyocyte in the OC and IC, Imaris was used to capture *Snapshots* of the necessary views of the cell, FIJI was used to take measurements, and Excel was used to compile these measurements into a .csv and to calculate apicobasal length and volume.

In Imaris: To capture the *en face* view to obtain apical surface area and circularity, a single *Snapshot* of the apical surface of the cardiomyocyte was taken, ensuring that the surface was flat and parallel to the screen. If a cardiomyocyte was large enough that it curved in the Z-plane, multiple *Snapshots* were captured. To do this, an *Oblique Slicer* (or more, if necessary) was placed where the greatest curvature existed, perpendicular to the long axis of the cell. Then two (or more, if necessary) *Snapshots* of the cell were taken — the first on one side of the *Oblique Slicer*, and a second on the other side of the *Oblique Slicer*, ensuring that each “side” of the cell was flat and parallel to the screen. To capture the cross-sectional view to obtain apicobasal length and volume measurements, two *Oblique Slicers* were placed while the cell was still in an *en face* view. These *Oblique Slicers* were perpendicular to the apical surface of the cell and approximately bisected the cell (one bisected the major axis and the other bisected the minor axis). The *Volume* of the 3D reconstruction was clicked *Off*, and the image was reoriented so that the “faces” of the *Oblique Slicers* were visible, showing cross-sections of the cell. The intersection of the two *Oblique Slicers* marked the current cell of interest. Two *Snapshots* of the cell were taken, one from each *Oblique Slicer*, ensuring that the *Slicers* were flat and parallel to the screen.

In FIJI: The scale was first set based on the scale bar provided by Imaris. To measure apical surface area and circularity, the *Freehand Selection* tool was used to trace the lateral membrane of the cardiomyocyte. If a cardiomyocyte was curved in the Z-plane and required two (or more, if necessary) *Snapshots*, the lateral membrane of one “side” of the cell was traced, stopping where the cell meets the *Oblique Slicer*. The selection was copied and pasted onto the *Snapshot* of the other “side”, the “sides” were positioned so that they joined together, and the border of the entire cell was traced. The *Measure* function was used to obtain *Area* and *Circularity* values. For each of the two cross-sectional *Snapshots* taken to measure apicobasal length, the basal width (the distance between the basal tips of the lateral membranes), apical width (the distance between the apical tips of the lateral membranes), and length of each lateral membrane (from basal tip to apical tip) were measured using the *Measure* function.

In Excel: To obtain apicobasal length, the average (mean) of the lengths of the four lateral membranes was calculated. To obtain volume, the average (mean) of the two basal width values and the two apical width values was calculated; then, the mean basal width, the mean apical width, and the mean apicobasal length were multiplied.

#### Subcellular actomyosin localization

Prior to image analysis, F-actin and pMyosin (where appropriate) signal in each ventricle was assessed in Imaris. Briefly, a *Mask* of the ventricular myocardium was generated, the *Mean* signal intensity for each channel was located in the *Statistics* tab, and these values were tested for outliers in R (R Core Team, 2024). Outliers were removed from further analysis.

For each cardiomyocyte in the OC and IC, Imaris was used to capture *Snapshots* of the necessary views of the cell, FIJI was used to take signal intensity measurements, and Excel was used to compile these measurements into a .csv and to calculate mean signal intensities.

In Imaris: Two cross-sectional views (Figure 2A and B) were captured as described above in the “*Cardiomyocyte morphometrics*” section, with the exception that a separate *Snapshot* was taken for each channel (for the membrane marker, for F-actin, and for pMyosin).

In FIJI: All images were converted to 8-bit and the channels for each *Snapshot* were *Merged*. Using the membrane marker channel as a guide, the *Freehand Line* tool, set to a width of ∼1 μm, was used to trace just inside the membrane. The *Mean* signal intensities of the F-actin signal and the pMyosin signal were measured along each membrane using the *Measure* function, toggling between channels as needed. Four *Mean* measurements were taken per *Snapshot*: basal, apical, and two lateral. To prevent the inclusion of signal from the lateral membranes into the basal/apical datasets and *vice versa*, a ∼1 μm segment in each “corner” (where the lateral and basal/apical membranes intersect) was avoided when performing the initial traces.

In Excel: For F-actin and for pMyosin (when present) in each cell, the averages (means) of the two basal signal intensities, of the two apical signal intensities, and of the four lateral signal intensities were calculated.

#### Total F-actin and pMyosin in the ventricular myocardium

All analysis was completed in Imaris. A *Surface* of the myocardium was generated, based on a channel that was only present in the myocardium and using a *Surfaces Detail* of 2 μm. Thresholds were utilized to ensure the surface encapsulated the ventricular myocardium but did not include excessive space around the myocardium. The *Surface* was *Cut* at the AVC/ventricle boundary and at the ventricle/OFT boundary. The *Mean* signal intensity of each channel was located in the *Statistics* tab. These intensities were reported in graphs as relative fluorescence units (rfu).

### Statistical analysis, plotting, and figure design

Statistics were performed in R (R Core Team, 2024) and RStudio (RStudio Team, 2024). For all comparisons, statistical significance was determined by the Wilcoxon test, which does not assume normal data distribution. Statistically significant p-values are indicated in graphs by asterisks as follows: * denotes p<0.05; ** denotes p<0.01; *** denotes p<0.001. All experiments were performed at least two independent times, and the exact number of embryos and/or cells examined for each experiment is reported in the corresponding figure legend.

The following R packages were used to restructure, statistically analyze, and plot the quantitative data captured from micrographs: dplyr (Wickham et al., 2023), ggbeeswarm (Clarke et al., 2023), ggplot2 (Wickham, 2016), ggpmisc (Aphalo, 2024), ggpubr (Kassambara, 2023), ggtext (Wilke & Wiernik, 2022), Lattice (Sarkar, 2008), plyr (Wickham, 2011), Tidyverse (Wickham et al., 2019), and ggnewscale (Campitelli, 2025). In Figures 1, 3, 5, 6, Figure 2—figure supplement 3, Figure 3—figure supplement 1, and Figure 3—figure supplement 3, SuperPlots (Lord et al., 2020) were employed to portray variation between samples within the same experimental group. Figures were produced in Adobe InDesign and Inkscape.

## Data availability

All data are available in the main text. Materials, Excel spreadsheets, and raw images are available upon request to the corresponding author.

## Acknowledgments

We thank members of the Yelon laboratory for valuable discussions, A. Negrete and M. Ayan for help with genotyping, and A. Yarbrough and the UCSD Animal Care Program for zebrafish care.

## Funding

American Heart Association grant 23TPA1072669 (DY)

Saving tiny Hearts Society (DY)

National Institutes of Health fellowship F32 HL147435 (DML)

American Heart Association fellowship 20POST35110077 (DML)

Max Planck Society (DYRS)

EMBO fellowship LTF1569 (RP)

Alexander von Humboldt Foundation fellowship (RP)

Cardio-Pulmonary Institute grant EXC 2026 project ID 390649896 (RP)

## Author contributions

Conceptualization: DML, DY

Data curation: DML, DY

Formal analysis: DML, GBA, DY

Methodology: DML, GBA, DY

Investigation: DML, GBA, DY

Resources: DML, RP, DYRS

Visualization: DML, DY

Validation: DML, DY

Supervision: DYRS, DY

Writing—original draft: DML

Writing—review & editing: DML, GBA, RP, DYRS, DY

## Supplementary Figures

**Figure 1—figure supplement 1.**
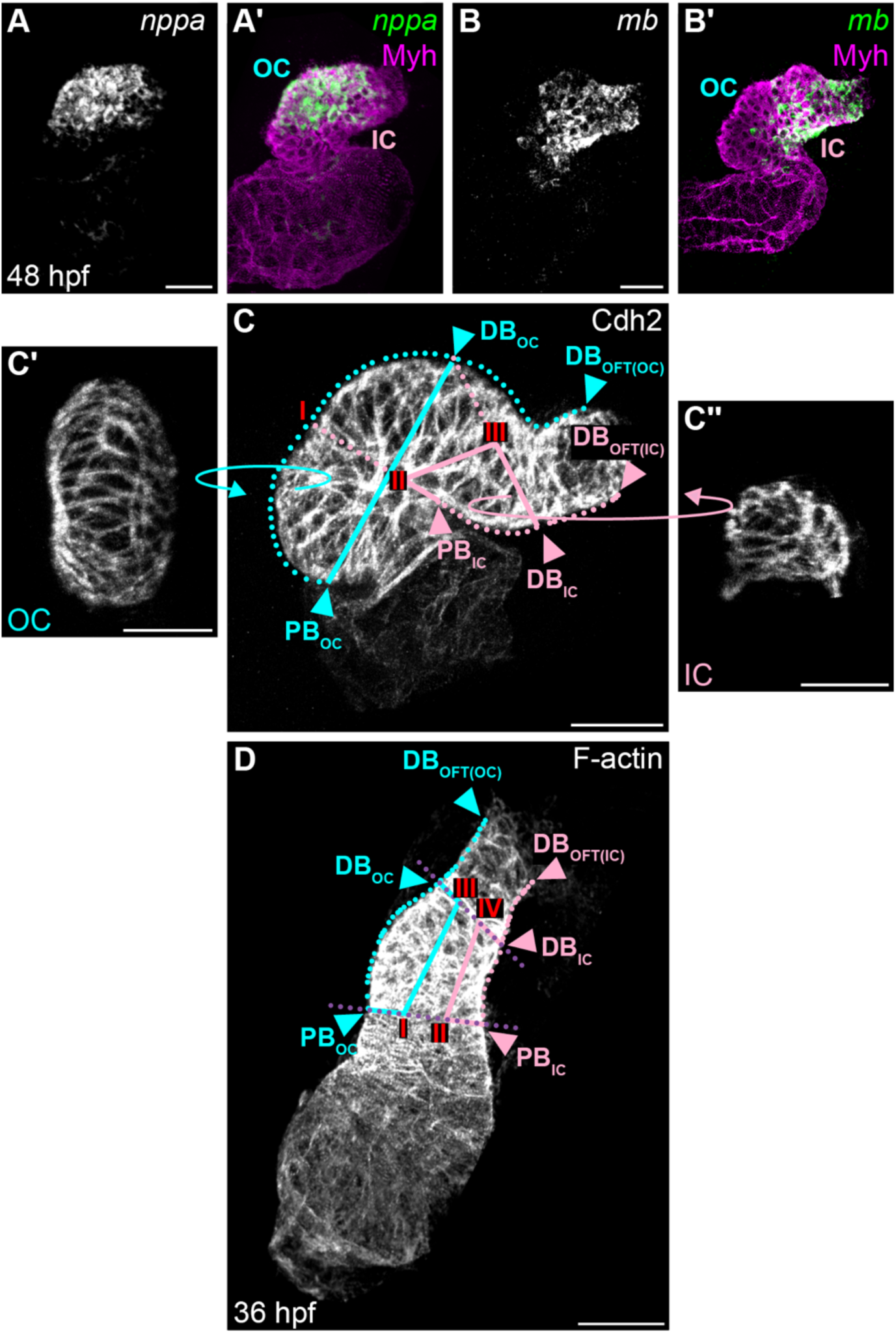
Defining the ventricular OC and IC. (**A and B**) Fluorescent *in situ* hybridization (FISH) for *nppa* (A) and *mb* (B) in 48 hpf wild-type hearts. (Aʹ and Bʹ) FISH signal is merged with immunostaining for Myosin heavy chain (Myh), outlining the entire myocardium. (A) *nppa* is enriched primarily in the OC of the ventricle; (B) *mb* is enriched in the ventricular IC and the outflow tract (OFT). (**C**) 3D reconstruction of a 48 hpf wild-type heart immunostained for Cdh2, which labels the lateral membranes of cardiomyocytes. Lines represent measurements used to segment the OC and IC (refer to Supplementary Materials and Methods for more detail, including information on the red “I”, “II”, and “III”). Once boundaries have been drawn in (C), the resulting OC (Cʹ) and IC (Cʹʹ) are swiveled to be viewed *en face*. (**D**) 3D reconstruction of a 36 hpf wild-type heart stained with Phalloidin, outlining F-actin distribution throughout the entire heart. Lines represent measurements used to delineate the OC and IC (refer to Materials and Methods for more detail, including information on the red “I”, “II”, “III”, and “IV”). PB_OC_: OC proximal boundary. DB_OC_: OC distal boundary. DB_OFT(OC)_: distal boundary of the OC side of the OFT. PB_IC_: IC proximal boundary. DB_IC_: IC distal boundary. DB_OFT(IC)_: distal boundary of the IC side of the OFT. N=12 embryos. Scale bars = 40 μm (A-C); 50 μm (D).

**Figure 1—figure supplement 2.**
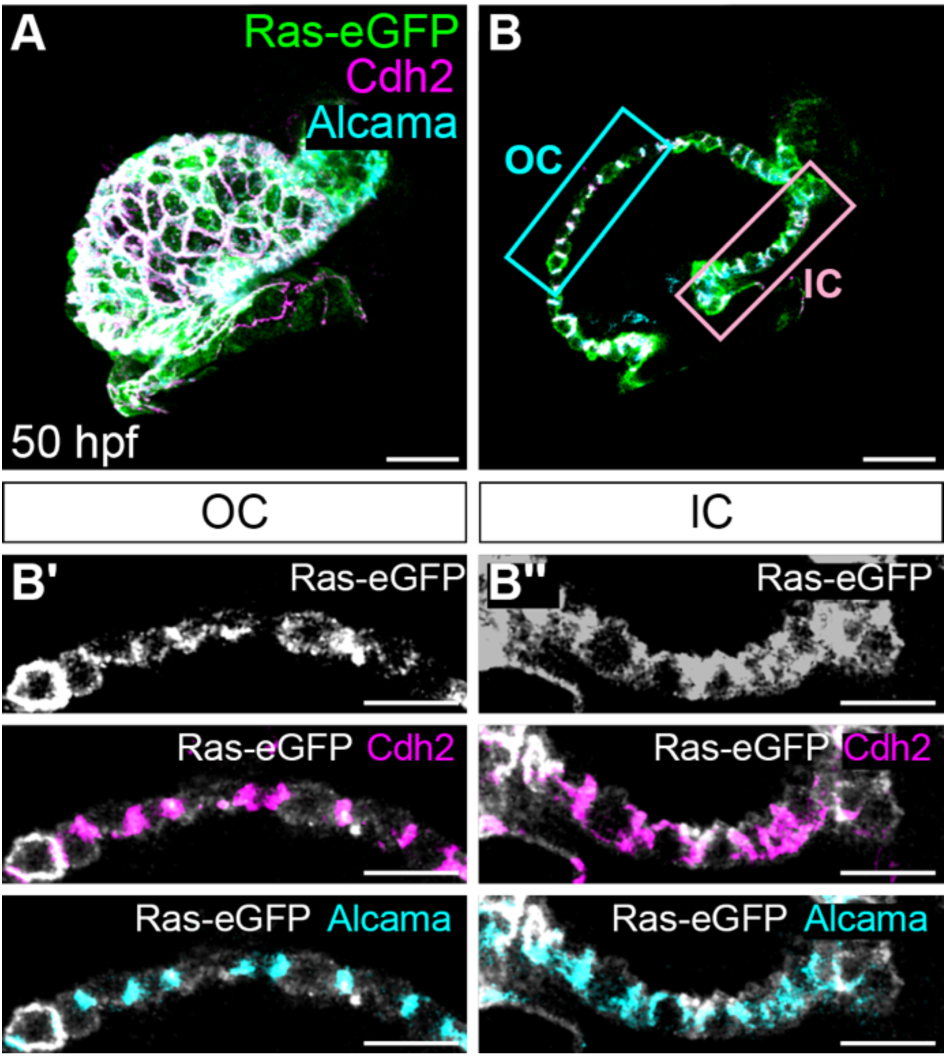
Comparing lateral membrane markers. (**A and B**) Heart from a 50 hpf embryo carrying *Tg(myl7:Ras-eGFP)* (D’Amico et al., 2007), immunostained for membrane-bound eGFP and for cell adhesion molecules Cdh2 and Alcama. (B) shows a section through the 3D reconstruction shown in (A). (Bʹ) and (Bʹʹ) are magnified views of blue (OC) and pink (IC) boxed regions in (B). Colocalization of Cdh2 and Alcama signals with the Ras-eGFP signal at the lateral membranes highlights the ability of these markers to label the entire lateral membrane. N=8 embryos. Scale bars = 30 μm (A and B), 15 μm (Bʹ and Bʹʹ).

**Figure 2—figure supplement 1.**
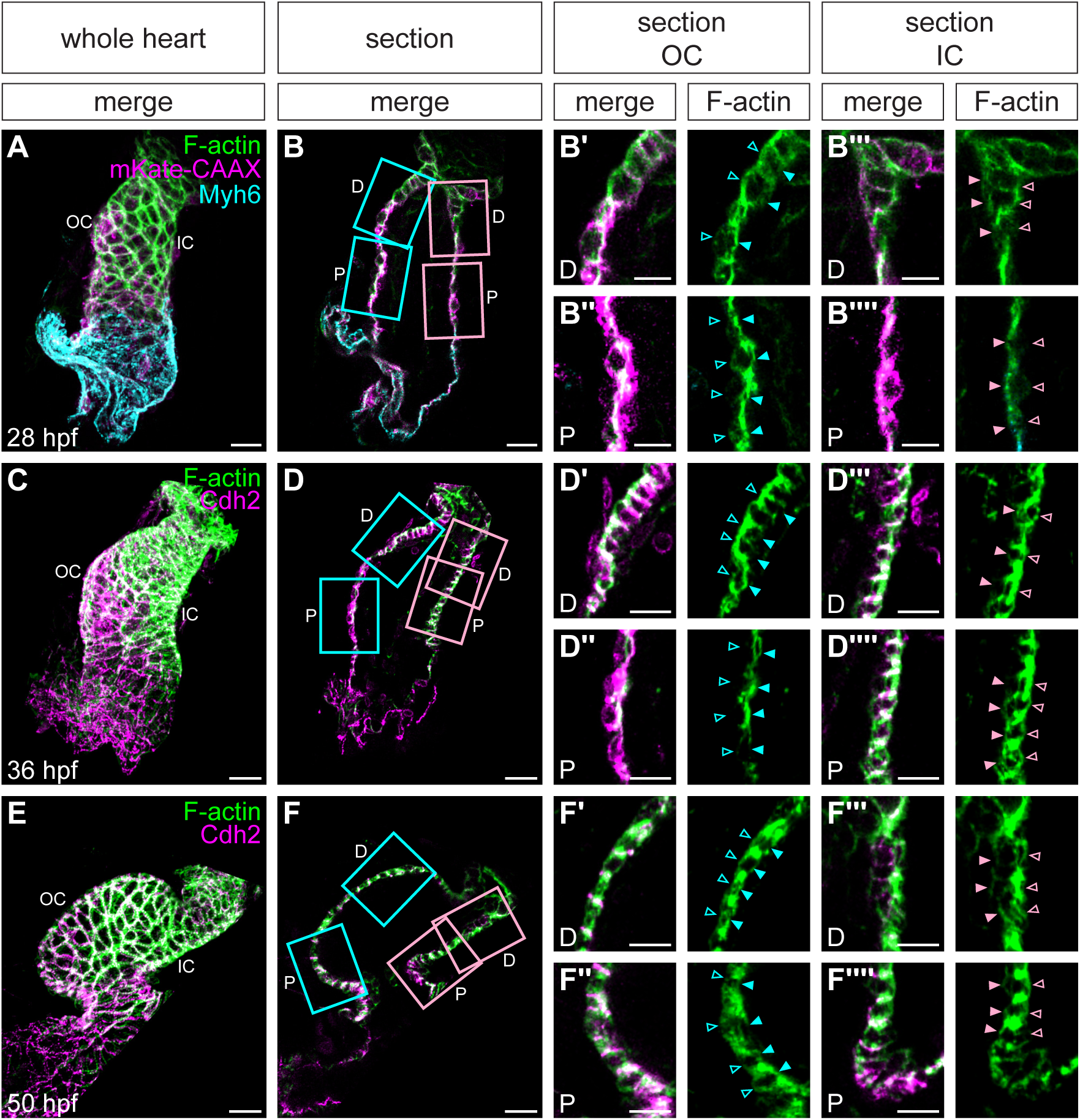
Regionalized subcellular F-actin localization changes as curvatures form. (**A**) Whole heart from a 28 hpf embryo carrying *Tg(myl7:mKate-CAAX)* (Lin et al., 2012), immunostained for membrane-bound mKate and for Myh6 to determine the atrial boundary. (**C and E**) Whole hearts from 36 hpf (C) and 50 hpf (E) embryos immunostained for Cdh2 to label lateral membranes of cardiomyocytes. All hearts stained with Phalloidin to label F-actin. (**B**, **D**, **and F**) Sections through hearts in (A, C, and E). (Bʹ-Bʹʹʹʹ, Dʹ-Dʹʹʹʹ, and Fʹ-Fʹʹʹʹ) Magnified views of blue (OC) and pink (IC) boxed sections in (B, D, and F). Distal (D) regions are shown in (Bʹ, Bʹʹʹ, Dʹ, Dʹʹʹ, Fʹ, and Fʹʹʹ); proximal (P) regions are shown in (Bʹʹ, Bʹʹʹʹ, Dʹʹ, Dʹʹʹʹ, Fʹʹ, and Fʹʹʹʹ). Empty arrowheads: apical membranes. Filled arrowheads: basal membranes. 28 hpf (N=11 embryos); 36 hpf (N=15 embryos); 50 hpf (N=10 embryos). Scale bars = 20 μm (A, B, C, D, E, and F), 10 μm (Bʹ-Bʹʹʹʹ, Dʹ-Dʹʹʹʹ, and Fʹ-Fʹʹʹʹ).

**Figure 2—figure supplement 2.**
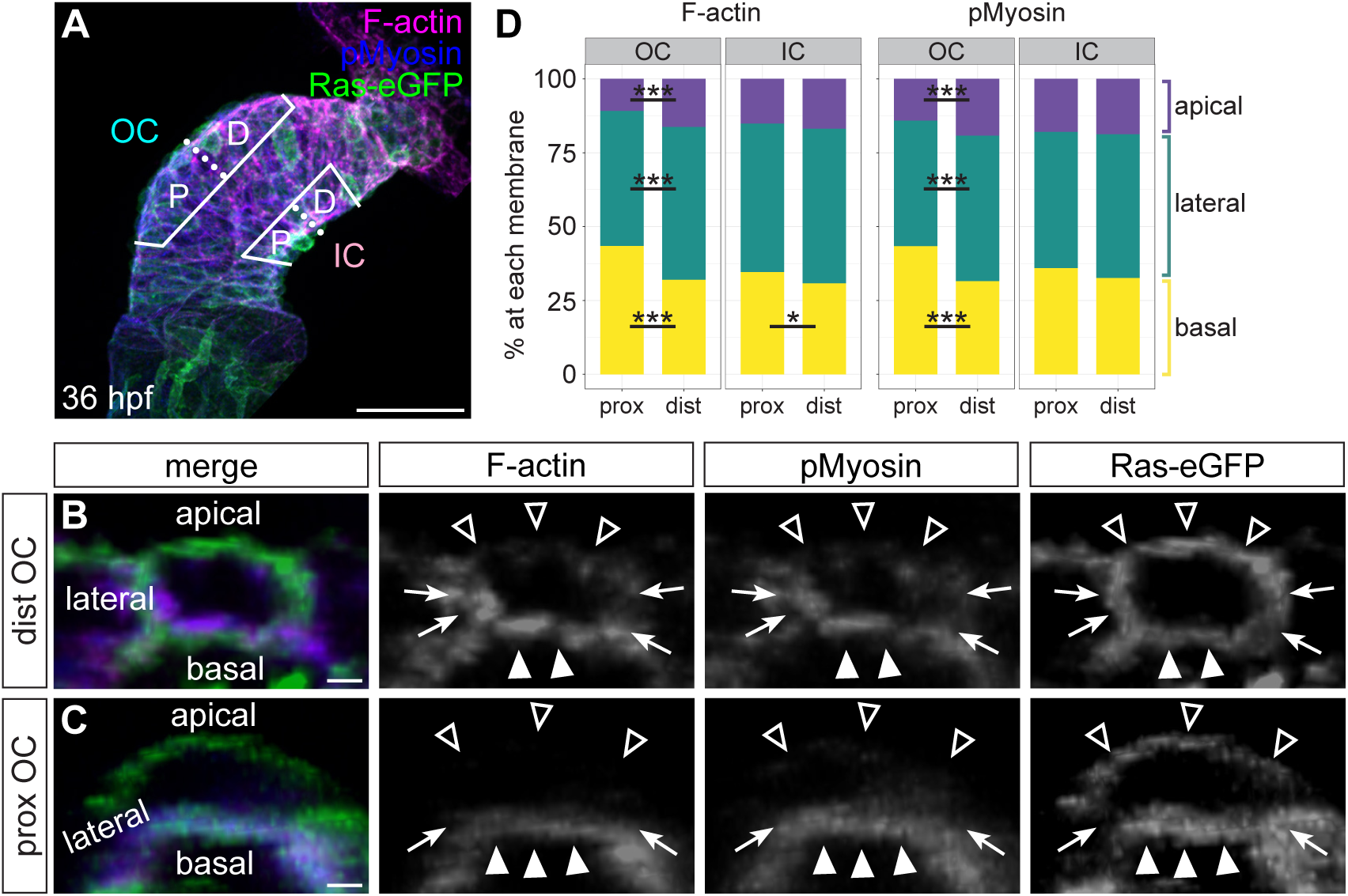
Differences in actomyosin localization along the proximodistal axis of the OC. (**A**) Whole heart from a 36 hpf embryo carrying *Tg(myl7:eGFP-Hsa.HRAS)*, immunostained for membrane-bound GFP and pMyosin and stained with Phalloidin to label F-actin (as in Figure 2A). Solid lines outline the OC and IC; dotted lines show division between the proximal (P) and distal (D) halves of each curvature. (**B and C**) Cross-sections through representative individual cardiomyocytes from the distal OC (B) or proximal OC (C). Empty arrowheads: apical membranes. Filled arrowheads: basal membranes. Arrows: lateral membranes. (**D**) Stacked bar charts showing the mean percentage of F-actin or pMyosin at each membrane. Refer to Tables S3 and S4 for summary statistics. For ratiometric depiction of these data, see Figure 2—figure supplement 3. * denotes p < 0.05 and *** denotes p < 0.001, Wilcoxon test. N=7 embryos; proximal OC (n=97 cells); distal OC (n=102 cells); IC (n=123 cells). Scale bars = 50 μm (A), 2 μm (B and C).

**Figure 2—figure supplement 3.**
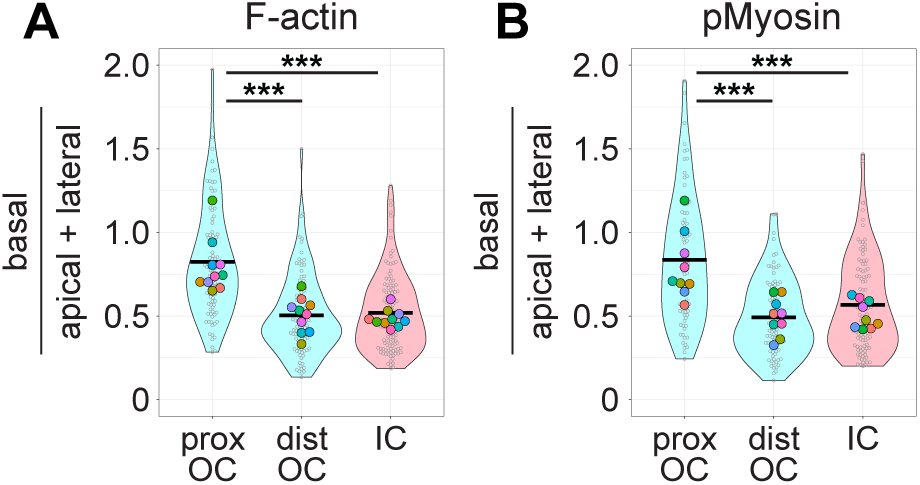
Ratiometric comparison of basal to apical and lateral actomyosin highlights basal enrichment of actomyosin along the proximodistal axis of the OC. (**A**) Violin plot depicts data from Figure 2—figure supplement 2D, recalculated as (mean basal F-actin / (mean apical + mean lateral F-actin)) for individual cells. Each small grey dot represents an individual cell, each black bar represents the mean of values from individual cells, and each large colored dot represents the mean of all values from an individual embryo. (**B**) Violin plot of recalculated values as in (A), but for pMyosin. *** denotes p < 0.001, Wilcoxon test. N=10 embryos; proximal OC (n=97 cells); distal OC (n=102 cells); IC (n=123 cells).

**Figure 3—figure supplement 1.**
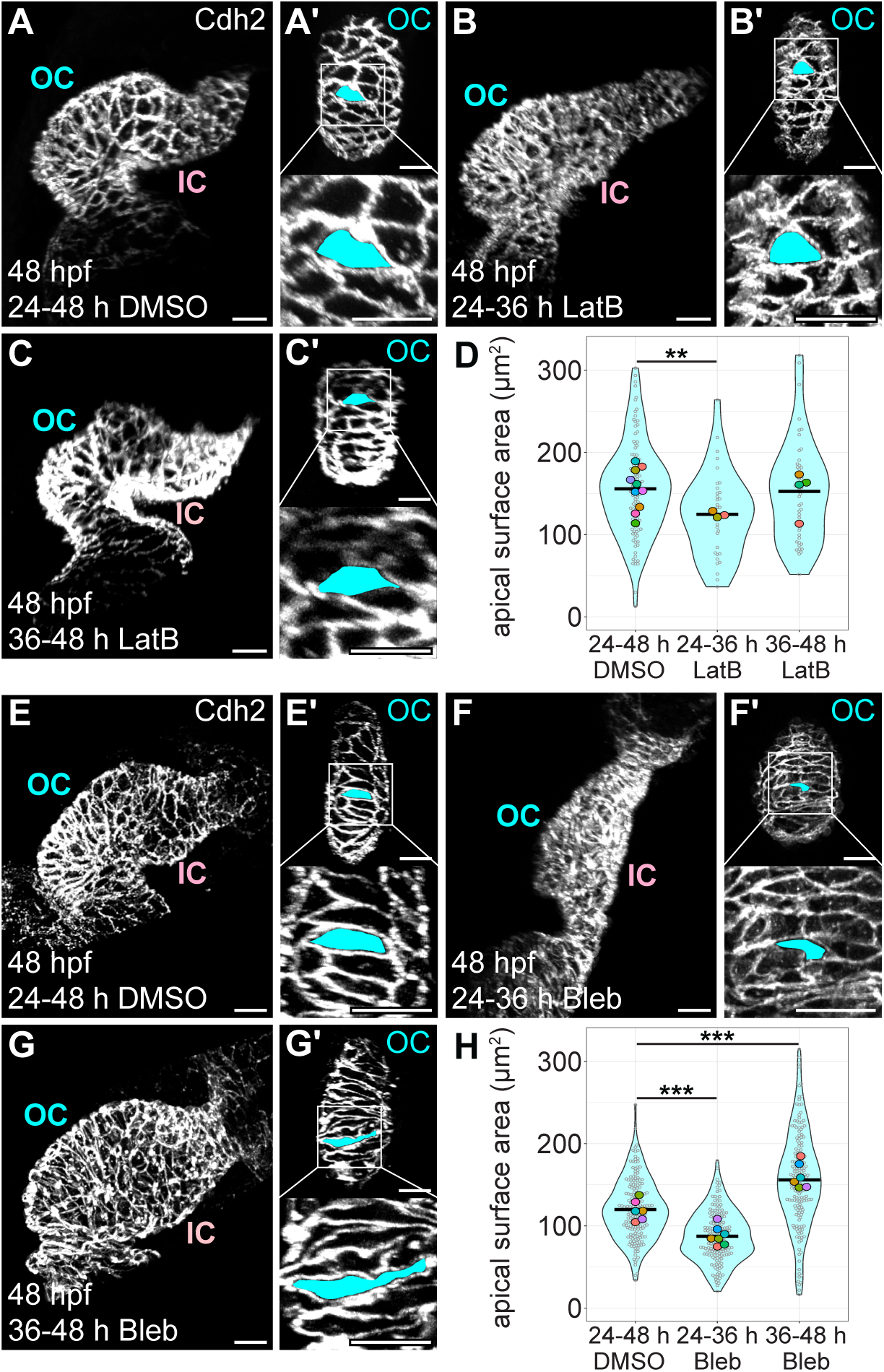
Pharmacological inhibition of actin polymerization or NMII activity during early curvature formation disrupts the acquisition of characteristic OC cardiomyocyte morphology. (**A-C and E-G**) 3D reconstructions of 48 hpf wild-type hearts (A-C and E-G) and OCs (Aʹ-Cʹ and Eʹ-Gʹ) from embryos treated with DMSO from 24-48 hpf (A and E), 100 ng/mL Latrunculin B (Lat B) from 24-36 hpf (B), 100 ng/mL Lat B from 36-48 hpf (C), 5 uM Blebbistatin (Bleb) from 24-36 hpf (F), or 5 uM Bleb from 36-48 hpf (G). Immunostaining for Cdh2 labels lateral membranes of cardiomyocytes. OCs (Aʹ, Bʹ, Cʹ, Eʹ, Fʹ, and Gʹ) are shown for hearts in (A, B, C, E, F, and G). Insets show higher magnification. Apical surface area of an individual cardiomyocyte is illustrated by blue fill. (**D and H**) Violin plots compare apical surface area of OC cells following different treatments. Each small grey dot represents an individual cell, each black bar represents the mean of values from individual cells, and each large colored dot represents the mean of all values from an individual embryo. ** denotes p < 0.01 and *** denotes p < 0.001, Wilcoxon test. For Lat B: DMSO 24-48 hpf (N=10 embryos, n=120 cells); Lat B 24-36 hpf (N=3 embryos, n=35 cells); Lat B 36-48 hpf (N=4 embryos, n=48 cells). For Bleb: DMSO 24-48 hpf (N=6 embryos, n=183 cells); Lat B 24-36 hpf (N=7 embryos, n=169 cells); Lat B 36-48 hpf (N=6 embryos, n=170 cells). Scale bars = 20 μm.

**Figure 3—figure supplement 2.**
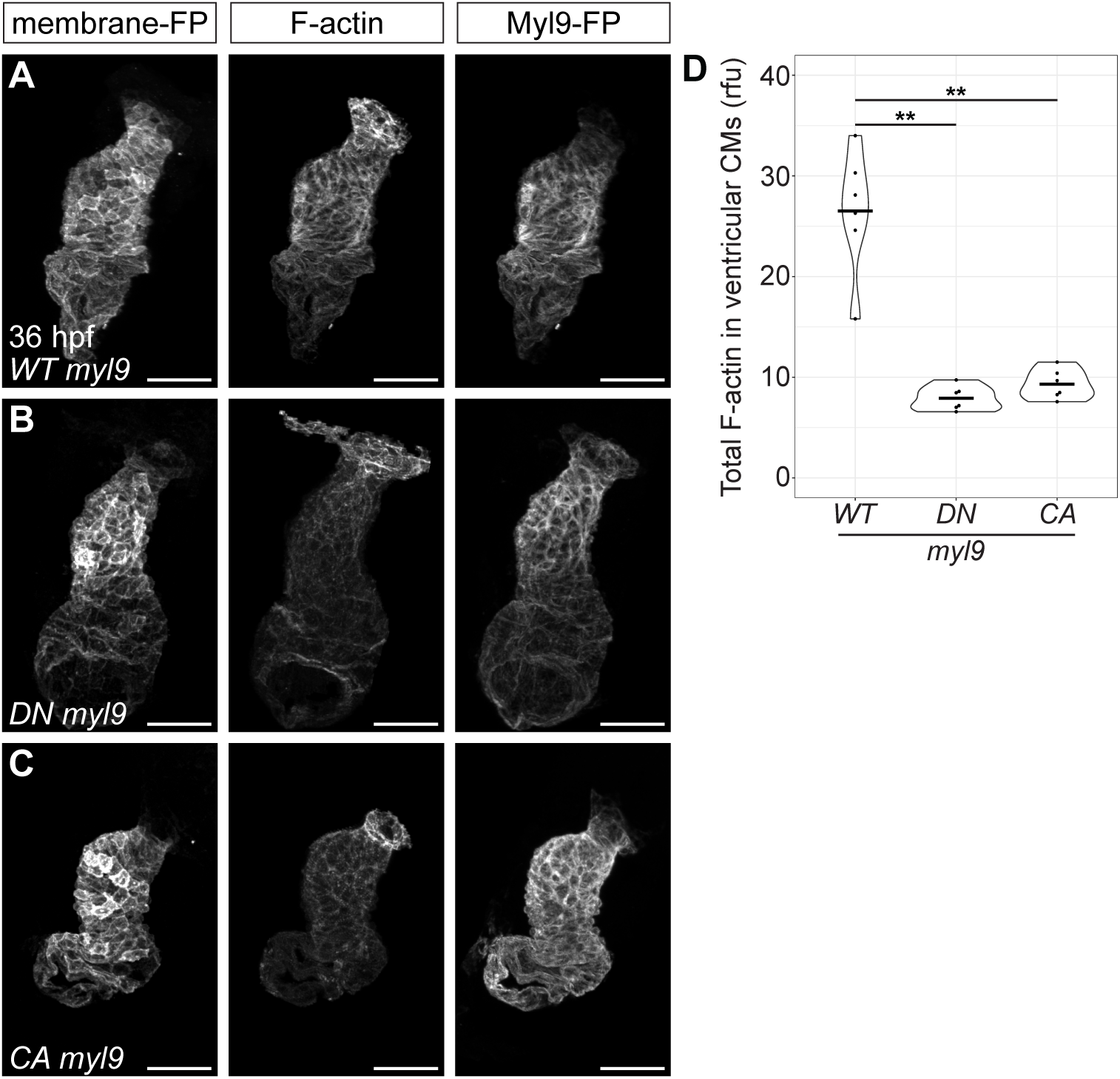
Tissue-specific modulation of NMII activity reduces the total amount of F-actin in the ventricular myocardium. (**A**) 3D reconstruction of a 36 hpf heart expressing *Tg(myl7:WT-myl9-mScarlet)* and *Tg(myl7:eGFP-Hsa.HRAS)*, immunostained for mScarlet and membrane-bound GFP. (**B**) 3D reconstruction of a 36 hpf heart expressing *Tg(myl7:DN-myl9-eGFP)* and *Tg(myl7:mKate-CAAX)*, immunostained for GFP and membrane-bound mKate. (**C**) 3D reconstruction of a 36 hpf heart expressing *Tg(myl7:CA-myl9-eGFP)* and *Tg(myl7:mKate-CAAX)*, immunostained for GFP and membrane-bound mKate. All hearts are stained with Phalloidin to label F-actin. (**D**) Violin plot compares relative fluorescence units (rfu) of total F-actin in the ventricular myocardium between embryos carrying the different transgenes. Each dot represents one ventricle. ** denotes p < 0.01, Wilcoxon test. N=6 embryos for each transgenic line. Scale bars = 50 μm.

**Figure 3—figure supplement 3.**
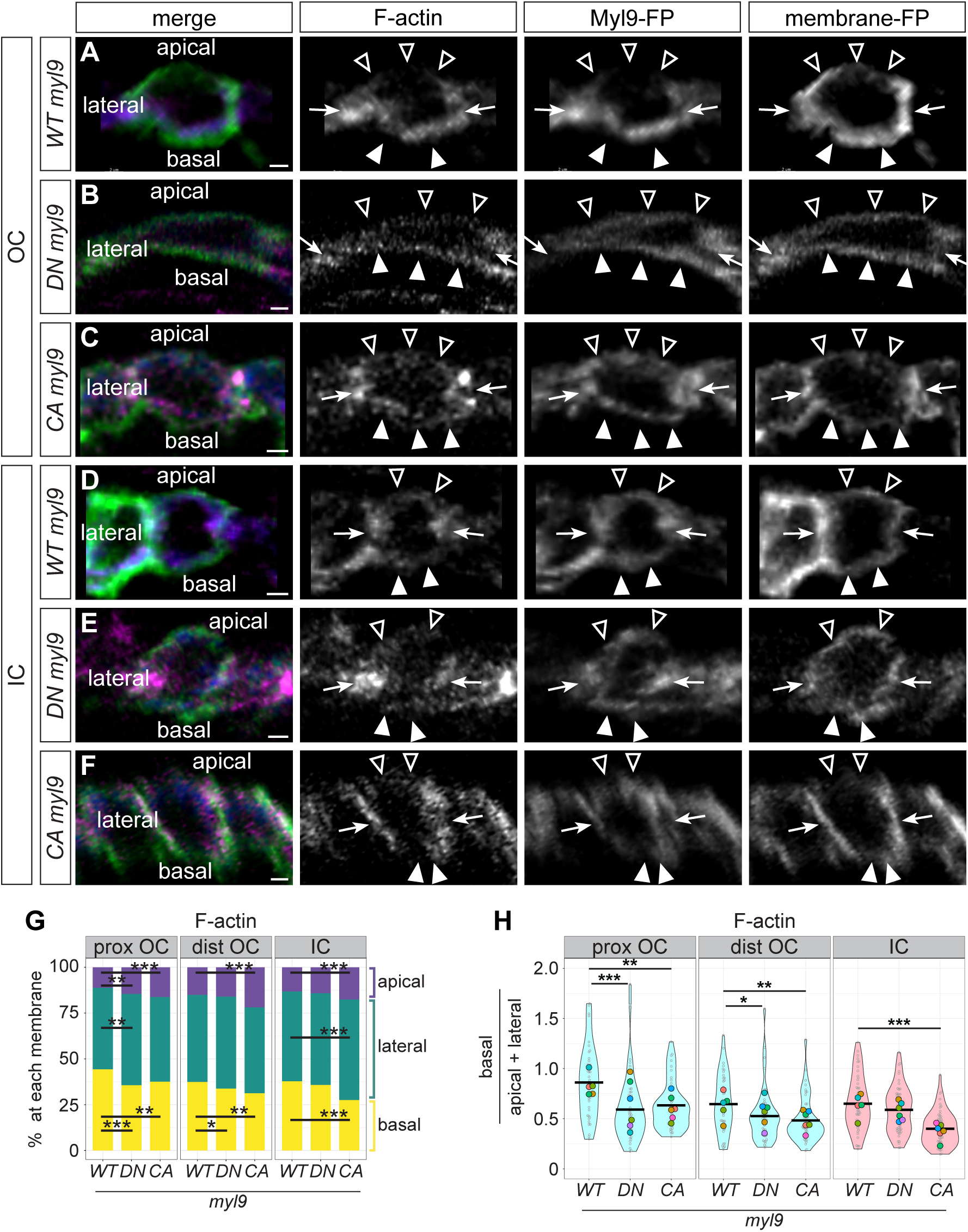
Tissue-specific modulation of NMII activity alters subcellular F-actin localization in OC and IC cardiomyocytes. (**A-F**) Cross-sections through representative individual cardiomyocytes from the OC (A-C) or IC (D-F) of 36 hpf embryos expressing *Tg(myl7:WT-myl9-mScarlet)* and *Tg(myl7:eGFP-Hsa.HRAS)* (A, D), *Tg(myl7:DN-myl9-eGFP)* and *Tg(myl7:mKate-CAAX)* (B, E), or *Tg(myl7:CA-myl9-eGFP)* and *Tg(myl7:mKate-CAAX)* (C and F), immunostained as in Figure 3—figure supplement 2. All hearts are stained with Phalloidin to label F-actin. In the “merge” column, magenta represents F-actin, blue represents the Myl9-bound fluorescent protein, and green represents the membrane-bound fluorescent protein. Empty arrowheads: apical membranes. Filled arrowheads: basal membranes. Arrows: lateral membranes. (**G**) Stacked bar charts show the mean percentage of F-actin at each membrane. Refer to Table S5 for summary statistics. (**H**) Data from (G), recalculated as in Figure 2—figure supplement 3. Each small grey dot represents an individual cell, each black bar represents the mean of values from individual cells, and each large colored dot represents the mean of all values from an individual embryo. * denotes p < 0.05, ** denotes p < 0.01, and *** denotes p < 0.001, Wilcoxon test. For *Tg(myl7:WT-myl9-mScarlet)*: proximal OC (N=5 embryos, n=39 cells); distal OC (N=5 embryos, n=42 cells); IC (N=5 embryos, n=50 cells). For *Tg(myl7:DN-myl9-eGFP)*: proximal OC (N=6 embryos, n=45 cells); distal OC (N=6 embryos, n=46 cells); IC (N=6 embryos, n=53 cells). For *Tg(myl7CA-myl9-eGFP)*: proximal OC (N=6 embryos, n=43 cells); distal OC (N=6 embryos, n=44 cells); IC (N=6 embryos, n=55 cells). Scale bars = 2 μm.

**Figure 5—figure supplement 1.**
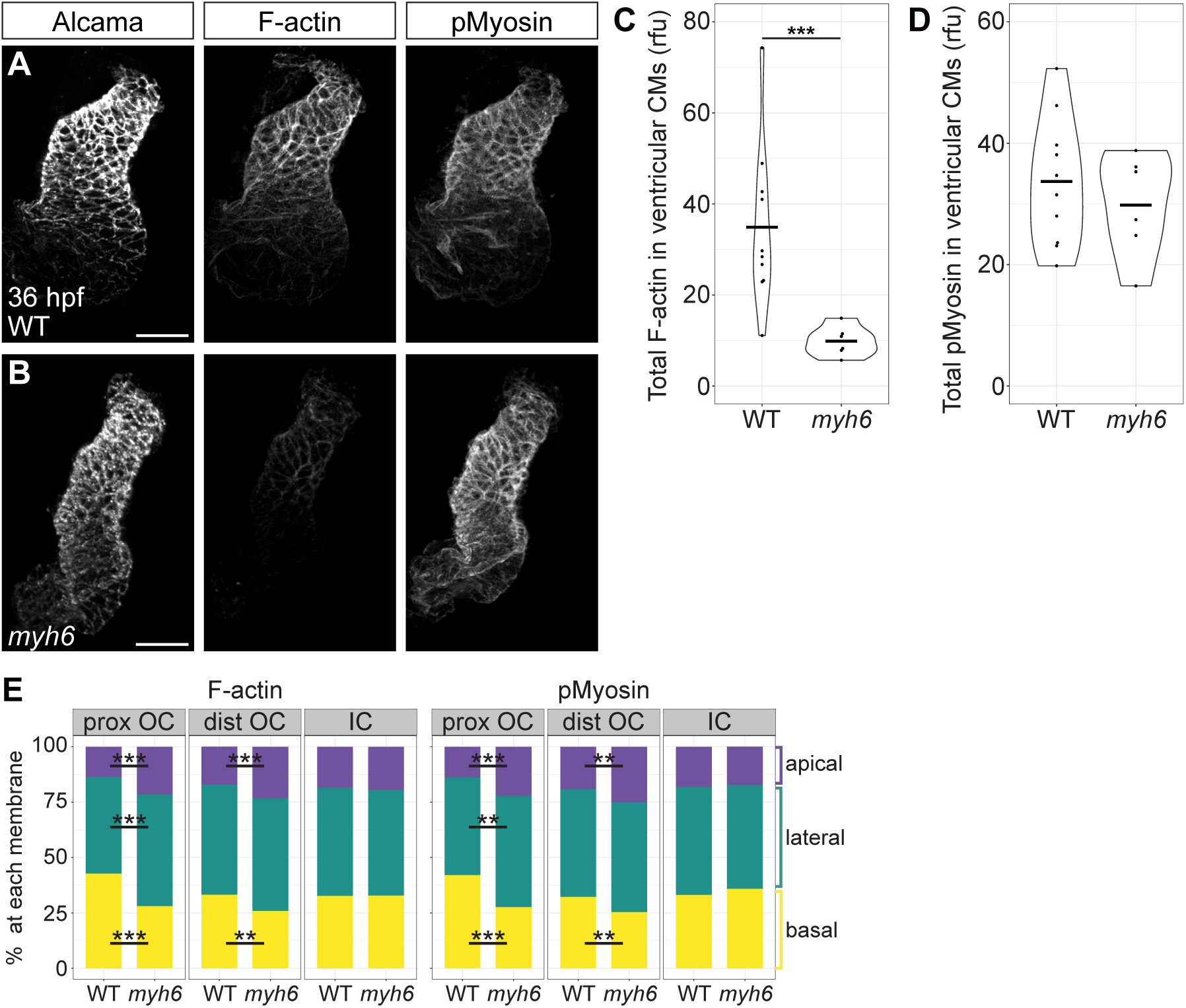
Reduced blood flow dampens the total amount of F-actin in the ventricular myocardium. (**A and B**) 3D reconstructions of wild-type (A) and *myh6* mutant (B) hearts at 36 hpf, immunostained for pMyosin and stained with Phalloidin to label F-actin. Immunostaining for Alcama labels lateral membranes of cardiomyocytes. (**C and D**) Violin plots compare rfu of total F-actin (C) or pMyosin (D) in the ventricular myocardium between wild-type and *myh6* mutant hearts. Each dot represents one ventricle. (**E**) Stacked bar charts include the same set of cardiomyocytes as in Figure 5O and P and show the mean percentage of F-actin or pMyosin at each membrane. Refer to Tables S6 and S7 for summary statistics. ** denotes p < 0.01 and *** denotes p < 0.001, Wilcoxon test. For whole ventricle signal intensities: wild-type (N=10 embryos); *myh6* (N=6 embryos). For actomyosin localization: wild-type proximal OC (N=5 embryos, n=39 cells); *myh6* proximal OC (N=4 embryos, n=32 cells); wild-type distal OC (N=5 embryos, n=41 cells); *myh6* distal OC (N=4 embryos, n=24 cells); wild-type IC (N=5 embryos, n=54 cells); *myh6* IC (N=4 embryos, n=35 cells). Scale bars = 40 μm.

**Figure 6—figure supplement 1.**
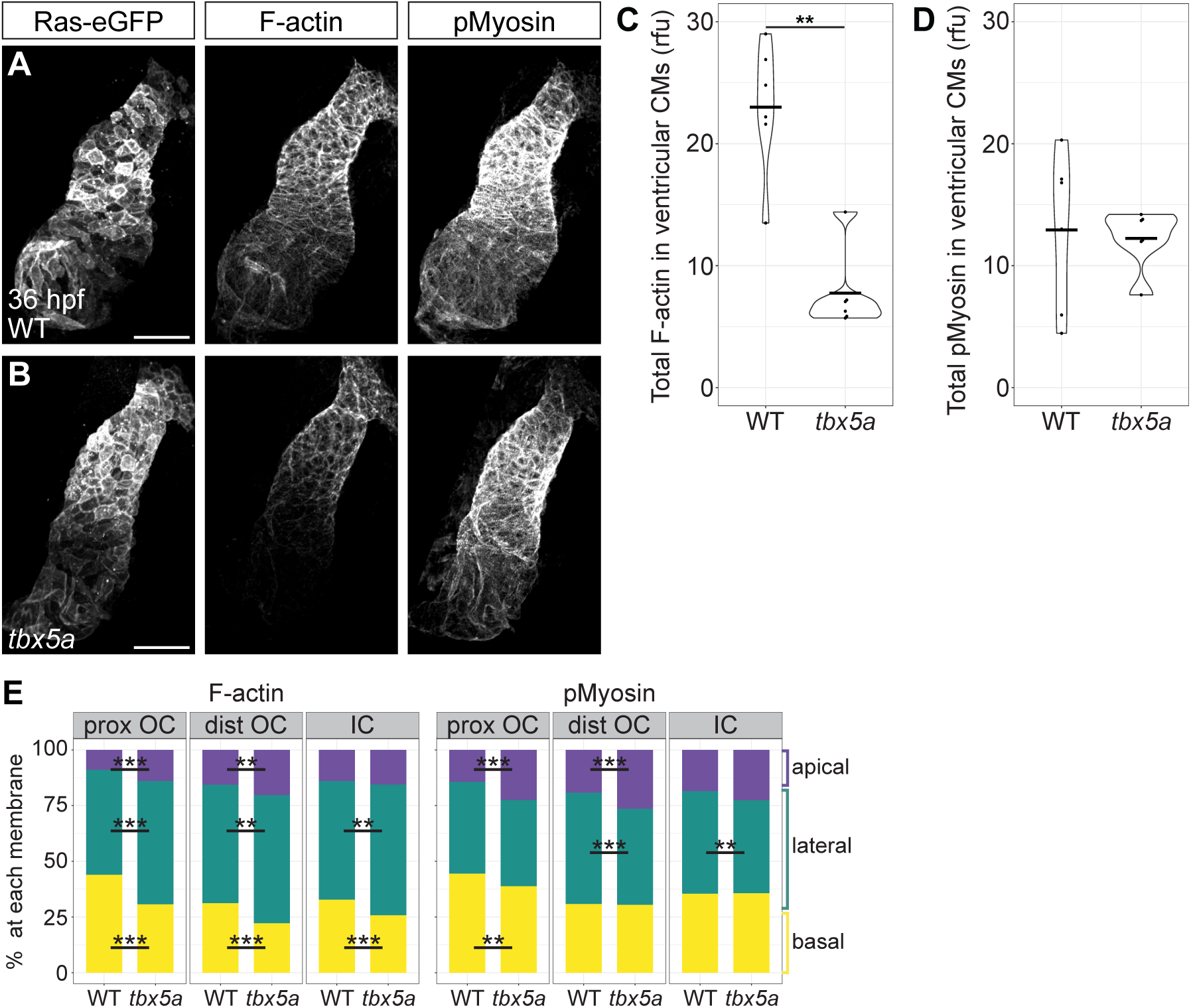
*tbx5a* mutants exhibit a reduced amount of total F-actin in the ventricular myocardium. (**A and B**) 3D reconstructions of wild-type (A) and *tbx5a* mutant (B) hearts at 36 hpf, immunostained for pMyosin and membrane-bound eGFP and stained with Phalloidin to label F-actin. (**C and D**) Violin plots compare rfu of total F-actin (C) or pMyosin (D) in the ventricular myocardium between wild-type and *tbx5a* mutant hearts. Each dot represents one ventricle. (**E**) Stacked bar charts include the same set of cardiomyocytes as in Figure 6O and P and show the mean percentage of F-actin or pMyosin at each membrane. Refer to Tables S8 and S9 for summary statistics. ** denotes p < 0.01 and *** denotes p < 0.001, Wilcoxon test. For whole ventricle signal intensities: wild-type (N=6 embryos); *tbx5a* (N=6 embryos). For actomyosin localization: wild-type proximal OC (N=5 embryos, n=58 cells); *tbx5a* proximal OC (N=5 embryos, n=64 cells); wild-type distal OC (N=5 embryos, n=61 cells); *tbx5a* distal OC (N=5 embryos, n=67 cells); wild-type IC (N=5 embryos, n=69 cells); *tbx5a* IC (N=5 embryos, n=81 cells). Scale bars = 50 μm.

**Figure 7—figure supplement 1.**
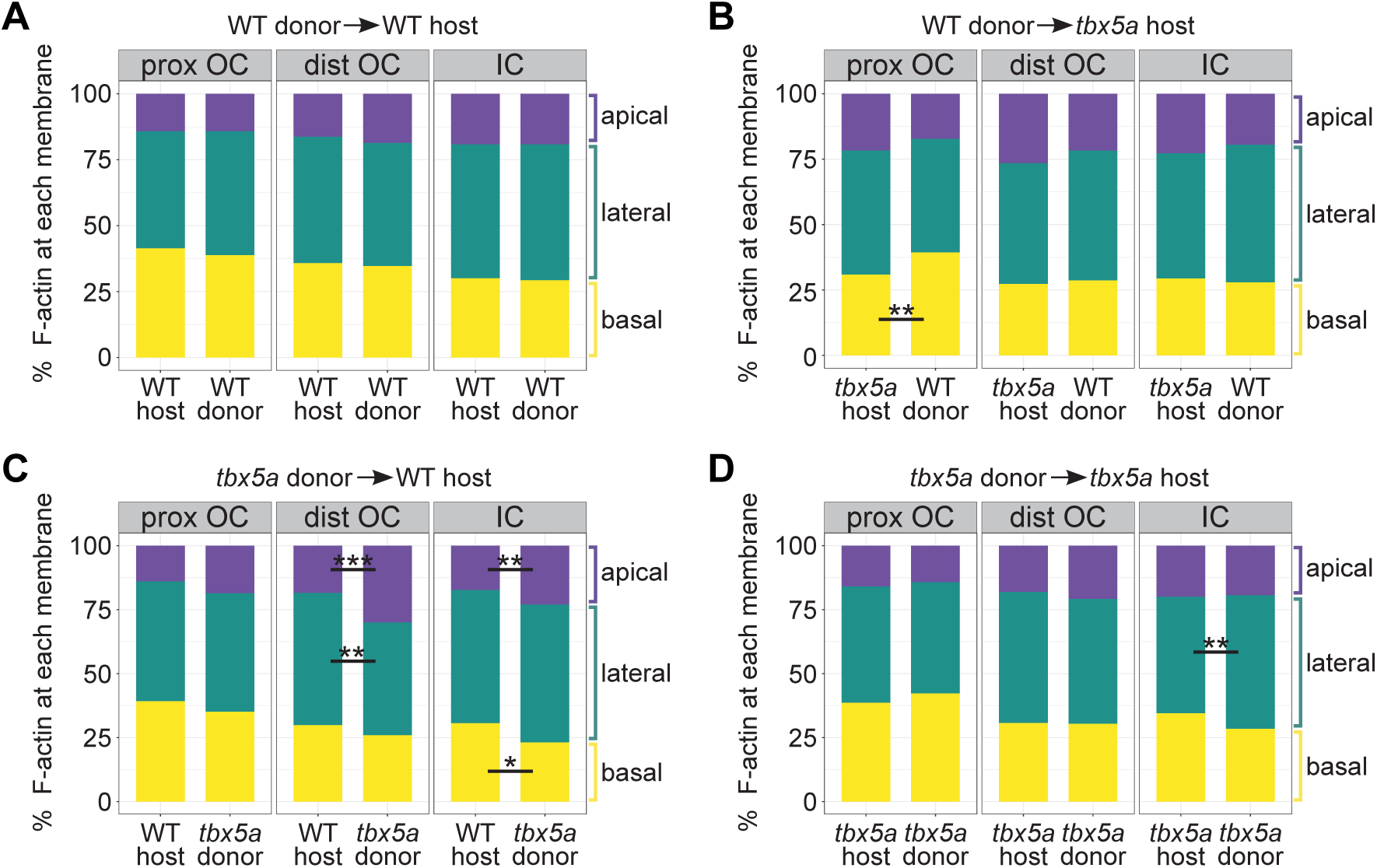
Subcellular F-actin organization in mosaic hearts from Figure 7, shown as percentage of F-actin at each membrane. (**A-D**) Stacked bar charts include the same sets of cardiomyocytes as in Figure 7G, J, M, and P and show the mean percentage of F-actin at each membrane. Refer to Tables S10, S11, S12, and S13 for summary statistics. Note that, in the *tbx5a* mutant into wild-type scenario (C), some F-actin proportions are significantly different between host- and donor-derived cardiomyocytes in the distal OC and the IC. However, the nature of these differences do not translate into differences between the ratios of (basal F-actin / (apical + lateral F-actin)) presented in Figure 7M. * denotes p < 0.05, ** denotes p < 0.01, and *** denotes p < 0.001, Wilcoxon test. For WT into WT transplants: host proximal OC (N=6 embryos, n=55 cells); donor proximal OC (N=7 embryos, n=14 cells); host distal OC (N=5 embryos, n=30 cells); donor distal OC (N=6 embryos, n=29 cells); host IC (N=4 embryos, n=30 cells); donor IC (N=4 embryos, n=29 cells). For WT into *tbx5a* transplants: host proximal OC (N=5 embryos, n=40 cells); donor proximal OC (N=5 embryos, n=25 cells); host distal OC (N=5 embryos, n=40 cells); donor distal OC (N=5 embryos, n=35 cells); host IC (N=5 embryos, n=18 cells); donor IC (N=5 embryos, n=14 cells). For *tbx5a* into WT transplants: host proximal OC (N=7 embryos, n=38 cells); donor proximal OC (N=5 embryos, n=14 cells); host distal OC (N=6 embryos, n=28 cells); donor distal OC (N=5 embryos, n=13 cells); host IC (N=5 embryos, n=44 cells); donor IC (N=5 embryos, n=18 cells). For *tbx5a* into *tbx5a* transplants: host proximal OC (N=4 embryos, n=16 cells); donor proximal OC (N=2 embryos, n=3 cells); host distal OC (N=4 embryos, n=21 cells); donor distal OC (N=4 embryos, n=14 cells); host IC (N=5 embryos, n=36 cells); donor IC (N=5 embryos, n=14 cells).

**Figure 7—figure supplement 2.**
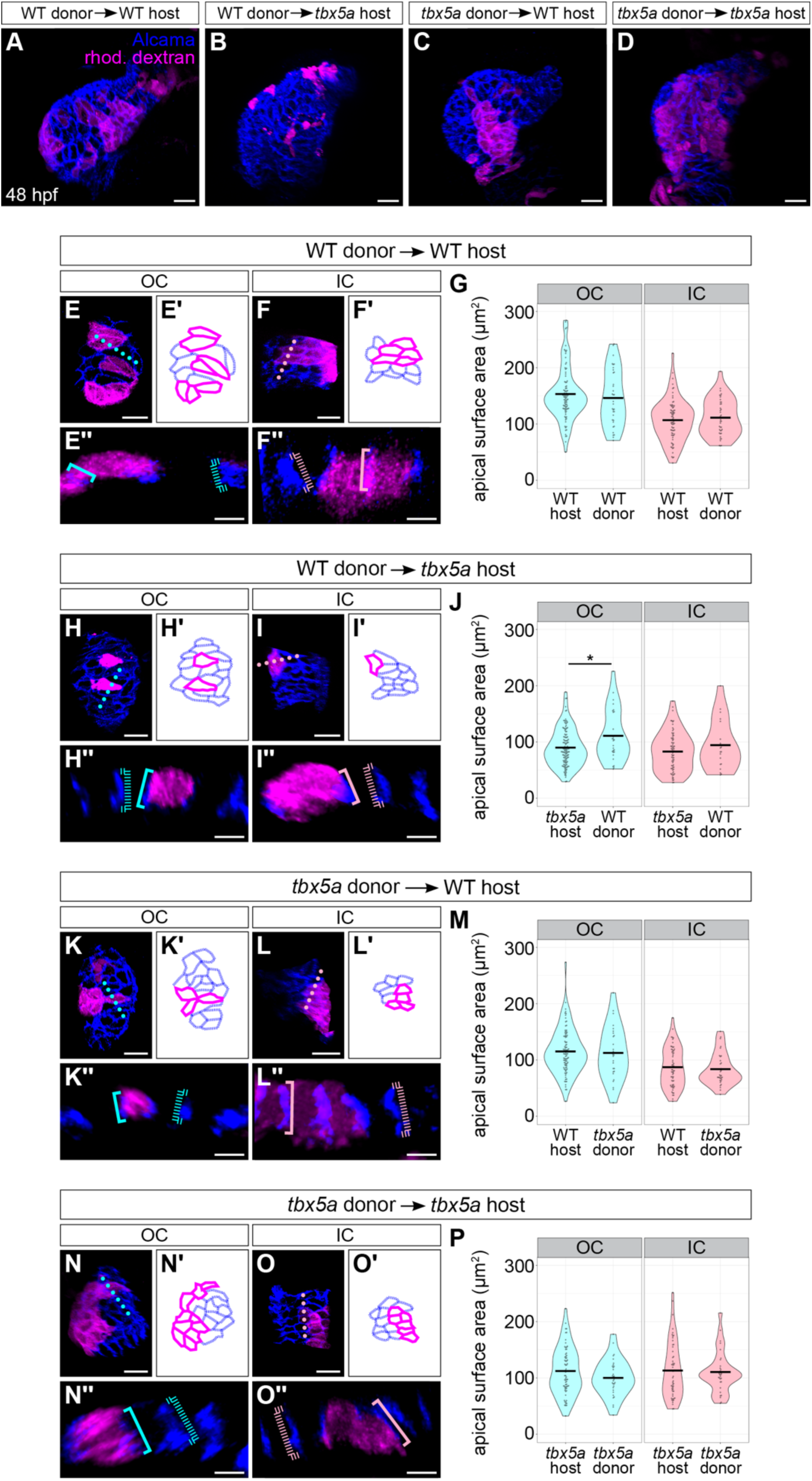
*tbx5a* functions in a partially cell-autonomous manner to support planar expansion of OC cardiomyocytes. (**A-D**) 3D reconstructions show examples of mosaic 48 hpf hearts resulting from blastomere transplantation. Immunostaining for Alcama labels lateral membranes of cardiomyocytes (blue), and donor-derived cells are labeled with rhodamine-dextran (magenta). OCs (**E**, **H**, **K**, **and N**) and ICs (**F**, **I**, **L**, **and O**) show additional examples of each transplant scenario. (Eʹ, Fʹ, Hʹ, Iʹ, Kʹ, Lʹ, Nʹ, and Oʹ) Tracings of cells in (E, F, H, I, K, L, N, and O). Magenta outlines indicate donor-derived cardiomyocytes; blue outlines indicate host-derived cells. (Eʹʹ, Fʹʹ, Hʹʹ, Iʹʹ, Kʹʹ, Lʹʹ, Nʹʹ, and Oʹʹ) Cross-sections through positions indicated by dotted lines in (E, F, H, I, K, L, N, and O). Blue and pink brackets highlight apicobasal length of individual cells, with dashed brackets for host-derived cells and solid brackets for donor-derived cardiomyocytes. (**G**, **J, M, and P**) Violin plots compare apical surface area of host-derived cells to those of donor-derived cardiomyocytes. Each dot represents an individual cell. * denotes p < 0.05, ** denotes p < 0.01, and *** denotes p < 0.001, Wilcoxon test. For WT into WT transplants: host OC (N=7 embryos, n=92 cells); donor OC (N=7 embryos, n=34 cells); host IC (N=6 embryos, n=73 cells); donor IC (N=6 embryos, n=39 cells). For WT into *tbx5a* transplants: host OC (N=7 embryos, n=123 cells); donor OC (N=7 embryos, n=27 cells); host IC (N=7 embryos, n=72 cells); donor IC (N=7 embryos, n=18 cells). For *tbx5a* into WT transplants: host OC (N=8 embryos, n=93 cells); donor OC (N=8 embryos, n=28 cells); host IC (N=5 embryos, n=59 cells); donor IC (N=5 embryos, n=34 cells). For *tbx5a* into *tbx5a* transplants: host OC (N=3 embryos, n=67 cells); donor OC (N=3 embryos, n=32 cells); host IC (N=3 embryos, n=56 cells); donor IC (N=3 embryos, n=30 cells). Scale bars = 20 μm (A-D, E, F, H, I, K, L, N, and O); 5 μm (Eʹʹ, Fʹʹ, Hʹʹ, Iʹʹ, Kʹʹ, Lʹʹ, Nʹʹ, and Oʹʹ).

**Figure 7—figure supplement 3.**
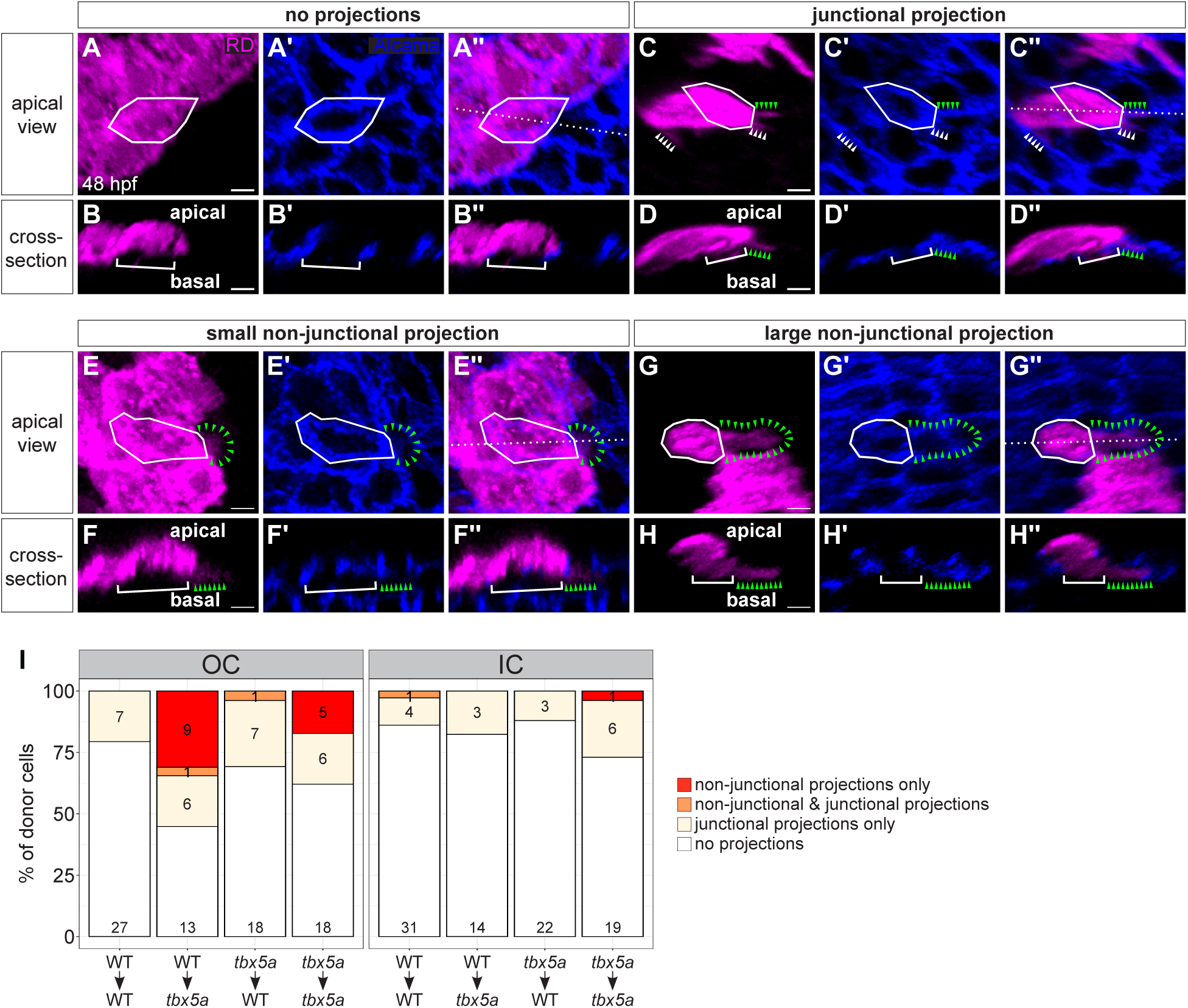
Wild-type cardiomyocytes extend excessive basal projections when positioned next to *tbx5a* mutant cardiomyocytes. (**A**, **C**, **E**, **and G**) Apical views of mosaic hearts at 48 hpf provide examples of donor cardiomyocytes that can be categorized by their types of basal projections. Immunostaining for Alcama labels lateral membranes of cardiomyocytes (blue), and donor-derived cardiomyocytes are labeled with rhodamine dextran (RD, magenta). In each panel, the apical surface of a single donor-derived cardiomyocyte is outlined. (A) Outlined cardiomyocyte has no visible projections. (C) Outlined cardiomyocyte has only thin projections (arrowheads) that extend along the junction between two neighboring cells. (E) Outlined cardiomyocyte has one small non-junctional projection (a broader projection that extends underneath a neighboring cell, instead of between two neighbors), outlined with arrowheads. (G) Outlined cardiomyocyte has one large non-junctional projection, outlined with arrowheads. (**B**, **D**, **F**, **and H**) Cross-sections through outlined cardiomyocytes in (A, C, E, and G); positions of cross-sections are noted by dotted lines in (Aʹʹ, Cʹʹ, Eʹʹ, and Gʹʹ). Solid brackets highlight the width of the main mass of the outlined donor-derived cardiomyocyte; green arrowheads highlight the basal projection. Cells with projections were detected in both proximal and distal regions of the curvatures. (**I**) Stacked bar charts show the proportions of donor-derived cardiomyocytes exhibiting different categories of projections, organized by transplant scenario and curvature. Only donor-derived cardiomyocytes that directly contact host-derived cardiomyocytes were considered in this analysis. Numerals on the charts refer to the number of donor cells in each category. Scale bars = 5 μm.

**Figure 7—figure supplement 4.**
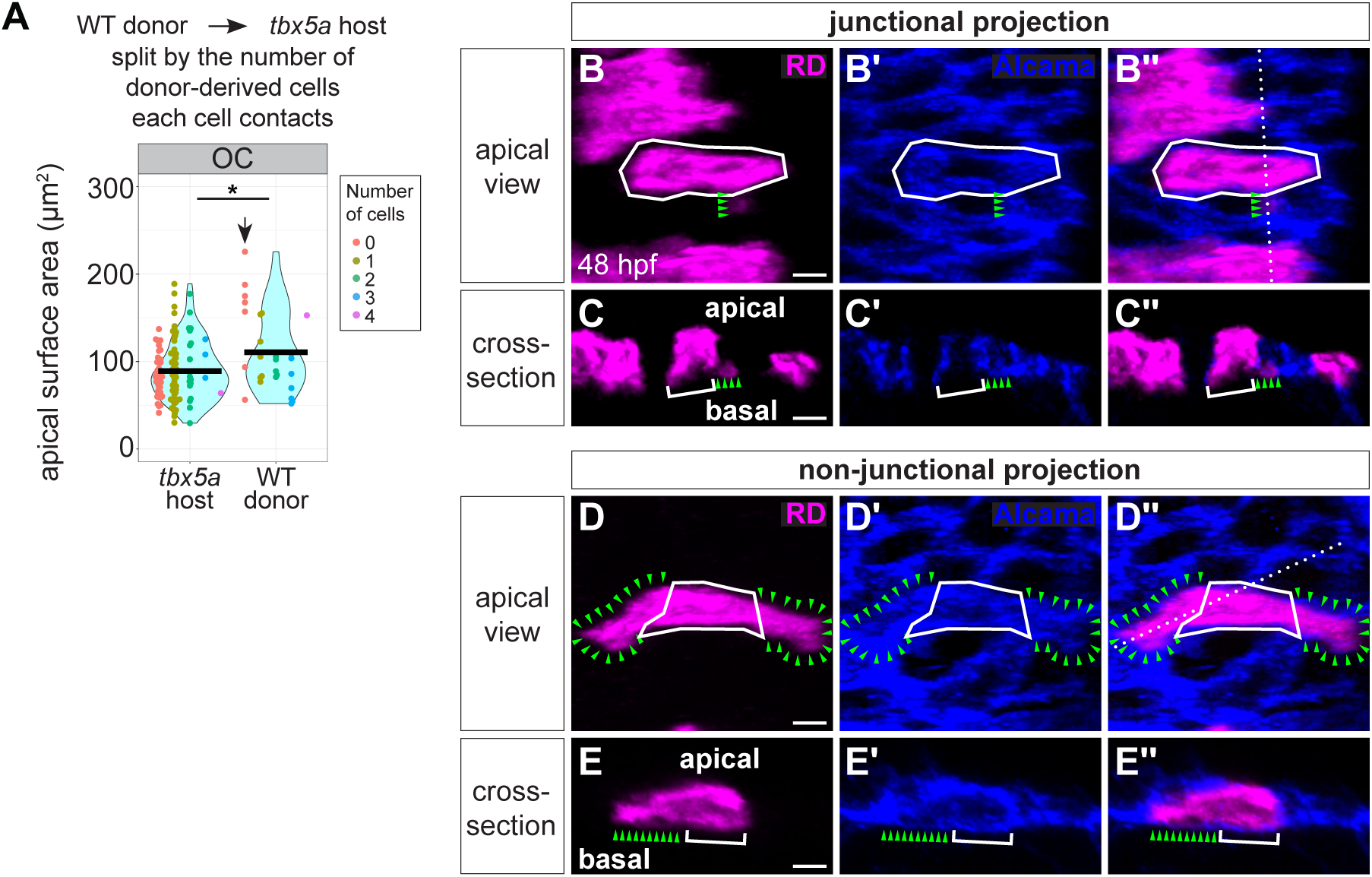
Single-cell wild-type clones in a *tbx5a* mutant OC send basal projections between and under neighboring host-derived cardiomyocytes. (**A**) OC data from Figure 7—figure supplement 2J, with individual data points colored based on how many other donor-derived cardiomyocytes that cardiomyocyte contacts. A donor-derived cardiomyocyte in contact with no other donor-derived cardiomyocytes is considered a single-cell clone (coral dots; arrow). These 7 cardiomyocytes have, on average, a larger apical surface area than donor-derived cardiomyocytes in contact with other donor-derived cardiomyocytes. (**B-E**) Two examples of the 7 donor-derived cardiomyocytes highlighted in (A). Three of the 7 single-cell donor-derived clones highlighted in (A) send thin basal projections that extend along the junction between two neighboring host-derived cardiomyocytes (example shown in B and C). The other 4 of the 7 single-cell donor-derived clones highlighted in (A) send larger basal projections underneath neighboring host-derived cardiomyocytes (example shown in D and E). (B and D) Apical views of examples of donor cell projections. Immunostaining for Alcama labels lateral membranes of cardiomyocytes (blue), and donor-derived cells are labeled with rhodamine dextran (RD, magenta). In each panel, the apical surface of a single donor-derived cardiomyocyte is outlined with a solid line. (B) Outlined cardiomyocyte has a thin projection (denoted with green arrowheads) that extend between two neighboring cells. (D) Outlined cardiomyocyte has two large non-junctional projections (*i.e.* broader projections that extend underneath a neighboring cardiomyocyte, instead of between two neighbors; outlined with green arrowheads). (C and E) Cross-sections through outlined cardiomyocytes in (B and D); positions of cross-sections noted by dotted lines in (Bʹʹ and Dʹʹ). Solid brackets highlight the width of the main mass of the outlined donor-derived cardiomyocyte; green arrowheads highlight the basal projection. * denotes p < 0.05, Wilcoxon test. Scale bars = 5 μm.

---

**Table S1.** Summary statistics corresponding to F-actin localization in Figure 2E. The stacked bar chart in Figure 2E shows the mean percentage of F-actin at each membrane. These summary statistics provide information for assessing the distribution of these data. S.D.: standard deviation; S.E.M.: standard error of the mean.

| Genotype/Stage | Location | # of cells | Membrane | Mean % | S.D. | S.E.M. | Range of % |
| --- | --- | --- | --- | --- | --- | --- | --- |
| wild-type<br>36 hpf | OC | 199 | apical | 13.7 | 6.5 | 0.5 | 2.0 - 41.7 |
|  |  |  | basal | 37.6 | 11.2 | 0.8 | 11.8 - 66.4 |
|  |  |  | lateral | 48.7 | 8.8 | 0.6 | 19.4 - 71.3 |
|  | IC | 123 | apical | 16.0 | 5.9 | 0.5 | 3.0 - 32.0 |
|  |  |  | basal | 32.8 | 9.3 | 0.8 | 15.7 - 56.2 |
|  |  |  | lateral | 51.2 | 8.6 | 0.8 | 25.6 - 69.7 |

**Table S2.** Summary statistics corresponding to pMyosin localization in Figure 2E. Information is presented as in Table S1.

| Genotype/Stage | Location | # of cells | Membrane | Mean % | S.D. | S.E.M. | Range of % |
| --- | --- | --- | --- | --- | --- | --- | --- |
| wild-type<br>36 hpf | OC | 174 | apical | 16.7 | 6.9 | 0.5 | 4.8 - 43.7 |
|  |  |  | basal | 37.3 | 12.0 | 0.9 | 10.2 - 65.6 |
|  |  |  | lateral | 45.9 | 8.6 | 0.6 | 23.3 - 67.8 |
|  | IC | 109 | apical | 18.4 | 6.7 | 0.6 | 5.6 - 39.4 |
|  |  |  | basal | 34.4 | 10.7 | 1.0 | 16.7 - 59.5 |
|  |  |  | lateral | 47.3 | 7.8 | 0.8 | 24.3 - 68.5 |

**Table S3.**
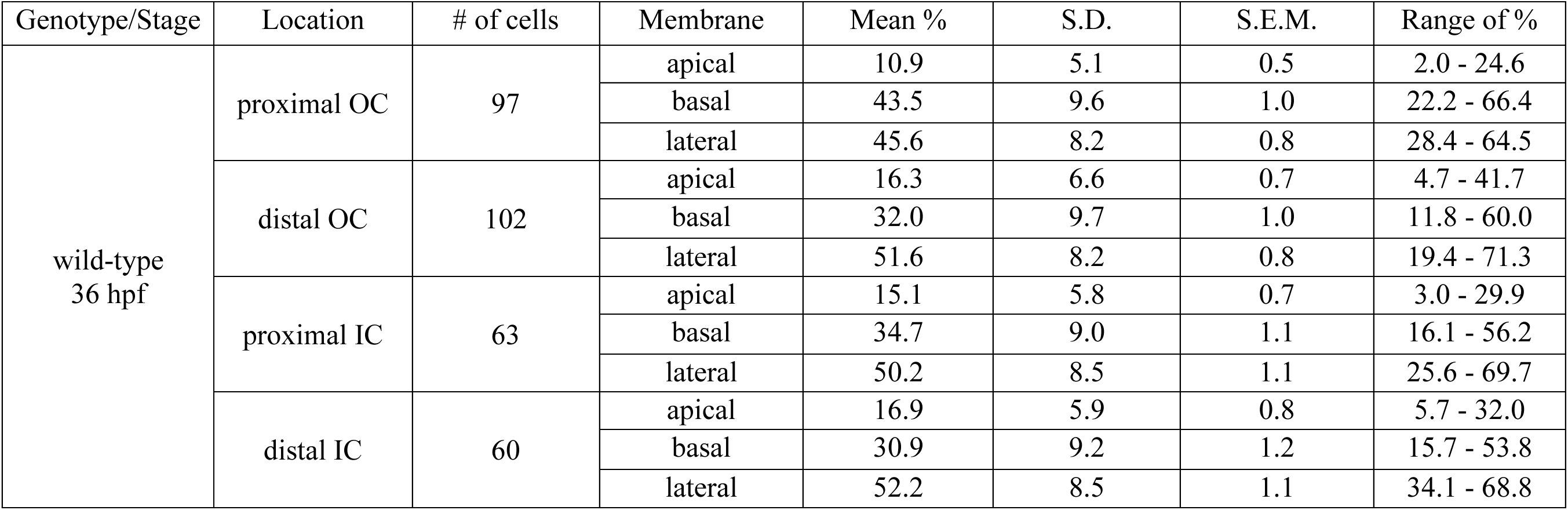
Summary statistics corresponding to F-actin localization in Figure 2—figure supplement 2D. Information is presented as in Table S1.

| Genotype/Stage | Location | # of cells | Membrane | Mean % | S.D. | S.E.M. | Range of % |
| --- | --- | --- | --- | --- | --- | --- | --- |
| wild-type<br>36 hpf | proximal OC | 97 | apical | 10.9 | 5.1 | 0.5 | 2.0 - 24.6 |
|  |  |  | basal | 43.5 | 9.6 | 1.0 | 22.2 - 66.4 |
|  |  |  | lateral | 45.6 | 8.2 | 0.8 | 28.4 - 64.5 |
|  | distal OC | 102 | apical | 16.3 | 6.6 | 0.7 | 4.7 - 41.7 |
|  |  |  | basal | 32.0 | 9.7 | 1.0 | 11.8 - 60.0 |
|  |  |  | lateral | 51.6 | 8.2 | 0.8 | 19.4 - 71.3 |
|  | proximal IC | 63 | apical | 15.1 | 5.8 | 0.7 | 3.0 - 29.9 |
|  |  |  | basal | 34.7 | 9.0 | 1.1 | 16.1 - 56.2 |
|  |  |  | lateral | 50.2 | 8.5 | 1.1 | 25.6 - 69.7 |
|  | distal IC | 60 | apical | 16.9 | 5.9 | 0.8 | 5.7 - 32.0 |
|  |  |  | basal | 30.9 | 9.2 | 1.2 | 15.7 - 53.8 |
|  |  |  | lateral | 52.2 | 8.5 | 1.1 | 34.1 - 68.8 |

**Table S4.** Summary statistics corresponding to pMyosin localization in Figure 2—figure supplement 2D. Information is presented as in Table S1.

| Genotype/Stage | Location | # of cells | Membrane | Mean % | S.D. | S.E.M. | Range of % |
| --- | --- | --- | --- | --- | --- | --- | --- |
| wild-type<br>36 hpf | proximal OC | 85 | apical | 14.1 | 5.3 | 0.6 | 4.8 - 28.7 |
|  |  |  | basal | 43.4 | 11.0 | 1.2 | 19.6 - 65.6 |
|  |  |  | lateral | 42.5 | 8.2 | 0.9 | 25.7 - 62.0 |
|  | distal OC | 89 | apical | 19.2 | 7.4 | 0.8 | 8.7 - 43.7 |
|  |  |  | basal | 31.6 | 9.9 | 1.0 | 10.2 - 52.6 |
|  |  |  | lateral | 49.2 | 7.6 | 0.8 | 23.3 - 67.8 |
|  | proximal IC | 56 | apical | 18.0 | 6.6 | 0.9 | 5.6 - 39.4 |
|  |  |  | basal | 36.0 | 10.6 | 1.4 | 17.2 - 59.5 |
|  |  |  | lateral | 46.0 | 7.8 | 1.0 | 24.3 - 61.4 |
|  | distal IC | 53 | apical | 18.8 | 7.0 | 1.0 | 8.1 - 33.4 |
|  |  |  | basal | 32.6 | 10.6 | 1.5 | 16.7 - 54.1 |
|  |  |  | lateral | 48.6 | 7.8 | 1.1 | 35.2 - 68.6 |

**Table S5.** Summary statistics corresponding to F-actin localization in Figure 3—figure supplement 3G. Information is presented as in Table S1.

| Location | Genotype/Stage | # of cells | Membrane | Mean % | S.D. | S.E.M. | Range of % |
| --- | --- | --- | --- | --- | --- | --- | --- |
| proximal OC | *WT myl9*<br>36 hpf | 39 | apical | 11.3 | 3.1 | 0.5 | 7.1 - 19.1 |
|  |  |  | basal | 44.4 | 10.6 | 1.7 | 22.9 - 62.2 |
|  |  |  | lateral | 44.4 | 8.7 | 1.4 | 28.3 - 63.3 |
|  | *DN myl9*<br>36 hpf | 45 | apical | 14.6 | 7.3 | 1.1 | 5.2 - 42.2 |
|  |  |  | basal | 35.7 | 12.3 | 1.8 | 14.8 - 68.8 |
|  |  |  | lateral | 49.7 | 12.0 | 1.8 | 15.4 - 80.0 |
|  | *CA myl9*<br>36 hpf | 43 | apical | 16.3 | 5.9 | 0.9 | 6.7 - 37.1 |
|  |  |  | basal | 37.6 | 8.3 | 1.3 | 24.2 - 55.9 |
|  |  |  | lateral | 46.1 | 7.5 | 1.1 | 32.8 - 63.2 |
| distal OC | *WT myl9*<br>36 hpf | 42 | apical | 15.1 | 5.0 | 0.8 | 7.6 - 25.7 |
|  |  |  | basal | 37.5 | 10.9 | 1.7 | 16.2 - 57.1 |
|  |  |  | lateral | 47.4 | 8.3 | 1.3 | 24.7 - 67.5 |
|  | *DN myl9*<br>36 hpf | 46 | apical | 16.1 | 4.7 | 0.7 | 6.8 - 26.7 |
|  |  |  | basal | 33.8 | 11.6 | 1.7 | 17.8 - 76.0 |
|  |  |  | lateral | 50.1 | 9.1 | 1.3 | 17.2 - 67.5 |
|  | *CA myl9*<br>36 hpf | 44 | apical | 21.9 | 7.4 | 1.1 | 8.4 - 41.7 |
|  |  |  | basal | 31.3 | 9.1 | 1.4 | 15.4 - 56.3 |
|  |  |  | lateral | 46.8 | 7.3 | 1.1 | 35.3 - 64.5 |
| IC | *WT myl9*<br>36 hpf | 50 | apical | 13.2 | 3.6 | 0.5 | 7.5 - 22.3 |
|  |  |  | basal | 37.8 | 10.0 | 1.4 | 16.5 - 55.8 |
|  |  |  | lateral | 48.9 | 7.9 | 1.1 | 34.9 - 65.4 |
|  | *DN myl9*<br>36 hpf | 53 | apical | 14.2 | 5.7 | 0.8 | 4.7 - 36.7 |
|  |  |  | basal | 35.8 | 9.1 | 1.3 | 17.4 - 53.7 |
|  |  |  | lateral | 50.0 | 7.8 | 1.1 | 36.5 - 67.6 |
|  | *CA myl9*<br>36 hpf | 55 | apical | 17.6 | 5.4 | 0.7 | 8.1 - 37.7 |
|  |  |  | basal | 27.6 | 8.4 | 1.1 | 12.8 - 48.4 |
|  |  |  | lateral | 54.8 | 7.0 | 0.9 | 38.0 - 68.5 |

**Table S6.** Summary statistics corresponding to F-actin localization in Figure 5—figure supplement 1E. Information is presented as in Table S1.

| Location | Genotype/Stage | # of cells | Membrane | Mean % | S.D. | S.E.M. | Range of % |
| --- | --- | --- | --- | --- | --- | --- | --- |
| proximal OC | wild-type<br>36 hpf | 39 | apical | 13.7 | 4.9 | 0.8 | 6.0 - 24.6 |
|  |  |  | basal | 42.8 | 9.5 | 1.5 | 22.5 - 63.8 |
|  |  |  | lateral | 43.6 | 8.1 | 1.3 | 28.4 - 61.6 |
|  | *myh6*<br>36 hpf | 32 | apical | 21.6 | 4.6 | 0.8 | 14.3 - 32.3 |
|  |  |  | basal | 28.1 | 8.3 | 1.5 | 12.1 - 42.7 |
|  |  |  | lateral | 50.3 | 7.6 | 1.3 | 38.1 - 69.0 |
| distal OC | wild-type<br>36 hpf | 41 | apical | 17.3 | 6.0 | 0.9 | 8.9 - 33.5 |
|  |  |  | basal | 33.2 | 9.7 | 1.5 | 11.8 - 52.7 |
|  |  |  | lateral | 49.5 | 8.8 | 1.4 | 19.4 - 66.0 |
|  | *myh6*<br>36 hpf | 24 | apical | 23.3 | 6.1 | 1.2 | 16.4 - 41.6 |
|  |  |  | basal | 25.9 | 5.6 | 1.1 | 16.6 - 35.4 |
|  |  |  | lateral | 50.8 | 5.8 | 1.2 | 39.5 - 61.9 |
| IC | wild-type<br>36 hpf | 54 | apical | 18.5 | 4.8 | 0.7 | 9.7 - 29.9 |
|  |  |  | basal | 32.7 | 8.6 | 1.2 | 17.5 - 54.3 |
|  |  |  | lateral | 48.7 | 7.4 | 1.0 | 26.6 - 62.5 |
|  | *myh6*<br>36 hpf | 35 | apical | 19.7 | 4.6 | 0.8 | 10.0 - 29.0 |
|  |  |  | basal | 32.9 | 10.4 | 1.8 | 10.5 - 54.7 |
|  |  |  | lateral | 47.5 | 7.6 | 1.3 | 31.8 - 65.9 |

**Table S7.** Summary statistics corresponding to pMyosin localization in Figure 5—figure supplement 1E. Information is presented as in Table S1.

| Location | Genotype/Stage | # of cells | Membrane | Mean % | S.D. | S.E.M. | Range of % |
| --- | --- | --- | --- | --- | --- | --- | --- |
| proximal OC | wild-type<br>36 hpf | 39 | apical | 13.8 | 5.9 | 0.9 | 4.8 - 28.1 |
|  |  |  | basal | 42.1 | 10.5 | 1.7 | 23.6 - 60.4 |
|  |  |  | lateral | 44.1 | 7.8 | 1.3 | 33.0 - 62.0 |
|  | *myh6*<br>36 hpf | 32 | apical | 22.2 | 6.2 | 1.1 | 9.9 - 35.1 |
|  |  |  | basal | 27.6 | 11.0 | 1.9 | 9.6 - 53.6 |
|  |  |  | lateral | 50.2 | 9.3 | 1.6 | 36.3 - 72.9 |
| distal OC | wild-type<br>36 hpf | 41 | apical | 19.2 | 7.6 | 1.2 | 8.7 - 36.9 |
|  |  |  | basal | 32.3 | 9.0 | 1.4 | 10.2 - 49.0 |
|  |  |  | lateral | 48.5 | 7.3 | 1.1 | 23.3 - 60.9 |
|  | *myh6*<br>36 hpf | 24 | apical | 25.2 | 7.0 | 1.4 | 15.6 - 41.1 |
|  |  |  | basal | 25.4 | 8.6 | 1.8 | 11.2 - 45.4 |
|  |  |  | lateral | 49.5 | 5.8 | 1.2 | 34.8 - 61.4 |
| IC | wild-type<br>36 hpf | 54 | apical | 18.2 | 6.9 | 0.9 | 9.1 - 39.4 |
|  |  |  | basal | 33.2 | 10.3 | 1.4 | 17.1 - 54.1 |
|  |  |  | lateral | 48.6 | 7.2 | 1.0 | 29.4 - 68.5 |
|  | *myh6*<br>36 hpf | 35 | apical | 17.3 | 4.9 | 0.8 | 6.3 - 28.0 |
|  |  |  | basal | 35.9 | 12.1 | 2.0 | 11.0 - 61.1 |
|  |  |  | lateral | 46.8 | 8.8 | 1.5 | 31.3 - 66.4 |

**Table S8.** Summary statistics corresponding to F-actin localization in Figure 6—figure supplement 1E. Information is presented as in Table S1.

| Location | Genotype/Stage | # of cells | Membrane | Mean % | S.D. | S.E.M. | Range of % |
| --- | --- | --- | --- | --- | --- | --- | --- |
| proximal OC | wild-type<br>36 hpf | 58 | apical | 9.0 | 4.4 | 0.6 | 2.0 - 22.3 |
|  |  |  | basal | 44.0 | 9.7 | 1.3 | 22.2 - 66.4 |
|  |  |  | lateral | 47.0 | 8.1 | 1.1 | 29.9 - 64.5 |
|  | *tbx5a*<br>36 hpf | 64 | apical | 14.0 | 8.2 | 1.0 | 3.6 - 42.3 |
|  |  |  | basal | 30.8 | 10.1 | 1.3 | 11.1 - 55.2 |
|  |  |  | lateral | 55.2 | 10.6 | 1.3 | 29.0 - 82.3 |
| distal OC | wild-type<br>36 hpf | 61 | apical | 15.7 | 7.0 | 0.9 | 4.7 - 41.7 |
|  |  |  | basal | 31.2 | 9.8 | 1.2 | 13.7 - 60.0 |
|  |  |  | lateral | 53.1 | 7.6 | 1.0 | 32.8 - 71.3 |
|  | *tbx5a*<br>36 hpf | 67 | apical | 20.4 | 10.1 | 1.2 | 5.7 - 55.2 |
|  |  |  | basal | 22.2 | 9.1 | 1.1 | 5.7 - 45.1 |
|  |  |  | lateral | 57.4 | 9.5 | 1.2 | 36.8 - 76.9 |
| IC | wild-type<br>36 hpf | 69 | apical | 14.0 | 5.9 | 0.7 | 3.0 - 32.0 |
|  |  |  | basal | 32.8 | 9.8 | 1.2 | 15.7 - 56.2 |
|  |  |  | lateral | 53.1 | 9.0 | 1.1 | 25.6 - 69.7 |
|  | *tbx5a*<br>36 hpf | 81 | apical | 15.6 | 9.4 | 1.1 | 3.2 - 38.9 |
|  |  |  | basal | 25.9 | 10.3 | 1.2 | 5.9 - 50.9 |
|  |  |  | lateral | 58.6 | 9.3 | 1.0 | 41.9 - 81.0 |

**Table S9.** Summary statistics corresponding to pMyosin localization in Figure 6—figure supplement 1E. Information is presented as in Table S1.

| Location | Genotype/Stage | # of cells | Membrane | Mean % | S.D. | S.E.M. | Range of % |
| --- | --- | --- | --- | --- | --- | --- | --- |
| proximal OC | wild-type<br>36 hpf | 46 | apical | 14.4 | 4.7 | 0.7 | 8.3 - 28.7 |
|  |  |  | basal | 44.5 | 11.4 | 1.7 | 19.6 - 65.6 |
|  |  |  | lateral | 41.1 | 8.4 | 1.2 | 25.7 - 60.4 |
|  | *tbx5a*<br>36 hpf | 64 | apical | 22.5 | 9.3 | 1.2 | 8.0 - 49.2 |
|  |  |  | basal | 38.9 | 10.9 | 1.4 | 16.6 - 64.4 |
|  |  |  | lateral | 38.6 | 7.4 | 0.9 | 20.1 - 59.9 |
| distal OC | wild-type<br>36 hpf | 48 | apical | 19.2 | 7.2 | 1.0 | 9.4 - 43.7 |
|  |  |  | basal | 30.9 | 10.6 | 1.5 | 14.4 - 52.6 |
|  |  |  | lateral | 49.9 | 7.7 | 1.1 | 35.5 - 67.8 |
|  | *tbx5a*<br>36 hpf | 67 | apical | 26.5 | 11.0 | 1.4 | 9.0 - 54.6 |
|  |  |  | basal | 30.5 | 11.1 | 1.4 | 8.4 - 53.1 |
|  |  |  | lateral | 43.1 | 7.6 | 0.9 | 24.4 - 64.7 |
| IC | wild-type<br>36 hpf | 55 | apical | 18.5 | 6.7 | 0.9 | 5.6 - 33.4 |
|  |  |  | basal | 35.5 | 11.1 | 1.5 | 16.7 - 59.5 |
|  |  |  | lateral | 46.0 | 8.3 | 1.1 | 24.3 - 66.0 |
|  | *tbx5a*<br>36 hpf | 81 | apical | 22.6 | 11.0 | 1.2 | 7.5 - 49.5 |
|  |  |  | basal | 35.8 | 11.6 | 1.3 | 10.9 - 57.8 |
|  |  |  | lateral | 41.6 | 7.2 | 0.8 | 27.3 - 58.5 |

**Table S10.** Summary statistics corresponding to F-actin localization in Figure 7—figure supplement 1A. Information is presented as in Table S1.

| Location | Type of cell/Stage | # of cells | Membrane | Mean % | S.D. | S.E.M. | Range of % |
| --- | --- | --- | --- | --- | --- | --- | --- |
| proximal OC | wild-type host<br>36 hpf | 55 | apical | 14.2 | 5.3 | 0.7 | 4.2 - 28.8 |
|  |  |  | basal | 41.4 | 10.8 | 1.5 | 21.9 - 63.9 |
|  |  |  | lateral | 44.3 | 7.5 | 1.0 | 30.9 - 63.2 |
|  | wild-type donor<br>36 hpf | 14 | apical | 14.2 | 5.0 | 1.3 | 7.2 - 25.6 |
|  |  |  | basal | 38.9 | 8.0 | 2.1 | 24.6 - 50.5 |
|  |  |  | lateral | 46.9 | 7.0 | 1.9 | 38.9 - 62.9 |
| distal OC | wild-type host<br>36 hpf | 30 | apical | 16.3 | 4.5 | 0.8 | 6.5 - 24.0 |
|  |  |  | basal | 35.8 | 7.5 | 1.4 | 19.2 - 46.4 |
|  |  |  | lateral | 47.9 | 8.1 | 1.5 | 31.0 - 66.6 |
|  | wild-type donor<br>36 hpf | 29 | apical | 18.6 | 6.5 | 1.2 | 11.1 - 40.0 |
|  |  |  | basal | 34.7 | 10.5 | 1.9 | 13.9 - 50.7 |
|  |  |  | lateral | 46.7 | 6.5 | 1.2 | 34.3 - 58.8 |
| IC | wild-type host<br>36 hpf | 30 | apical | 19.1 | 5.5 | 1.0 | 7.0 - 31.5 |
|  |  |  | basal | 30.0 | 11.1 | 2.0 | 10.1 - 51.5 |
|  |  |  | lateral | 50.8 | 9.2 | 1.7 | 34.6 - 69.6 |
|  | wild-type donor<br>36 hpf | 29 | apical | 19.1 | 4.0 | 0.7 | 12.2 - 27.6 |
|  |  |  | basal | 29.4 | 8.7 | 1.6 | 15.5 - 45.0 |
|  |  |  | lateral | 51.5 | 8.0 | 1.5 | 34.3 - 64.2 |

**Table S11.** Summary statistics corresponding to F-actin localization in Figure 7—figure supplement 1B. Information is presented as in Table S1.

| Location | Type of cell/Stage | # of cells | Membrane | Mean % | S.D. | S.E.M. | Range of % |
| --- | --- | --- | --- | --- | --- | --- | --- |
| proximal OC | *tbx5a* host<br>36 hpf | 40 | apical | 21.7 | 10.5 | 1.7 | 3.5 - 51.1 |
|  |  |  | basal | 31.0 | 12.0 | 1.9 | 8.8 - 60.1 |
|  |  |  | lateral | 47.3 | 7.3 | 1.2 | 32.4 - 61.3 |
|  | wild-type donor<br>36 hpf | 25 | apical | 17.2 | 6.5 | 1.3 | 7.6 - 33.8 |
|  |  |  | basal | 39.4 | 11.7 | 2.3 | 15.4 - 59.0 |
|  |  |  | lateral | 43.3 | 6.8 | 1.4 | 30.2 - 55.4 |
| distal OC | *tbx5a* host<br>36 hpf | 40 | apical | 26.5 | 10.3 | 1.6 | 9.7 - 53.4 |
|  |  |  | basal | 27.4 | 8.6 | 1.4 | 11.8 - 45.5 |
|  |  |  | lateral | 46.1 | 8.6 | 1.4 | 25.8 - 62.8 |
|  | wild-type donor<br>36 hpf | 35 | apical | 21.7 | 7.7 | 1.3 | 5.7 - 39.7 |
|  |  |  | basal | 28.7 | 7.6 | 1.3 | 18.4 - 49.1 |
|  |  |  | lateral | 49.6 | 9.4 | 1.6 | 23.1 - 67.7 |
| IC | *tbx5a* host<br>36 hpf | 18 | apical | 22.8 | 6.0 | 1.4 | 12.2 - 36.9 |
|  |  |  | basal | 29.4 | 8.2 | 1.9 | 15.2 - 41.4 |
|  |  |  | lateral | 47.8 | 9.4 | 2.2 | 32.6 - 72.6 |
|  | wild-type donor<br>36 hpf | 14 | apical | 19.5 | 4.7 | 1.3 | 10.9 - 29.7 |
|  |  |  | basal | 27.9 | 6.5 | 1.7 | 18.1 - 44.4 |
|  |  |  | lateral | 52.6 | 6.3 | 1.7 | 41.2 - 66.8 |

**Table S12.** Summary statistics corresponding to F-actin localization in Figure 7—figure supplement 1C. Information is presented as in Table S1.

| Location | Type of cell/Stage | # of cells | Membrane | Mean % | S.D. | S.E.M. | Range of % |
| --- | --- | --- | --- | --- | --- | --- | --- |
| proximal OC | wild-type host<br>36 hpf | 38 | apical | 14.0 | 5.7 | 0.9 | 5.0 - 27.3 |
|  |  |  | basal | 39.3 | 10.3 | 1.7 | 12.6 - 58.6 |
|  |  |  | lateral | 46.7 | 7.7 | 1.2 | 28.4 - 61.6 |
|  | *tbx5a* donor<br>36 hpf | 14 | apical | 18.6 | 10.5 | 2.8 | 5.4 - 41.1 |
|  |  |  | basal | 35.1 | 11.2 | 3.0 | 16.9 - 59.3 |
|  |  |  | lateral | 46.3 | 7.5 | 2.0 | 33.1 - 58.6 |
| distal OC | wild-type host<br>36 hpf | 28 | apical | 18.5 | 8.6 | 1.6 | 6.8 - 43.5 |
|  |  |  | basal | 29.9 | 10.2 | 1.9 | 10.3 - 45.7 |
|  |  |  | lateral | 51.6 | 6.0 | 1.1 | 43.2 - 65.1 |
|  | *tbx5a* donor<br>36 hpf | 13 | apical | 30.0 | 10.0 | 2.8 | 8.9 - 43.8 |
|  |  |  | basal | 26.0 | 6.7 | 1.9 | 15.2 - 33.6 |
|  |  |  | lateral | 44.0 | 8.9 | 2.5 | 32.5 - 61.9 |
| IC | wild-type host<br>36 hpf | 44 | apical | 17.3 | 6.3 | 0.9 | 8.6 - 37.7 |
|  |  |  | basal | 30.7 | 12.0 | 1.8 | 12.1 - 56.7 |
|  |  |  | lateral | 52.0 | 9.8 | 1.5 | 31.1 - 73.3 |
|  | *tbx5a* donor<br>36 hpf | 18 | apical | 23.0 | 7.6 | 1.8 | 9.3 - 39.1 |
|  |  |  | basal | 23.1 | 9.4 | 2.2 | 9.8 - 41.0 |
|  |  |  | lateral | 53.9 | 8.0 | 1.9 | 39.3 - 68.2 |

**Table S13.**
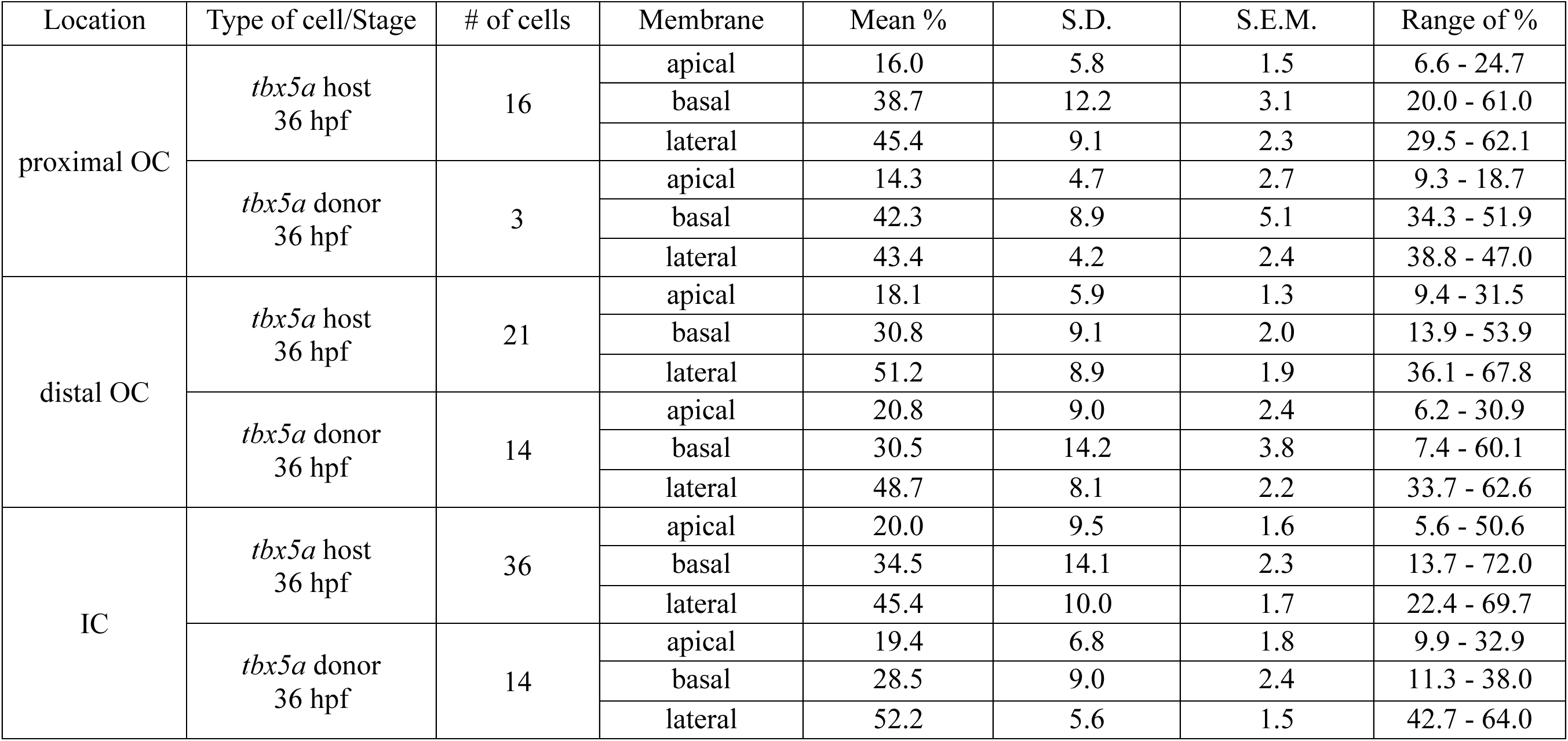
Summary statistics corresponding to F-actin localization in Figure 7—figure supplement 1D. Information is presented as in Table S1.

